# Precision Targeting of Heparan Sulfate Proteoglycans Broadly Dysregulates Survival Signaling and Metabolic Adaptation in Pancreatic Cancer

**DOI:** 10.64898/2026.10.01.756059

**Authors:** Nancy D. Ebelt, Suvithanandhini Loganathan, Lara C. Avsharian, Katarzyna Dabrowska, Nathaniel Hansen, Khyatiben V. Pathak, Patrick Pirrotte, Edwin R. Manuel

## Abstract

Treatments for pancreatic ductal adenocarcinoma (PDAC) often fail because of the cancer’s ability to adapt its signaling and metabolic pathways to circumvent treatment mechanisms. PDAC hallmarks that promote treatment resistance (increased fatty acid metabolism, survival signaling, desmoplasia, and macropinocytosis) are extensively regulated by heparan sulfate proteoglycans (HSPGs) and their binding partners. The pro-tumor functions of HSPGs are controlled by site-specific cleavage of their heparan sulfate chains by mammalian heparanase; however, cleavage by bacterial heparinase III (HepIII) represses these functions and prevents downstream, pro-tumor signaling. Using tumor-targeting, attenuated *Salmonella typhimurium* (strain YS1646), we have limited HepIII transgene expression to tumor tissue. Degradation of tumor cell heparan sulfates by HepIII resulted in decreased macropinocytosis, reduced survival signaling, lipid storage depletion, and decreased expression of translational machinery. This tumor-targeting treatment has the potential to improve the efficacy of several approved pancreatic cancer therapies, which is predicted to increase patient survival.

## Introduction

Resistance to targeted cancer therapies is frequently developed due to the ease in which tumor cells are able to adapt their metabolic processes and compensate for impaired survival signaling with pathway redundancy(*1, 2*). While combination therapies exist to target mechanisms of resistance and are in use for many cancer types(*3–5*), resistance still occurs(*6*), often involving metabolic circumvention(*6, 7*). Unfortunately, adding supplementary targeted therapies to overcome acquired resistance to combination treatment often results in additive or synergistic toxicities in healthy tissues that utilize the same pathways for normal function(*8–11*). Strategic combinations of tumor-targeting delivery methods with polypharmacologic treatments may overcome these issues. Pancreatic ductal adenocarcinoma (PDAC) is set to become the second leading cause of cancer deaths in the USA by 2030(*12*) due, in part, to inadequate early detection methods(*13*), but also due to rapid resistance that is acquired after chemotherapeutic treatment(*14*), which remains the only effective therapy(*15*). Treatments that target multiple hubs of PDAC survival will be needed to overcome chemo-resistance mechanisms and improve patient survival.

Due to the preponderance of mutations in *KRAS*, *TP53*, *SMAD4*, and *CDKN2A*(*16*), PDAC has particular dependences on certain signaling and metabolic pathways. Major pathways activated downstream of mutant KRAS include MAPK signaling, PI3K/Akt/mTOR signaling, and RalGEF signaling(*17, 18*), while inhibition of the tumor suppressor genes (*TP53, SMAD4, CDKN2A*) allows for proliferation without DNA damage checkpoints(*19, 20*). Sustained proliferation and KRAS signaling leads to overexpression of growth factor receptors to increase survival signaling, and increased signaling through Wnt/β-catenin, NOTCH, and TGF-β to maintain stemness, increase proliferation and survival, and promote epithelial-to-mesenchymal transition (EMT)(*21–23*). Within metabolic rewiring, PDAC relies heavily upon fatty acid oxidation for energy and shows upregulation of various enzymes involved in *de novo* fatty acid synthesis(*24–26*). Additionally, rewiring of glucose metabolism in PDAC includes increased expression of glucose transporter 1 and other glycolytic enzymes to promote glycolysis, but glucose carbons are frequently shuttled toward anabolic pathways to increase synthesis of macromolecules such as amino acids, lipids, and nucleotides(*27*). Altered amino acid metabolism includes increased amino acid transporters at the cell surface, increased proline oxidase, asparagine depletion, and a dependency on glutamine metabolism(*28–32*). To further ease the stress of sustained proliferative growth, cancer cells supplement their anabolic processes with scavenged end-products and intermediates as well as fatty acids for oxidation with autophagy and macropinocytosis of the tumor microenvironment (TME)(*33–35*). Dependency on fatty acid oxidation, glucose-driven anabolic pathways, EGFR signaling, and macropinocytosis represent targetable vulnerabilities for PDAC treatment(*33, 34, 36–42*).

Molecules at the crux of many of these processes are heparan sulfate proteoglycans (HSPG) whose activity is largely regulated by interactions between, and cleavage of, their heparan sulfate (HS) chains(*43–45*). Mammalian heparanase (HSPE) regulates specific cleavage of HS, a function that is appropriated by many cancer types to drive tumor progression(*46–48*). HPSE facilitates tumor growth through a number of HSPG. Growth factor and morphogen signaling, notably through FGFR, VEGFR, and Wnt, are heavily influenced by HSPE wherein HS bound ligands and receptors can be activated and mobilized through specific cleavage of their associated HS chains(*49*), and other HSPG such as Syndecans (SDC) 1-4 can act as coreceptors to facilitate signaling(*50, 51*). Migration, metastasis, and other ECM-regulated events can be modulated by HSPE through the interactions of syndecans with integrins(*52*), through cleavage of HS on Collagen XVIII, through the HSPG Perlecan as a co-receptor for Hedgehog signaling(*53*), and through interactions of glypicans (another class of HSPG) with matrix metalloproteinases, laminins, fibronectin and collagen(*54*). Cytokine reserves are also held in the ECM by HSPG(*55*) similar to growth factors and other ligands. Additionally, HPSE cleavage of HS chains found on SDC1 results in its recycling and redistribution on the surface of cells to regulate macropinocytosis(*56–60*).

As a prominent facilitator of tumor growth and metastasis, inhibition of HSPE is an ongoing and active field of research with various classes of inhibitors under investigation(*61–65*). However, as with most targets, lack of tumor-specific inhibition poses a major risk due to the broad activity of HSPE in healthy tissue that regulates anti-tumor mechanisms(*66*). Interestingly, bacterial lyases, such as HepIII, also degrade HS, but at alternate sites and with different outcomes. HSPE cleaves HS chains at two specific linkages (out of a possible 32) resulting in distinct and typically larger fragments(*67, 68*) that promote cancer progression, therapeutic resistance, and worse prognosis in patients(*48, 69–71*). In contrast, HepIII cleaves HS at additional unique linkages resulting in degradation into smaller fragments that suppress neovascularization, tumor growth and metastasis(*72–75*). As a therapeutic approach to simultaneously target several major nodes of PDAC survival, we have engineered tumor-colonizing, attenuated *Salmonella typhimurium* strain YS1646 (YS) expressing functional bacterial HepIII (YS-HepIII), which degrades heparan sulfates attached to HSPGs, causing their inactivation. This method circumvents the issues of off-target effects from HSPE inhibition or HS degradation in healthy tissue due to restriction of HepIII action to tumor tissue via the YS delivery vector(*76, 77*). We hypothesize that targeting PDAC tumors with YS-HepIII will dysregulate multiple nodes of PDAC survival, leading to increased sensitivity to approved chemotherapeutics, like gemcitabine(*78*), or previously ineffective metabolic interventions such as fatty acid synthase inhibitors and mTOR inhibitors, whose mechanisms of action are circumvented by macropinocytosis(*79, 80*)

## Results

### Bacterial HepIII Expressed from Recombinant YS1646 (YS-HepIII) Results in Specific Degradation of Heparan Sulfates

To minimize toxicity to the YS delivery vector and to allow for regulated, inducible expression of HepIII, we cloned individual HepIII sequences from various bacterial species downstream of the pBAD promoter taken from the arabinose-inducible operon of *E. coli*(*81*). At the N-terminus of the HepIII sequence is an export signal sequence (Fla) from the flagellin protein (FliC), which should localize HepIII protein to the outer membrane(*82*), and the C-terminus of the HepIII sequence was fused to a 6x His-tag (**Figure 1A**). This system allows for precise tuning of transgene expression, localization of the functional enzyme on the YS surface, and easy identification of the enzyme through the His-tag. Titration of L-arabinose at low levels allows for pronounced expression of the transgene (EGFP or HepIII) compared to cultures that are uninduced (0.000% L-Ara) (**Figure 1B**). Three HepIII sequences from varying bacterial species were cloned into the pBAD plasmid to determine differences in their HS degradation efficiencies (**Supp Figure 1A-C**), and degradation by each clone was measured *in vitro* by comparing the shift in size of purified HS that was co-incubated with the HepIII-expressing YS cultures. *In vitro* HS degradation shows that all three HepIII transgenes cloned into YS show some amount of HS degradation compared to parental YS, with the HepIII from *Bacteroides eggerthii* showing the most complete degradation (gel shift), even compared to commercially available, purified HepIII enzyme from *F. heparinum* (**Figure 1C**). All future experiments utilize YS expressing HepIII from *Bacteroides eggerthii*, from here on refferred to as YS-HepIII. Titration of L-arabinose reveals some leakiness in the pBAD promoter wherein there is comparable enzymatic activity without L-arabinose addition (**Supp Figure 2A**), possibly explaining the similar abundance of dead or dying bacteria in uninduced versus induced cultures (**Supp Figure 2B**). Lack of glucose (a pBAD repressor) in our *in vitro* system could account for pBAD promoter leakiness, so we titrated L-ara in the presence of varying glucose concentrations and observed HS degradation by YS-HepIII. In cultures supplemented with high glucose similar to what is found in the bloodstream (1mg/ml)(*83*), HS degradation is inhibited in non-L-arabinose-induced cultures, and only minimal HepIII activity can be detected with high concentrations of L-arabinose. In lower concentrations of glucose, similar to what might be found in tumor tissue (0.02mg/ml), HepIII enzymatic activity is no longer inhibited (**Supp Figure 2C**). These results indicate that *in vivo*, HepIII expression and activity should be restricted until YS-HepIII reach the tumor compartment. All future experiments include YS-HepIII induced at 0.01% L-arabinose without glucose unless otherwise noted.

**Figure 1.**
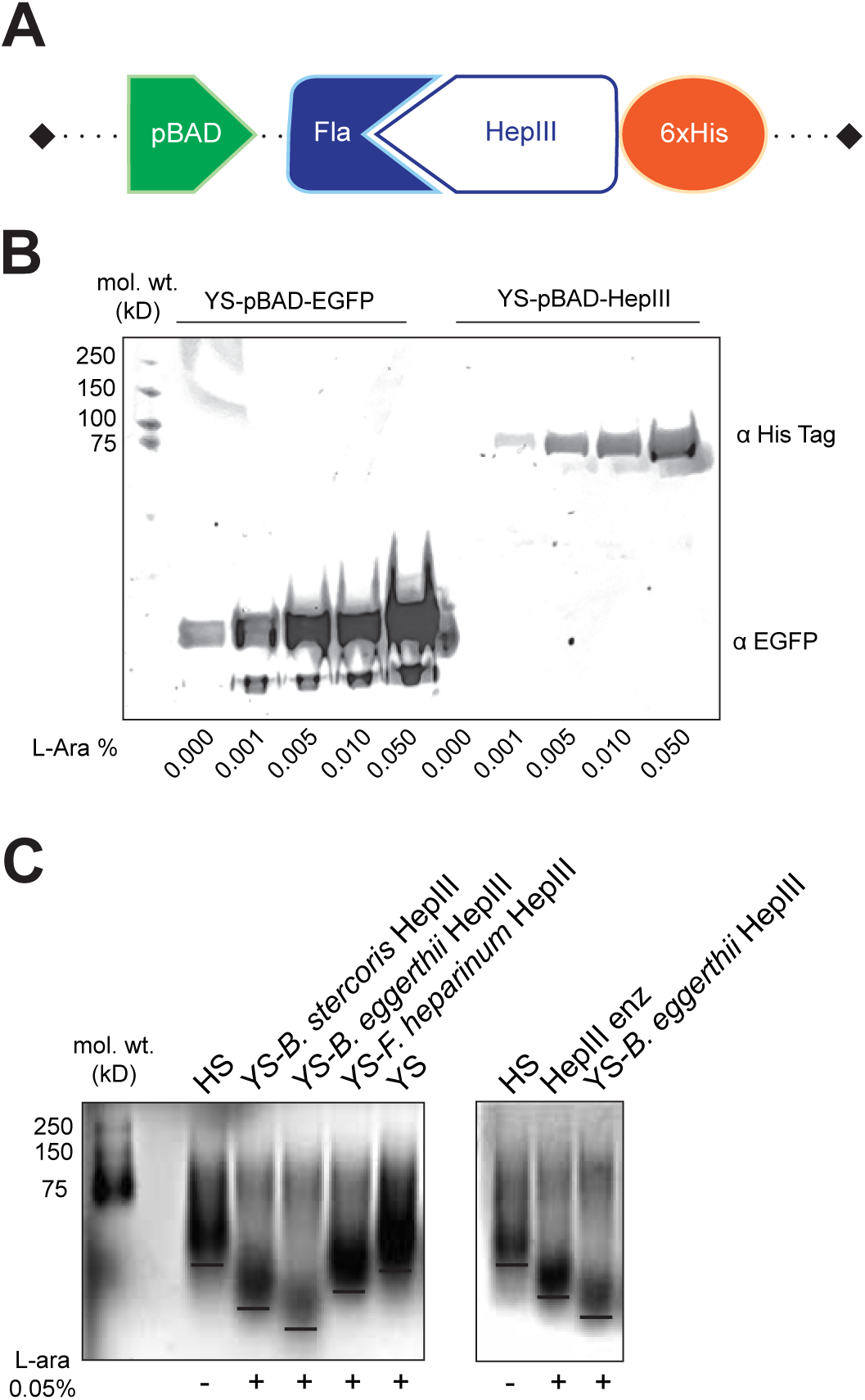
Transgenic Heparinase III (HepIII) Expressed from *Salmonella typhimurium* strain YS1646 (YS) Efficiently Degrades Heparan Sulfate (HS) *in vitro*. **(A)** Pictogram of HepIII cloning strategy showing HepIII attached to the N-terminal flagellin signal sequence (Fla) and a C-terminal 6xHis tag sequence all downstream of the L-arabinose inducible pBAD promoter. **(B)** ST electroporated with pBAD-EGFP or pBAD-HepIII were induced in culture with varying concentrations of L-arabinose (L-ara) for 45 minutes. Western blots of lysates were probed for the 6x His tag and EGFP. Molecular weight ladders in kD are labeled (mol. wt.) **(C)** YS-HepIII containing HepIII sequences from various bacterial species and parental YS were co incubated with purified HS after induction. After overnight degradation, supernatants were electrophoresed on acrylamide gels to visualize HS degradation compared to undigested HS and purified HepIII enzyme from *F. heparinus.* Gels were stained with Alcian blue to visualize HS smears.

### YS-HepIII Digests HS on the Surface of PDAC Cell Lines

To determine whether YS expressing the high functioning HepIII enzyme from *B. eggerthii* can degrade HS on the surfaces of pancreatic cancer cells, YS expressing either HepIII or EGFP were co-incubated with various PDAC cell lines (**Figure 2A**). Purified HepIII enzyme from *F. heparinum* (commercially available) was used to compare HepIII specific changes, and untreated cells (no YS) show base levels of HS staining on cell surfaces in full media. YS-HepIII treatment resulted in markedly decreased HS staining on the cell surfaces in most cell lines, after immunofluorescence with an antibody detecting HS, compared to untreated cells or cells co-incubated with YS-EGFP (**Figure 2B-E**). Pancreatic cancer cells treated with YS-HepIII showed significant depletion of HS, similar to that observed with purified HepIII enzyme. These results indicate that HS degradation achieved by co-incubation with YS-HepIII was due to specific expression of recombinant HepIII, and not due to endogenous HS lyase activity found in parental YS. Degredation of HS on the HSPG SDC1 was explored by western blot, wherein HepIII purified enzyme or YS-HepIII treatment is shown to shift SDC1 into a uniform size after degradation of variable length HS chains (approximately 90kD). PANC1 cells, which showed no observable degradation of surface HS by immunofluorescence, also showed negligible levels of SDC1 (**Figure 2D**). Altogether, these data indicate that SDC1 is a major source of surface bound HS in PDAC cells. Indeed, high SDC1 protein expression is found in the majority of human PDAC samples(*84, 85*), and high mRNA expression of SDC1 and 4 significantly correlates with poor overall survival in patients with PDAC (**Figure 2F**).

**Figure 2.**
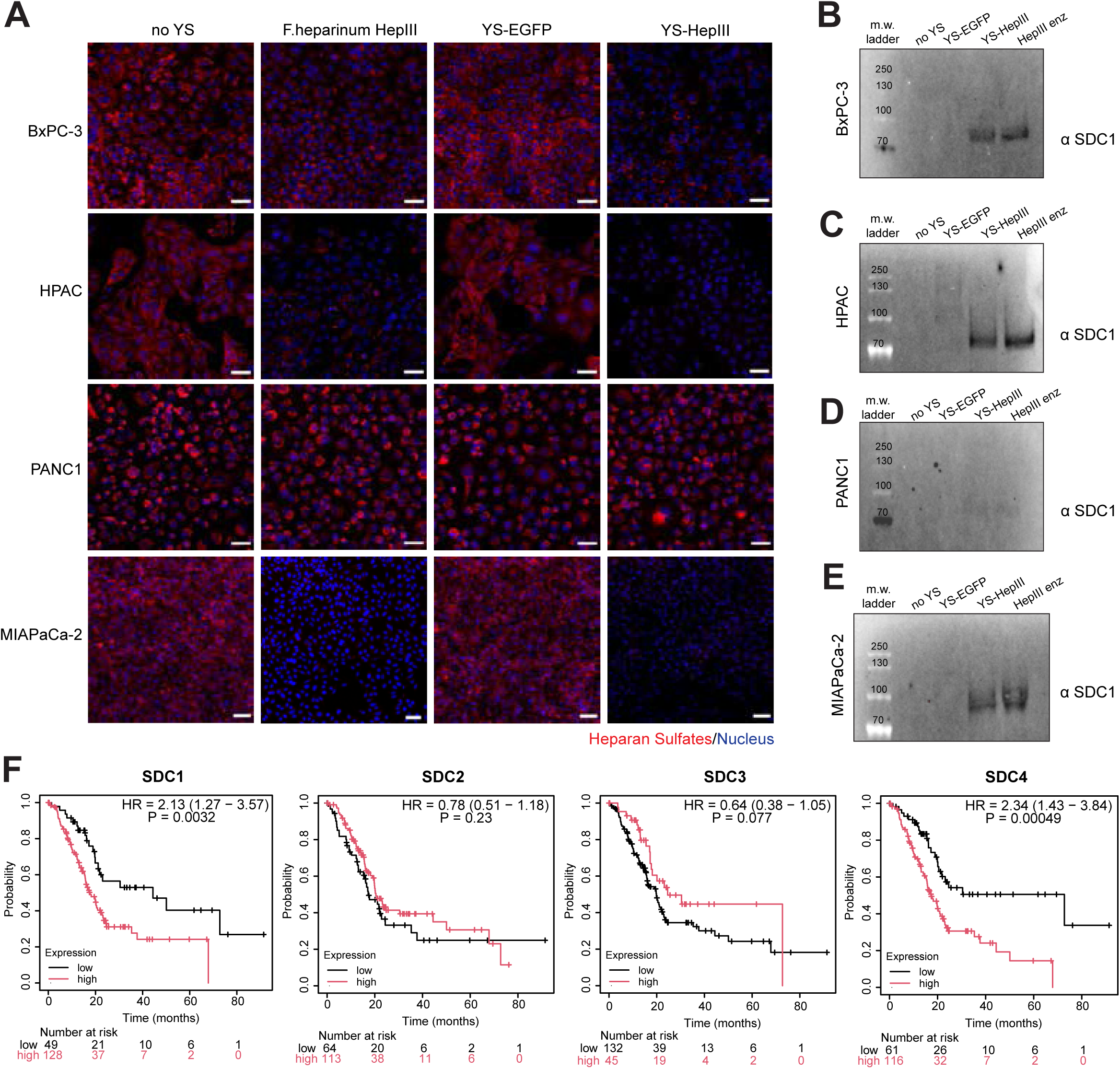
YS1646 expressing Heparinase III (YS-HepIII) degrade heparan sulfates (HS) on human pancreatic cancer cell lines in 2D culture. BxPC-3, HPAC, PANC1, and MIAPaCa-2 cell lines were plated to 80-90% confluency in full media overnight. The next day, the cells were left untreated (no YS), treated with purified HepIII enzyme from *F. heparinus* (5mU/mL), induced YS-EGFP, or induced YS-HepIII for three hours. After co-incubation, cells were washed thoroughly, and **(A)** fixed for immunofluorescence using an anti-HS antibody (red) with DAPI counterstain for nuclei (blue), or **(B-E)** cells were lysed and probed for Syndecan-1 (SDC1) staining by western blot. Representative fields are shown for immunofluorescence (A), and blots visualized by secondary antibody fluorescence are shown **(B-E)**. Molecular weight (m.w.) band sizes are labeled in kD. **(F)** Kaplan-Meier plots of high (red) or low (black) Syndecan 1-4 mRNA expression compared to overall survival probability (months) for human patients with PDAC (n=177). Individuals still in the study at the labeled timepoints are described in the “number at risk” table below each graph. Hazard ratios (HR) and their 95% confidence intervals are reported within each graph as well as the p-values from log-rank tests.

### YS-HepIII Treatment Induces Starvation Signaling Pathways in PDAC Tumors

HSPGs on the cell surface regulate numerous intracellular signaling pathways and are assumed to experience degradation of their associated HS chains similar to SDC1 (**Figure 2B-E**). We sought next to explore the effects of YS-HepIII treatment on various signaling networks including those downstream of HSPGs and associated co-receptors (**Figure 3A**). To do so, we first subjected HPAC cells in culture to starvation conditions (amino acid-low and serum-free), co-incubated with no YS, EGFP- or HepIII-expressing YS, and then stimulated signaling by addition of complete media. Using a phospho-proteomic microarray, we interrogated numerous phospho-sites on various signaling proteins to confirm whether signaling downstream of HSPGs is affected by YS-HepIII treatment. As expected, addition of YS-EGFP alone dysregulates various signaling pathways, as tumor cells in culture are affected by the inherent oncolytic capabilities of YS(*86*), and they must also compete with replicating YS (especially those that are induced to produce transgene proteins such as EGFP or HepIII) for available nutrients(*87, 88*). To determine the effects of YS alone, we pursued targets wherein the addition of HepIII was not significantly additive to the YS effect (**Figure 3B**). In general, phosphorylation states of signaling proteins in cells treated with YS (EGFP or HepIII) resembled those of the cells in starvation, despite stimulation with compete media. Exceptions include ICK (Y156/Y157), JAK2 (Y1007/Y1008), LATS1 (S464), MAPK13 (Y182), and PKG (T515/T517), which vary in their facilitation or inhibition of tumor growth (**Figure 3B**). The next set of targets represent those where HepIII is additive or synergistic to the effects of YS alone (**Figure 3C**). Again, many targets are returned to a status similar to that of when cells were in starvation after incubation with YS-HepIII prior to complete media stimulation, however, a majority of targeted phospho-sites are increased or decreased beyond levels seen in starving cells e.g. CSF1R (Y809), ErbB3 (Y1328), MAP3K7 (S439 and T184/T187), MERTK (Y749), Met (Y1234/Y1235), PECAM-1 (Y713), and Raf1 (S301/T303) (**Figure 3C**). Finally, we found targets that were unique to YS-HepIII treatment, in that compared to YS-HepIII, untreated (noYS) and YS-EGFP-treated cells equally differed in measured phosphorylation (**Figure 3D**). In this category, most changes attributable to YS-HepIII treatment differed greatly from phosphorylation levels seen in starving, unstimulated cells. Notably, activating phosphorylations Y656/Y657 on FGFR2, a receptor that binds SDC1 and whose ligand binds other HSPGs, are decreased 4-fold in YS-HepIII treated, stimulated cells compared to untreated, stimulated cells. Conversely, activating phosphorylations Y1289 and Y1307 on Erbb3 are nearly 3-fold upregulated by YS-HepIII treatment, possibly representing a resistance mechanism. Other notable targets include pro-survival kinases ERK3, ICK, MSK1, and Raf1 whose activating phosphorylations (S189, Y159, S212, and S296, respectively) are significantly inhibited after YS-HepIII treatment (**Figure 3D**). To determine the overall effect of YS-HepIII treatment on post-stimulation signaling, we compared fold changes for all targets between the YS-HepIII stimulated group and the untreated stimulated group. With this data, a kinase-substrate enrichment analysis was performed to infer significant changes in kinase activity not directly assayed using the phospho-protein array (**Figure 3E**). Kinases with significantly downregulated activity after YS-HepIII treatment include the receptors IGF1R, MerTK, and FGFR1, as well as the intracellular signaling kinases ERK3, PAK2, and ERK2. Kinases with increased activity include the TrkB and EphA2 receptors, and the intracellular signaling kinases LYN, CAMKK1 and 2, FGR, CDK20, MEK5, CAMK2A, and NuaK1 and 2 (**Figure 3E**). Overall, these results confirm that YS-HepIII treatment decreased phosphorylation of known HSPG and co-receptors, (e.g. FGFR1 and 2, EGFR and family members, VEGFR1, IGF1R) as well as for their downstream signaling pathways. However, increased activity in some receptors and intracellular signaling kinases could indicate early changes that promote resistance to the therapeutic actions of YS-HepIII (e.g. Erbb3, Met, CSF1R, ALK, and NuaK1 and 2).

**Figure 3.**
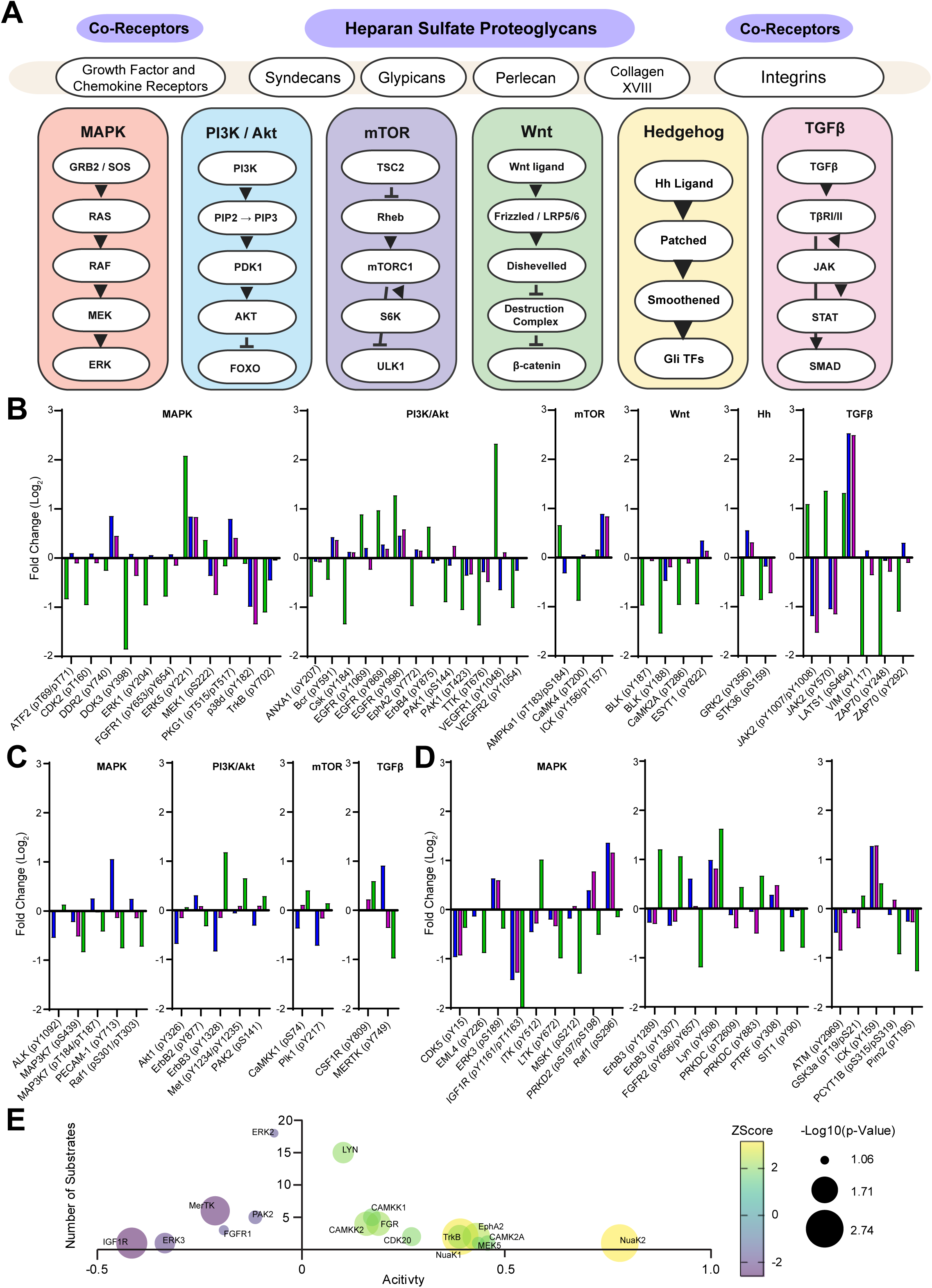
Degradation of heparan sulfate on HPAC cells in culture disrupts phosphorylation of intracellular signaling proteins. (**A**) Pictogram of intracellular signaling pathways (Mitogen Activate Protein Kinase (MAPK), Phosphatidylinositol 3-kinase/Akt (PI3K/Akt), mammalian Target of Rapamycin (mTOR), Wnt, Hedgehog (Hh), and Transforming Growth Factor β (TGFβ)) and individual proteins that are activated ( ) or inhibited ( ) downstream of heparan sulfate proteoglycans and their co-receptors. (**B-D**) HPAC cells in culture were starved overnight in minimal media (serum-free, low amino-acids, low glucose) prior to 3-hour treatment with PBS (no **YS**), YS-EGFP, or YS-HepIII. After treatment, cells were thoroughly washed and left unstimulated or stimulated for 15 minutes with complete media (DMEM with 10% FBS and 2mM L-Glutamine). All groups were lysed and phospho-protein levels were analyzed by microarray. Results are presented as stimulated no YS, YS-EGFP, or YS-HepIII groups compared to no YS, unstimulated cells (baseline, 0). Targets (protein name and phospho-site in parentheses) are loosely organized by pathway for ease of the viewer. (**B**) Targets that are significantly affected by treatment with YS alone (YS-EGFP and YS-HepIII show equal effect compared to no YS). (**C**) Targets that are significantly affected by treatment with both YS-EGFP and YS-HepIII (YS-HepIII effect is synergistic to YS-EGFP effect compared to no YS). (**D**) Targets that are significantly, uniquely affected by treatment with YS-HepIII (YS-EGFP and no YS show equal effect compared to YS-HepIII). (**E**) Phopsho-site fold changes were used to analyze kinase activity differences between no YS and YS-HepIII treated cells using the Robust Inference of Kinase Activity (ROKAI) algorithm. Change in kinase activity in YS-HepIII treated cells compared to no YS-treated cells is represented by a bubble plot where activity change is located on the axis, Z-score is represented by a change in bubble color, the number of substrates assayed for each kinase is on the Y-axis, and the size of the bubble corresponds to the -Log_10_ adjusted p-value.

### Targeting HS using YS-HepIII Decreases Macropinocytosis and Migration, but not Viability

The digestion of HS by YS-HepIII (**Figure 2**) resulting in repression of several downstream MAPK signaling pathways (**Figure 3**) may also decrease macropinocytosis, which is highly regulated by HS and MAPK signaling. *In vitro* assays have shown that cellular uptake of large dextrans (∼70kD) occurs by macropinocytosis(*33*), so we investigated whether uptake of these large dextrans could be inhibited by YS-HepIII treatment. In complete media, containing full serum, macropinocytotic uptake of dextrans is low but measurable. When cells were starved overnight in serum-free minimal media, the uptake of dextrans increased significantly, but a short incubation with the macropinocytosis-specific inhibitor EIPA, (*33*), induced YS-HepIII, or purified HepIII enzyme inhibited dextran uptake (**Figure 4A**). Decreased dextran uptake achieved in purified HepIII- or YS-HepIII-treated cells mimicked that of EIPA, suggesting a specific effect on macropinocytosis (**Figure 4A**).

**Figure 4.**
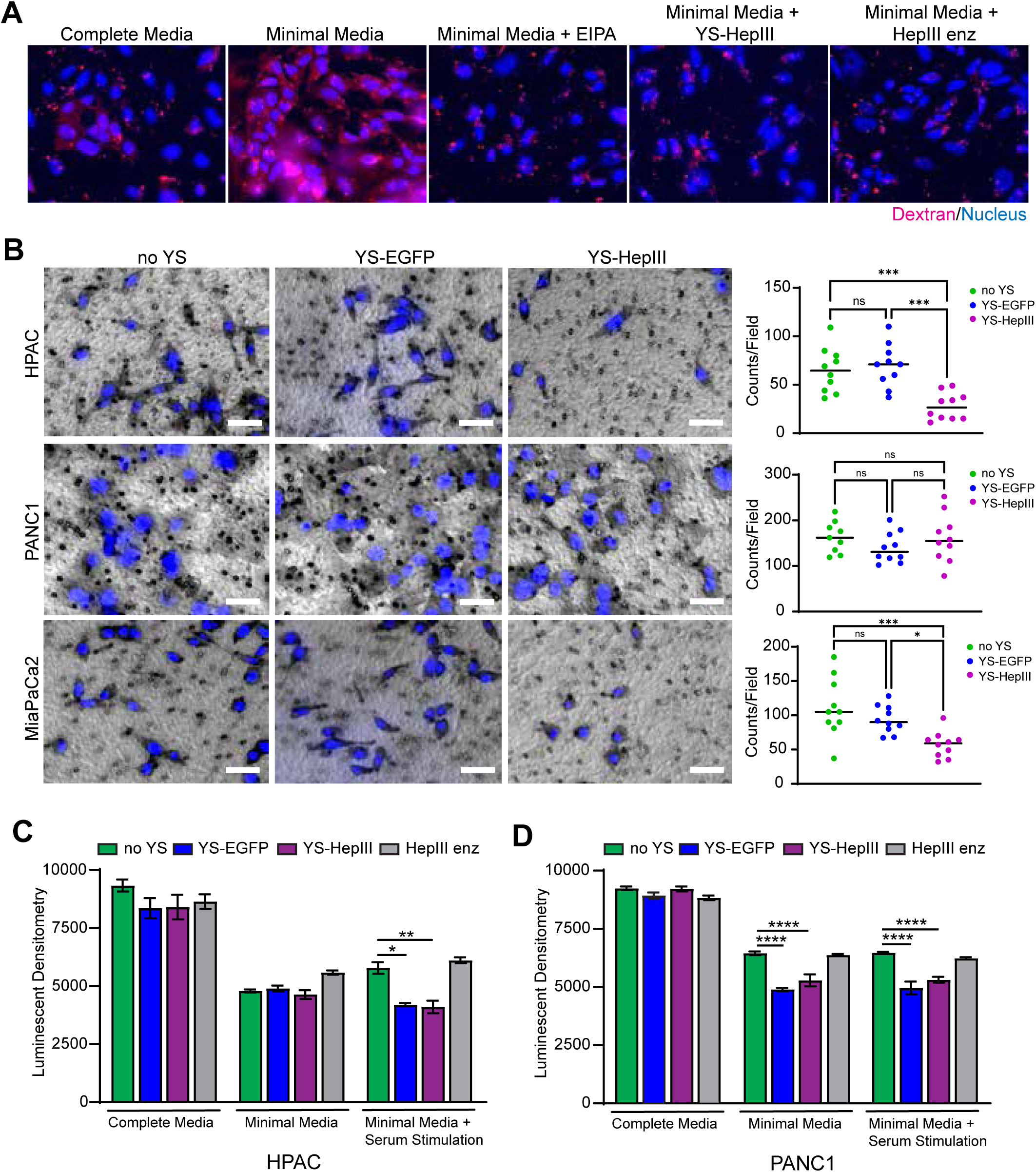
YS-HepIII decreases macropinocytosis and migration but does not significantly affect cell viability. (**A**) HPAC cells in culture were left in complete media or starved overnight in minimal media (serum-free, low amino-acids, low glucose) prior to 3-hour treatment with EIPA (50µM), YS-HepIII, or HepIII enzyme (5mU/mL). After treatment, cells were washed and 70kD dextrans conjugated to TexasRed were added to cells for 30 minutes (20mg/ml). After washing off un-endocytosed dextrans, cells were fixed and counterstained with DAPI. Fluorescence was imaged and representative images are shown. (**B**) HPAC, MIAPaCa-2, and PANC1 cells in monolayer culture were treated with PBS (no YS), pre-induced YS-EGFP, or pre-induced YS-HepIII for 2 hours. Cells were scraped and plated in the upper chambers of transwells in serum-free DMEM with full serum DMEM placed in lower chambers. Cells were allowed to migrate to bottom chambers for 20 hours and then cells left in the upper chamber were removed prior to fixing cells that had migrated to the lower chamber side of the membrane. Attached cells were stained with DAPI and fluorescence was overlayed with bright-field images. Representative images are shown (scale bar = 50µm). Cell nuclei counts per field (n=10 fields at 10x magnification) are shown to the right of images and significance was measured using a one-way ANOVA followed by Tukey’s post-hoc test for multiple comparisons *p<0.05, ***p<0.001. (**C-D**) HPAC and PANC1 cells were plated in triplicate and treated according to the following schedule: 1) “Complete media” wells were allowed to attach overnight in DMEM supplemented with 10% FBS and 2mM L-Glutamine and treated the next day with PBS (no YS), YS-EGFP, YS-HepIII, or HepIII enzyme for three hours prior to quantification of luminescence signal representing intracellular ATP levels; 2) “Minimal Media” and “Minimal Media + Serum Stimulation” wells were allowed to attach overnight in complete media followed by overnight starvation in minimal media (BME unsupplemented). The next day, all wells were treated with PBS (no YS), YS-EGFP, YS-HepIII, or HepIII enzyme for three hours. “Minimal Media” wells were then washed and luminescence signal was quantified; 3) “Minimal Media + Serum Stimulation” wells were stimulated with 10% FBS for 15 minutes after treatments were completed. Cells were then washed and luminescence signal was quantified. Significant changes in viability were analyzed using a one-way ANOVA followed by Tukey’s post-hoc test for multiple comparisons *p<0.05, **p<0.01, ****p<0.0001.

Additionally, inhibition of receptors like EGFR, FGFR, VEGFR, and focal-adhesion regulators like PAK2 (**Figure 3B, E**) is known to have a detrimental effect on cell migration(*89–92*). Using trans-well migration assays, we observed that the pancreatic cancer cell lines HPAC, PANC1, and MiaPaCa2 were capable of migration from serum-free compartments to those with serum-rich media. Pre-treatment with YS-EGFP did not significantly alter migration for any cell line, but pre-treatment of HPAC and MiaPaCa2 cell lines with YS-HepIII decreased migration significantly compared to untreated or YS-EGFP-treated controls (**Figure 4B**). Interestingly, migration of PANC1 cells, which did not show significant degradation of surface HS after treatment with YS-HepIII (**Figure 2A)**, was not significantly affected by YS-HepIII treatment (**Figure 4B**).

Decreased macropinocytosis and migration in starvation conditions as well as decreased growth factor signaling could all potentially be indicators of decreased cell viability. To assess viability, equal numbers of cells were plated *in vitro* under starvation conditions with and without serum stimulation after treatment with YS-EGFP, YS-HepIII, and HepIII enzyme, and ATP levels were quantified using a luminescence assay (**Figure 4C, D**). In HPAC cells, decreased viability was equally observed in cells treated with both YS-EGFP and YS-HepIII only in cells that were starved and then stimulated with serum. Under these conditions, addition of purified HepIII enzyme had no effect on viability compared to untreated cells, indicating that the observed decrease in viability is a specific effect of YS co-incubation (**Figure 4C**). PANC1 cells show similar effects in both starvation conditions and after serum stimulation, further supporting that any decreases in viability are effects of YS co-incubation alone (**Figure 4D**).

### YS-HepIII Degrades Tumor-Associated HS, but is not Sufficient to Inhibit Tumor Growth

*In vitro* assays involving co-incubation of YS and mammalian cancer cells lines are limited in the amount of time that co-incubation can occur due to accumulating waste products; thus, secondary and tertiary effects of YS-HepIII action may not be observed e.g. decreased viability. To overcome these limitations, observe the long-term effects of YS-HepIII-affected cell signaling, and focus on cancer cell metabolism without interference by the immune system, we utilized xenograft models of pancreatic cancer in immunodeficient mice. First, we confirmed colonization by YS of HPAC xenograft tumors that were treated with vehicle (no YS), YS-EGFP, or YS-HepIII, and quantified HS around the leading edge of colonized YS using tumors that had received three daily treatments of YS, and were harvested forty-eight hours after induction with L-arabinose that occurred twenty-four hours after the last YS treatment. Colonization of HPAC tumors was comparable between YS-EGFP and YS-HepIII tumors (**Figure 5A**), however significant HS degradation was only observed in YS-HepIII treated tumors (**Figure 5A**

**Figure 5.**
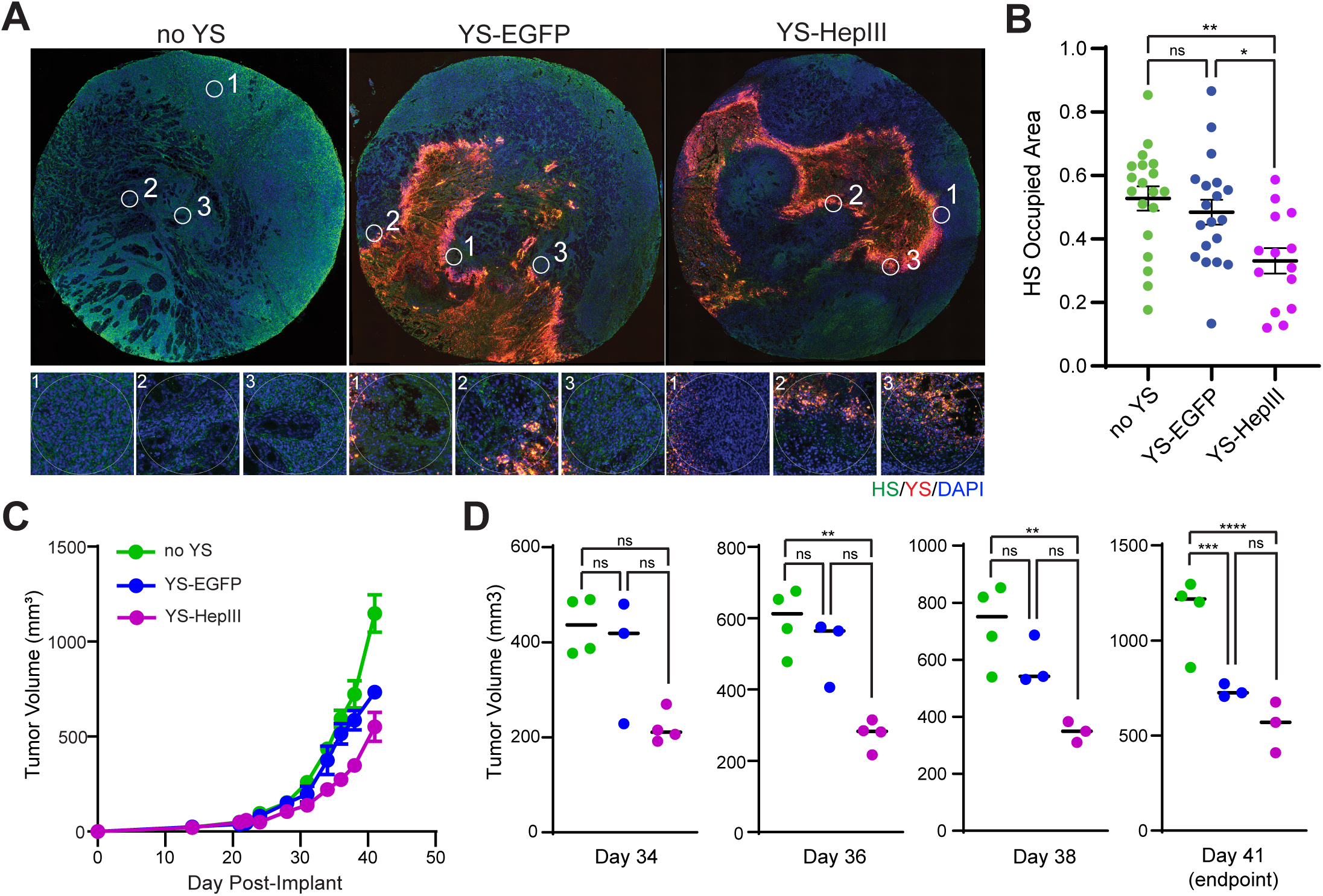
YS-HepIII Degrades Tumor-Derived Heparan Sulfate (HS) *in vivo* but Only Provides a Modest Therapeutic Effect on Tumor Growth. (**A**) Male and female (female n=8, male n=12) NSG mice aged 6-8 weeks were implanted subcutaneously with 1x10^6^ HPAC. Sex-matched littermates were randomized into groups and treatment began on Day 24 post-implantation (tumor volume averaged 95mm^3^). PBS (no YS), ST-EGFP, or ST-HepIII was administered by tail vein for three consecutive days at 2.5x10^6^ cfu/mouse. Forty-eight hours after the third treatment, mice were administered 40mg L-Arabinose intraperitoneally to induce EGFP or HepIII expression. One day after induction with L-arabinose, mice were euthanized and tumors were excised and fixed overnight in 4% paraformaldehyde at room temperature followed by processing and embedding in paraffin wax. Tumor sections were stained for HS (green) and YS (red), and nuclei were counterstained with DAPI (blue). Fluorescence was imaged for each tumor. Representative whole tumor images are shown with regions of interest (ROI, white circle) near YS-colonized areas are noted. Magnified images of representative ROI are shown below whole tumors. (**B**) HS occupied area in each ROI is shown as a dot plot with significance measured by one-way ANOVA followed by Tukey’s post-hoc test for multiple comparisons *p<0.05, **p<0.01. (C) Male and female (female n=6, male n=5) NSG mice were implanted subcutaneously with 1x10^6^ HPAC as above. Sex-matched littermates were randomized into groups and PBS (no YS), ST-EGFP, or ST-HepIII was administered by tail vein for three consecutive days at 2.5x10^6^ cfu/mouse. Forty-eight hours after the third treatment, mice were administered 40mg L-Arabinose intraperitoneally. Tumor measurements were taken thrice weekly, and tumor volume was measured using the following formula: ((short length^2^ x long length)/2. (D) Tumor volume differences were quantified at day 34, 36, 38, and 41 (experimental endpoint) and significance was determined using a one-way ANOVA followed by Tukey’s post-hoc test for multiple comparisons **p<0.01, ****p<0.0001.

(representative regions of interest), **Supplemental Figure 3**)**, B**). Despite specific degradation of HS *in vivo* by YS-HepIII, HPAC tumor growth was only mildly inhibited by the treatment, with most of the effect coming from YS alone (YS-EGFP and YS-HepIII having equally inhibited tumor growth by endpoint) (**Figure 5C, D**). These results are thus far consistent with most of the anticancer effect coming from YS treatment despite the unique inhibition of certain signaling pathways by YS-HepIII. To further delve into possible redundancies or compensations that are conferring resistance to our HS-targeted therapy, we next extensively profiled the metabolomic state of tumors post-treatment as well as gene expression signatures.

### Metabolite Profiling Confirms Increased Metabolic Stress after YS-HepIII Treatment

Unbiased metabolomics performed on the HPAC tumors that were harvested for **Figure 5** detected mostly amino acids, followed by organic acids, lipids, nucleotides, and a small percentage of carbohydrates (**Figure 6A**), and these categories were further broken into subclass e.g. “PC” phosphatidylcholine as a subclass of lipids, and represented as a heatmap comparing individual samples and individual metabolites by subclass (**Figure 6B**). A principal component analysis shows that the vehicle treated group was highly variable while YS-EGFP treated tumors clustered tightly for PC1 but diverged in PC2 (**Figure 6C**). Analysis of significantly altered metabolites revealed YS-HepIII-dependent increases in several lipids (a phosphatidylethanolamine (ChemSpider ID 24769153), a phosphatidylcholine (ChemSpider ID 24766646), and a lysophosphatidylglycerol (ChemSpider ID 7826081)) and a β-amino fatty acid (ChemSpider ID 260867) (**Figure 6D**). Typically, increases in these lipid subclasses would indicate a more rapidly dividing cell, needing more membrane components to continue proliferating(*93*), however, it is less known what significance an increase in β-amino fatty acids has on cancer cell metabolism. Metabolites significantly decreased by YS-HepIII treatment include a product of Prostaglandin E1 breakdown (2,3-dinor prostaglandin E1, ChemSpider ID 17220784) as well as a product of prostacyclin breakdown (2,3-dinor-6-keto-PGF1alpha, ChemSpider ID 4446210), a decreased amino fatty acid (4-acetamidobutyrate, ChemSpider ID 17180) and the Ala-Leu dipeptide (ChemSpider ID 87401) (**Figure 6D**). These changes indicate a higher persisting amount of Prostaglandin E1 and prostacyclin (due to less catabolism) which slow proliferation through their activation of adenylyl cyclase(*94, 95*). Decreased 4-acetamidobutyrate (an intermediate in GABA production from putrescine(*96*)) may slow proliferation by decreasing GABA catabolism and GABAergic signaling(*97, 98*).

**Figure 6.**
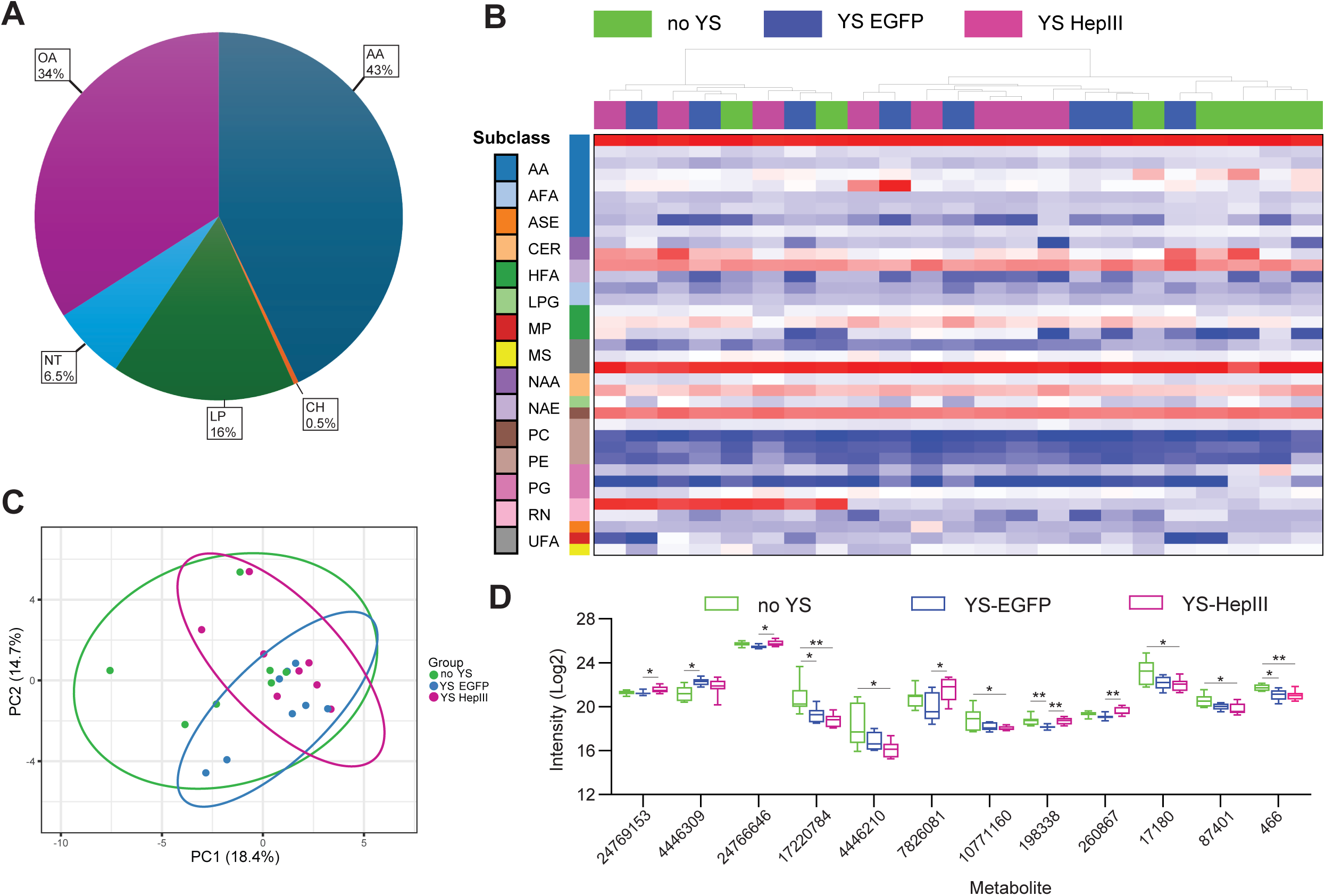
YS-HepIII Treatment Alters Metabolite Composition in Tumors. Established HPAC xenograft tumors in NSG mice were treated for three consecutive days with PBS (no YS), 2.5x10^6^ cfu/mouse YS-EGFP, or 2.5x10^6^ cfu/mouse YS-HepIII. Forty-eight hours after the third treatment, YS-EGFP and YS-HepIII-treated mice were administered 40mg/mouse L-arabinose to induce EGFP or HepIII expression. One day after induction with L-arabinose, tumors were excised and subject to untargeted metabolomic analysis. (**A**) Pie chart showing the distribution of discovered metabolites: Amino Acids (AA), Organic Acids (OA), Lipids (LP), Nucleotides (NT), Carbohydrates (CH). (**B**) Heatmap of total discovered metabolites arranged by sample and subclass: Amino Acid (AA), Amino Fatty Acid (AFA), Alkyl Sulfate Ester (ASE), Ceramide (CER), Hydroxy Fatty Acid (HFA), Lysophosphatidylglycerol (LPG), Monosaccharide Phosphate (MP), Monosaccharide (MS), N-acyl Amine (NAA), N-acyl Ethanolamine (NAE), Phosphatidylcholine (PC), Phosphatidylethanolamine (PE), Prostaglandin (PG), RNA Nucleoside (RN), Unsaturated Fatty Acid (UFA). (**C**) Principal component analyses of all discovered metabolites by treatment group. (**D**) Bar graph of significantly altered metabolites by treatment group. Metabolites are labeled with their ChemSpider ID: (2R)-3-{[(2-Aminoethoxy)(hydroxy)phosphoryl]oxy}-2-[(1Z,11Z)-1,11-octadecadien-1-yloxy]propyl (4Z,7Z,10Z,13Z,16Z,19Z)-4,7,10,13,16,19-docosahexaenoate (24769153), (5Z,7E,9S,11Z,14Z,17Z)-9-Hydroxy-5,7,11,14,17-icosapentaenoic acid (4446309), 1-palmitoyl-2-[(9Z,12Z,15Z)-octadecatrienoyl]-sn-glycero-3-phosphocholine (24766646), 2,3-dinor Prostaglandin E1 (17220784), 2,3-Dinor-6-keto-PGF1alpha (4446210), 3-({[(2S)-2,3-Dihydroxypropoxy](hydroxy)phosphoryl}oxy)-2-hydroxypropyl palmitate (7826081), 10-Undecenoic acid (10771160), 2-Aminoheptanoic acid (198338), 3-Aminooctanoic acid (260867), 4-acetamidobutyrate (17180), Ala-Leu dipeptide (87401), (+/-)-pantetheine (466). Statistical significance was analyzed using a one-way ANOVA followed by Tukey’s post-hoc test for multiple comparisons *p<0.05, **p<0.01.

A more in-depth lipid analysis showed a clearer picture of changes in lipid metabolism dependent on YS-HepIII treatment. In this lipid-specific analysis, many subclasses were detected, with the largest percentages belonging to triglycerols (TG) and phosphatidylcholines (PC) (**Figure 7A**). The magnitudes of each lipid detected are shown by sample and grouped into subclass as a heatmap (**Figure 7B).** A principal component analysis shows that lipid profiles of vehicle treated tumors (no YS) show little variation in either component compared to both YS treated groups, with YS-EGFP treated tumors showing the most variance in both components and YS-HepIII treated tumors lying between the other two groups (**Figure 7C**). Looking only at significantly affected lipids, storage TG make up the majority of these targets and overall are decreased, indicative of lipolysis caused by cell starvation (**Figure 7D**). Although fatty acid oxidation is a common strategy utilized by PDAC tumors to meet increased metabolic demand(*41*), the amplification of lipolysis after YS-HepIII treatment suggests a decrease in scavenged fatty acids to use for fuel (i.e. decreased macropinocytosis). Other lipids significantly impacted by YS-HepIII treatment compared to vehicle treatment include decreased acylcarnitine (AcCa) 20:1, PC 20:4_17:2, PC (P-15:2_24:0), and sphingomyelin (SM) (d38:2). Decreased AcCa would inhibit the fatty acid oxidation pathway(*99*), and decreased production may indicate a switch to other metabolic processes. As integral components of the plasma membrane(*100*), lack of PC and SM could result in halted cell proliferation.

**Figure 7.**
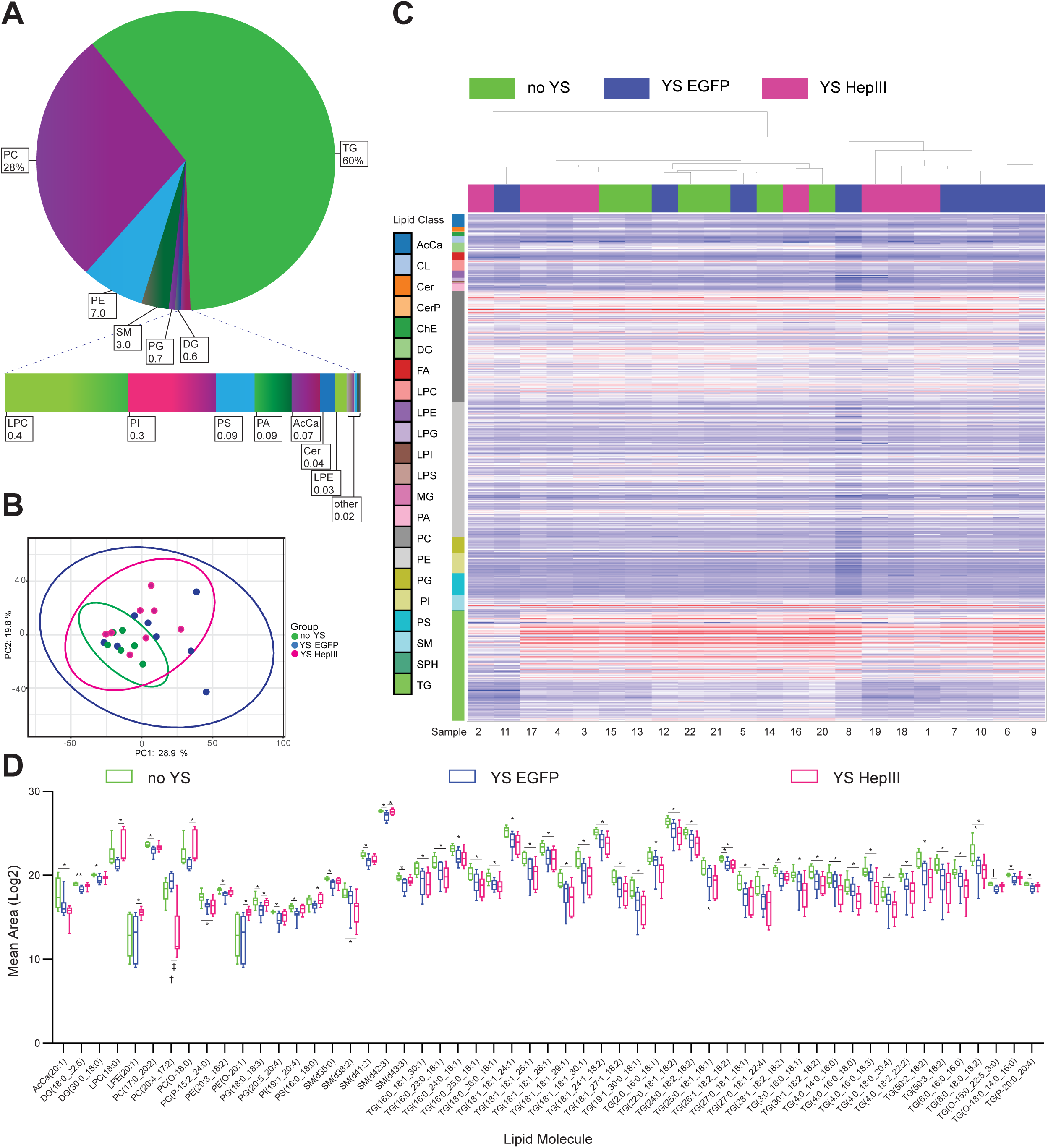
YS-HepIII Treatment Alters Lipid Composition in Tumors. Established HPAC xenograft tumors in NSG mice were treated for three consecutive days with PBS (no YS), 2.5x10^6^ cfu/mouse YS-EGFP, or 2.5x10^6^ cfu/mouse YS-HepIII. Forty-eight hours after the third treatment, YS-EGFP and YS-HepIII-treated mice were administered 40mg/mouse L-arabinose to induce EGFP or HepIII expression. One day after induction with L-arabinose, tumors were excised and subject to untargeted lipidomic analysis. (**A**) Pie chart showing the distribution of discovered lipids by subclass. (**B**) Heatmap of total discovered metabolites arranged by sample and subclass: Acylcarnitine (AcCa), Cardiolipin (CL), Ceramide (Cer), Ceramide phosphate (CerP), Cholesteryl ester (ChE), Diacylglycerol (DG), Fatty Acid (FA), Lysophosphatidylcholine (LPC), Lysophosphatidylethanolamine (LPE), Lysophosphatidylglycerol (LPG), Lysophosphatidylinositol (LPI), Lipopolysaccharide (LPS), Monoglyceride (MG), Phosphatidic Acid (PA), Phosphatidylcholine (PC), Phosphatidylethanolamine (PE), Glycerophosphoglycerol (PG), Phosphatidylinositol (PI), Phosphatidylserine (PS), Sphingomyelin (SM), Sphingosine (SPH), Triglycerol (TG). (**C**) Principal component analysis of all discovered lipids by treatment group. (**D**) Bar graph of significantly altered lipids by treatment group. Statistical significance was analyzed using a one-way ANOVA followed by Tukey’s post-hoc test for multiple comparisons *p<0.05, **p<0.01, †p<0.001, ‡p<0.0001.

### Gene Expression Analyses of Tumors Expose YS-HepIII-Mediated Effects on Major Nodes of Cancer Cell Survival

While metabolomic and lipidomic analyses of treated tumors support that YS-HepIII induces general metabolic stress and increased catabolism of lipid storage molecules, it must be noted that at the time of tumor harvest, YS have generally not colonized the entirety of the tumor, and effects may be diluted by uncolonized tumor regions. To explore transcriptional changes in regions of tumor cells restricted to those in contact with YS-HepIII (or control treatments), we employed spatial transcriptomics on tumors from **Figure 5A** using the NanoString GeoMx platform. Within tumor sections, regions of interest (ROI), equivalent to those used to quantify HS staining, that contained living tumor cells in conjunction with YS-HepIII or YS-EGFP were selected for spatial transcriptomics. We found that affected gene expression closely follows patterns seen in phosphoproteomics assays from **Figure 3**, in that YS-EGFP alone appears to have a significant effect on expression of metabolically relevant genes in HPAC tumor cells, likely due to a depletion of nutrients by YS that are being heavily scavenged to produce the transgene protein (EGFP). The expression of HepIII also synergizes with many YS-specific effects, and there are several transcripts uniquely affected by YS-HepIII treatment. When comparing vehicle (no YS) treated tumors with any YS-treated tumors (wherein the addition of HepIII was not significantly additive to the YS effect), expression changes included mostly downregulated genes relating to protein chaperones, translation initiation, transmembrane signaling receptors, and intracellular signaling molecules (**Figure 8A, Supplemental Table 1**). Genes that are synergistically affected by the addition of HepIII to YS largely include downregulated ribosomal proteins and other factors necessary for protein translation (**Figure 8B, Supplemental Table 2**). Another interesting set of genes includes those that are differentially regulated by YS-EGFP compared to both vehicle-treated tumors and YS-HepIII-treated tumors, and expression levels in vehicle- and YS-HepIII-treated tumors are equivalent (**Figure 8C**). Of these, it appears that the specific action of HepIII reverses tumor cell survival mechanisms that are activated during YS-EGFP colonization. Specifically, YS-EGFP treatment increases tumor cell expression of the glucose transporter type-1 gene (*SLC2A1*), genes for division and migration (*LAMC2, MZT2B*), and stress signaling molecules (*NDRG1, LCN2*) compared to vehicle treated tumors, while YS-HepIII rescues or even decreases their expression beyond that of vehicle treated tumors (**Figure 8C, Supplemental Table 3**). Lastly, we interrogated genes that are specific to the action of HepIII. Upregulated genes include those in the Wnt signaling pathway (*Wnt 9A, HOXB3*), SNARE-dependent exocytosis (*STX1B*), positive and negative regulators of intracellular signaling (*PTPRCAP, LDLRAD4*), and a fructose and mannose transporter (*SLC5A10*). Downregulated genes include a lactate exporter (*SLC16A3*), molecules related to cell motility (*PODXL, MIF, ITGB1*), a glycolytic enzyme (*GPI*), and a protease inhibitor (*CSTB*) (**Figure 8D, Supplemental Table 4**). Taking together all genes that are significantly affected by YS-HepIII compared to vehicle-treated tumors, an enrichment analysis of biological processes, pathways, and protein classes reveals the multi-faceted effects of YS-HepIII treatment on tumor cell gene expression (**Figure 8E, Supplemental Table 5).** Of the top 25 most significant pathways affected by YS-HepIII, there are clear effects on gene expression (transcription and splicing), translation (all steps), metabolism (RNA, proteins, and selenoamino-acids), apoptosis and the cell cycle, responses to stress including cellular starvation and amino acid deficiency, and pathways regulating migration, EMT, and metastasis (axon guidance, SLIT-ROBO expression and signaling, developmental biology) (**Figure 8E**). Targeting these various principal pathways reflects the ubiquitous nature of HS-based regulation, and the multitude of therapeutic effects that can occur through YS-HepIII treatment. With this gene expression data and data from our lipidomics studies, we next sought combination therapies that would exploit the vulnerabilities in cancer cell signaling and metabolism caused by YS-HepIII treatment.

**Figure 8.**
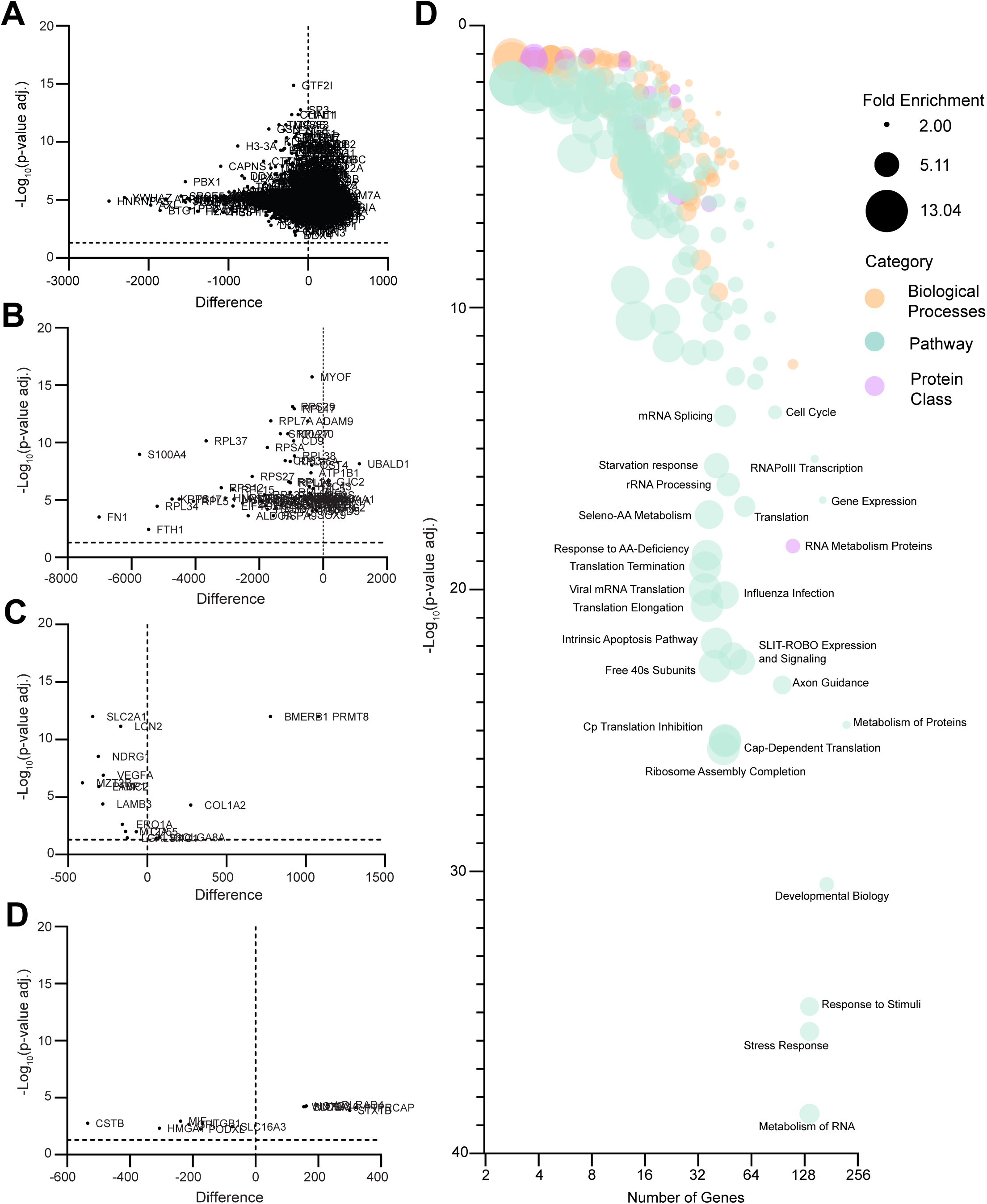
Spatial Transcriptomics of Tumor Cells Reveals Significant Changes in Expression of Genes for Important Cellular Processes in Regions of YS-HepIII Colonization. Established HPAC xenograft tumors in NSG mice were treated for three consecutive days with PBS (no YS), 2.5x10^6^ cfu/mouse YS-EGFP, or 2.5x10^6^ cfu/mouse YS-HepIII. Forty-eight hours after the third treatment, YS-EGFP and YS-HepIII-treated mice were administered 40mg/mouse L-arabinose to induce EGFP or HepIII expression. One day after induction with L-arabinose, tumors were excised, fixed, and paraffin embedded. Tumor sections were stained for HS (green) and YS (red), and nuclei were counterstained with DAPI (blue). Regions of interest (ROI) for spatial transcriptomic profiling were selected in the exact regions as those elected for HS quantification. Upwards of 18,000 genes were probed using the GeoMx® Human Whole Transcriptome Atlas. **(A)** Volcano plot of differentially expressed genes that were significant between no YS and YS-EGFP, and no YS and YS-HepIII, but not significant between YS-EGFP and YS-HepIII. Data represent averaged YS-EGFP and YS-HepIII values compared to no YS values. Each point is labeled with its gene name. **(B)** Volcano plot of differentially expressed genes that were significant between all three groups (no YS, YS-EGFP, and YS-HepIII) wherein YS-HepIII effects are additive to YS-EGFP effects in the same direction. Data represent YS-HepIII values compared to no YS values. Each point is labeled with its gene name. **(C)** Volcano plot of differentially expressed genes that were significant between no YS and YS-EGFP, and YS-EGFP and YS-HepIII, but not significant between no YS and YS-HepIII. Data represent YS-HepIII values compared to YS-EGFP values. Each point is labeled with its gene name. **(D)** Volcano plot of differentially expressed genes that were significant between no YS and YS-HepIII, and YS-EGFP and YS-HepIII, but not significant between no YS and YS-EGFP. Data represent averaged no YS and YS-EGFP values compared to YS-HepIII values. Each point is labeled with its gene name. **(E)** All genes significantly different between YS-HepIII and no YS groups were ranked and analyzed for statistical enrichment in Biological Processes, Pathways, and Protein Classes. Statistically significant terms are presented as a bubble graph organized by -Log_10_ p-values and number of genes from the total gene set found in each term gene set. Bubble color is used to differentiate the categories Biological Process (orange), Pathway (aqua), and Protein Class (purple), and bubble sizes correspond to fold enrichment. The top 25 most significant pathways are labeled.

### YS-HepIII Treatment Synergizes with Fatty Acid Synthase Inhibition but not mTOR Inhibition

PDAC relies heavily on fatty acid oxidation for ATP production and upregulates genes for *de novo* fatty acid synthesis(*41, 101*), but macropinocytosis can confer resistance to fatty acid synthase inhibition(*80*). In light of our data showing tremendous lipolysis of the lipid storage TG *in vivo,* in an environment where macropinocytosis is likely decreased, and less fatty acids are replenished from the TME, we hypothesized that fatty acid synthesis would be particularly vulnerable to inhibition(*80*). Indeed, upregulation of genes following YS-HepIII treatment such as ELOVL5, which encodes a fatty-acid elongase, supports an increase in fatty acid synthesis(*102, 103*). Using the HPAC xenograft model in NSG mice, we found that the fatty acid synthase inhibitor GSK2194069 (FASNi) synergized with YS-HepIII treatment to significantly inhibit tumor growth (**Figure 9A**). No synergistic effects were observed when GSK2194069 treatment was combined with YS-EGFP indicating HepIII-specific synergy.

**Figure 9.**
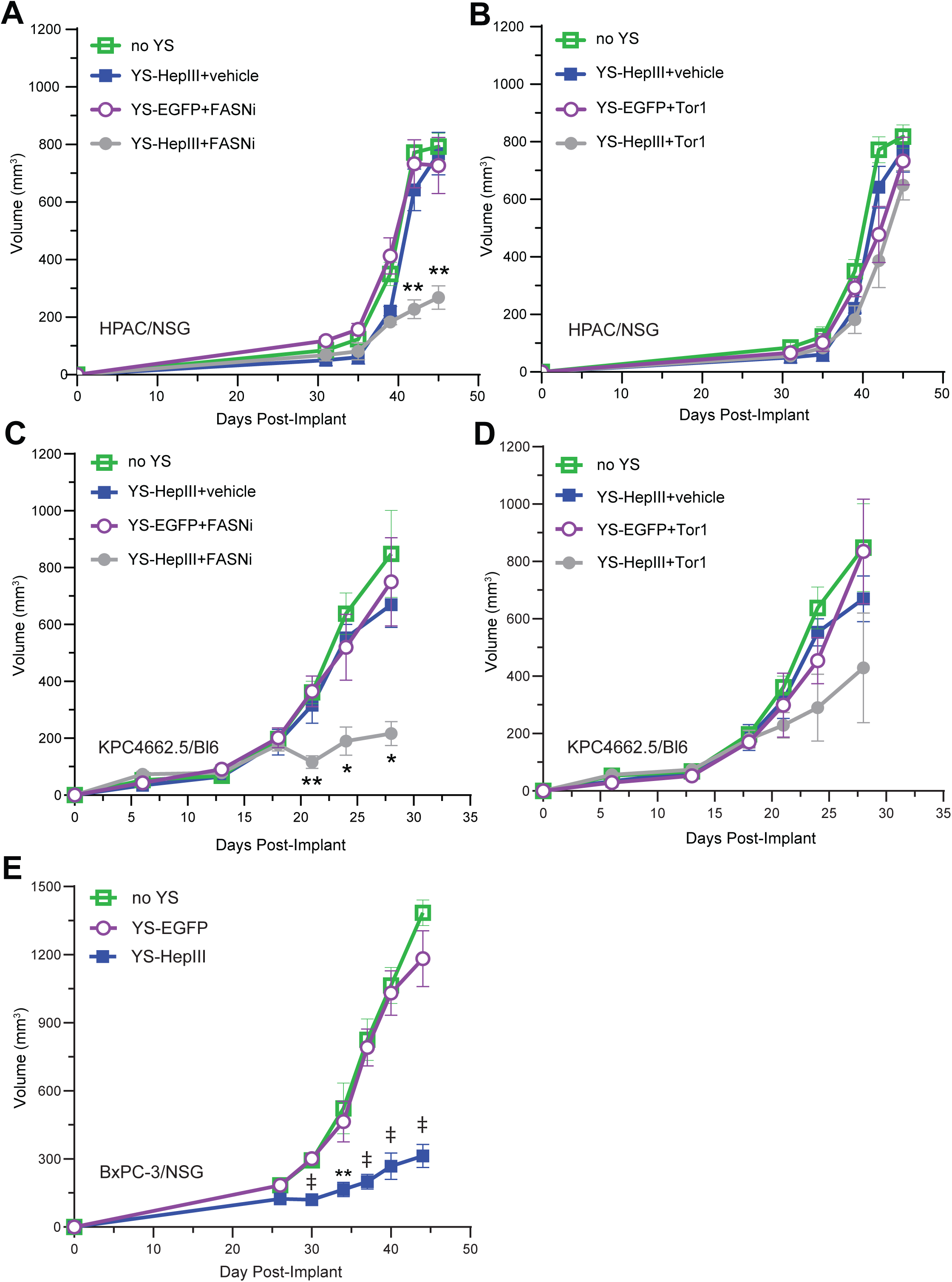
YS-HepIII Treatment Synergizes with Fatty Acid Synthase Inhibition. (**A-B**) Male and female NSG mice aged 6-8 weeks were implanted subcutaneously with 1x10^6^ HPAC cells into the area near the right flank in a total volume of 100µl HBSS. Sex-matched littermates were randomized into groups and treatment began on Day 31 post-implantation. YS-EGFP or YS-HepIII was administered by tail vein after subculturing to 0.7 O.D. Mice were administered YS for three consecutive days at 2.5x10^6^ cfu/mouse. Forty-eight hours after the third treatment, mice were administered 40mg L-Arabinose intraperitoneally. Tumor measurements were taken thrice weekly, and tumor volume was measured using the following formula: ((short length^2^ * long length)/2). (**A**) Therapeutic curve of YS-HepIII and fatty acid synthase inhibitor (FASNi) treatment combination in immune-deficient mice. NSG mice bearing HPAC xenograft tumors were split into the following treatment groups: 1) no YS (n=4), 2) YS-HepIII + vehicle (n=3), 3) YS-EGFP + FASNi (n=3), 4) YS-HepIII + FASNi (n=4). Starting on the day of L-arabinose induction, groups with vehicle treatment were given 10% DMSO in PBS and groups with FASNi treatment were given 40µg GSK2194069 in vehicle both by oral gavage every other day until experimental endpoint. (**B**) Therapeutic curve of YS-HepIII and the mTOR inhibitor Torin1 (Tor1) treatment combination in immune-deficient mice. NSG mice bearing HPAC xenograft tumors were split into the following treatment groups: 1) no YS (n=4), 2) YS-HepIII + vehicle (n=3), 3) YS-EGFP + Tor1 (n=4), 4) YS-HepIII + Tor1 (n=4). Starting on the day of L-arabinose induction, groups with vehicle treatment were given 10% DMSO in PBS and groups with Tor1 treatment were given 200µg Tor1 in vehicle both by intraperitoneal injection every other day until experimental endpoint. (**C-D**) Male and female C57Bl/6 mice aged 6-8 weeks were implanted subcutaneously with 2x10^5^ KPC4662.5 cells into the area near the right flank in a total volume of 100µl HBSS. Sex-matched littermates were randomized into groups and treatment began on Day 13 post-implantation. YS-EGFP or YS-HepIII administration and induction followed the same dosing, schedule, and route as **A**,**B**. Tumor measurements were taken thrice weekly, and tumor volume was measured using the following formula: ((short length^2^ * long length)/2). (**C**) Therapeutic curve of YS-HepIII and FASNi treatment combination in immunocompetent mice. C57Bl/6 mice bearing KPC4662.5 tumors were split into the following treatment groups: 1) no YS (n=4), 2) YS-HepIII + vehicle (n=4), 3) YS-EGFP + FASNi (n=4), 4) YS-HepIII + FASNi (n=4). Starting on the day of L-arabinose induction, groups with vehicle treatment were given 10% DMSO in PBS and groups with FASNi treatment were given 40µg GSK2194069 in vehicle both by oral gavage every other day until experimental endpoint. (**D**) Therapeutic curve of YS-HepIII and the Tor1 treatment combination in immunocompetent mice. C57Bl/6 mice bearing KPC4662.5 tumors were split into the following treatment groups: 1) no YS (n=4), 2) YS-HepIII + vehicle (n=3), 3) YS-EGFP + Tor1 (n=3), 4) YS-HepIII + Tor1 (n=4). Starting on the day of L-arabinose induction, groups with vehicle treatment were given 10% DMSO in PBS and groups with Tor1 treatment were given 200µg Tor1 in vehicle both by intraperitoneal injection every other day until experimental endpoint. (**A-D**) Once control group tumors reached a maximum diameter of 15mm, mice were euthanized. Statistically significant changes in tumor growth was analyzed using unpaired t-tests by date post-implantation of YS-HepIII+FASNi or Tor1 compared to YS-EGFP+FASNi or Tor1, **p<0.01. (**E**) Male and female NSG mice aged 6-8 weeks were implanted subcutaneously with 1x10^6^ BxPC-3 cells into the area near the right flank in a total volume of 100µl HBSS. Sex-matched littermates were randomized into groups and treatment began on Day 26 post-implantation. No YS (n=4), YS-EGFP (n=5), or YS-HepIII (n=7) administration and induction followed the same dosing, schedule, and route as **A**,**B**. Tumor measurements were taken thrice weekly, and tumor volume was measured using the following formula: ((short length^2^ * long length)/2). Once control group tumors reached a maximum diameter of 15mm, mice were euthanized. Statistically significant changes in tumor growth was analyzed using unpaired t-tests by date post-implantation of YS-HepIII compared to YS-EGFP, **p<0.01, ‡p<0.0001.

Macropinocytosis has also been shown to confer sensitivity to mTOR inhibitors(*104*), however, we found no synergistic inhibition of HPAC tumor growth when combining the mTOR signaling inhibitor Torin1 to YS-HepIII treatment (**Figure 9B**). As YS-HepIII treatment alone showed tremendous downregulation of ribosomal protein genes, decreased availability of the Ala-Leu dipeptide, and enrichment of genes in cellular starvation and amino acid deficiency response, we hypothesized that the mTOR pathway may have already been significantly inhibited after YS-HepIII treatment. This result underscores the global effects that HS degradation has on multiple mechanisms of PDAC cell survival. Interestingly, these combination studies performed similarly in immunocompetent mice (**Figure 9C, D**) implanted with a syngeneic KRAS- mutant pancreatic cancer cell line, show that synergy between YS-HepIII and FASNi but not mTOR inhibition continues in the presence of a more physiologically relevant tumor microenvironment.

As mutant KRAS upregulates SDC1 on tumor cell surfaces, we tested the anti-tumor efficacy of YS-HepIII in a non-KRAS mutant PDAC cell line, BxPC-3. Interestingly, treatment with YS-HepIII was able to control tumor growth of BxPC-3 from the first treatment and maintained growth control well after the last treatment (**Figure 9E**). While BxPC-3 cells don’t harbor a KRAS mutation, they do have mutations in *SMAD4* and *TP53* similar to primary PDAC tumors. These results may indicate that YS-HepIII treatment can counteract the effects of these mutations, and YS-HepIII may synergize particularly with KRAS inhibitors.

These therapeutic combination experiments highlight the importance of YS-HepIII in interfering with macropinocytosis, but the data provided in this manuscript also suggest a number of new genes, proteins, and pathways that might be simultaneously targeted to achieve synergistic inhibition of PDAC tumor growth. Future studies will focus on new therapeutic combination strategies based on these data in conjunction with effects on the immune microenvironment of tumors.

## Discussion

Patients diagnosed with PDAC will typically receive FOLFIRINOX or gemcitabine with nab-paclitaxel, however, many patients will experience treatment-limiting toxicities, and most tumors will develop resistance(*105, 106*). Major mechanisms of innate or acquired resistance to chemotherapy in PDAC tumors include metabolic and cellular signaling plasticity, extreme fibrosis, upregulation of macropinocytosis and autophagy, upregulation of drug efflux pumps, and cooperative signaling with pro-tumorigenic immune cells(*78*). Although therapeutic strategies exist that target these mechanisms, such as the multi-target tyrosine kinase inhibitor sorafenib or the metabolic inhibitor devimistat, these drugs and others that target only one or a few fundamental PDAC processes have failed to improve outcomes for patients(*107, 108*).

Despite this, several new, targeted therapies directed at specific PDAC drivers have had some success: 1) the use of PARP inhibitor treatment for patients with germline BRCA mutations(*109*), 2) a 42% response rate after treatment of NRG1-fusion positive PDAC with Zenocutuzumab, and 3) mutant KRAS-specific targeting (G12C) with sotorasib(*110*) and adagrasib(*111*). Unfortunately, these mutations occur in less than 10% of total PDAC cases, and even emerging treatments for more prolific mutations (e.g. KRAS G12D occurring in about 40% of PDAC) are seeing objective response rates of only ∼30% with little data on whether resistance will be developed(*112–114*). Thus, treatment strategies that target multiple PDAC progression mechanisms will ultimately be needed to prevent resistance and improve patient survival.

In this study, we have targeted several survival pathways in PDAC by expressing HepIII on the surface of attenuated, tumor-targeting *Salmonella*. Our results indicate that YS-HepIII treatment alone is capable of decreasing signaling downstream of several HSPG and co-receptors, inhibiting macropinocytosis, and inhibiting migration. The transcriptional changes occurring due to HepIII enzymatic action or YS-HepIII synergy largely indicate that translation is being halted due to metabolic stress. This is supported by both the metabolomic and lipidomic analyses showing that cells are depleting storage lipids for energy while experiencing a lack of essential amino acids and necessary membrane components needed for growth and proliferation. Decreasing scavenging of ATP through decreased macropinocytosis may also inhibit drug efflux through ATP-dependent transporters(*115*).

Despite these significant effects, tumor control was not achieved with YS-HepIII treatment alone, but rather primed tumors for response to treatment with a fatty acid synthase inhibitor. While there is a myriad of other therapeutic molecules that might synergize with YS-HepIII therapy (e.g. gemcitabine, autophagy inhibitors), we first sought proof-of-concept experiments that would show how our omics data could help predict which combination therapies would achieve tumor control. Our goal is to continue this work to mechanistically interrogate possible mechanisms of YS-HepIII resistance highlighted in our data i.e. combining YS-HepIII therapy with inhibitors of Erbb3, Met, CSF1R, ALK, and NuaK1 and 2, combining YS-HepIII with inhibitors of PC and PE synthesis (in response to increased species found in our metabolomics data), and in response to our transcriptomic data, combining YS-HepIII with inhibitors of Wnt, SLC5A10, and SLC16A3. Performing these experiments in immune competent models will elucidate whether these treatment combinations remain efficacious in a more physiologically relevant setting.

Although previous studies have described the anti-tumor effects of HepIII treatment on cells and tumor models(*116, 117*), the data presented in this manuscript using specific and precise expression of active HepIII in tumors supports the translational potential of this treatment strategy for PDAC. Our data indicate that YS-HepIII treatment will be particularly useful to improve the efficacy of FDA-approved treatments or those currently in clinical trials such as the KRAS inhibitor daraxonrasib and the fatty acid synthase inhibitor TVB-2640. Improving treatment efficacy and preventing resistance is desperately needed to improve survival for patients with PDAC.

## Materials and Methods

### Generation of ST-HepIII and Maintenance of Cultures

Heparinase III amino acid sequences from *Flavobacterium heparinum* (UniProt (RRID:SCR_002380) Q59289), *Bacteroides stercoris* (UniProt C7EXL6), and *Bacteroides eggerthii* (UniProt A0A380Z8I2) were modified to remove the original signal peptides to prevent periplasmic targeting, and the type III secretion signal from the *Salmonella* flagellin gene (FliC)(*118*) was added to the N-terminus to target HepIII protein to the outer membrane. 6xHis tag sequences were added to the C-terminus. Amino acid sequences were reverse-translated and codon optimized for *Salmonella typhimurium* expression using JCat(*119*), and the resulting cDNA sequences were synthesized by Biomatik (Kitchener, Ontario, Canada). Full sequences are available in *Supplemental Materials and Methods*. Synthesized HepIII sequences were inserted in-frame into EGFP-pBAD (RRID:Addgene_54762, with cloning removing EGFP) and electroporated into YS1646 as previously described(*120*). YS1646 was obtained from ATCC (202165) and cultured in modified Luria Broth (LB) media containing Mg(C₂H₃O₂)₂ and CaCl_2_ in place of NaCl. Cultures and stocks were maintained without addition of L-Arabinose (A3256, Sigma-Aldrich, St. Louis, MO, USA).

### Confirmation of HepIII Expression and Enzymatic Activity, and Culture Viability

pBAD-HepIII and pBAD-EGFP positive YS clones identified by colony PCR were cultured in LB to 0.7 OD and then supplemented without (0.000%) or with varying concentrations of L-Arabinose for 45 minutes. Pellets were lysed in RIPA buffer (89900, Thermo Scientific, Waltham, MA, USA) supplemented with protease inhibitor cocktail (11836170001, Roche, Basel, Switzerland). Western blot of lysates was performed to detect EGFP (anti-EGFP antibody, sc-9996, Santa Cruz Biotechnology, Dallas, TX, USA, RRID:AB_627695) or HepIII (anti-His Tag antibody, 10001-0-AP, Proteintech, Rosemont, IL, USA, RRID:AB_11232228). Transgenic HepIII enzymatic activity against purified heparan sulfate (HS) (S5992, Selleck Chemicals, Houston, TX, USA) was assessed using a gel-shift assay. Pre-induced YS cultures or purified, recombinant HepIII and HepI enzymes (5mU/mL) (6145-GH-010 and 7897-GH-010, Bio-Techne, Minneapolis, MN, USA) were incubated overnight with 1mg/mL HS in a shaking incubator at 37℃. The next day, cultures were pelleted and supernatants were electrophoresed using 12% polyacrylamide gels. Gels were then stained in 0.5% Alcian Blue (A5268, Sigma-Aldrich) (in 7:2:1 H_2_O, EtOH, AcOH) to visualize HS smears. To visualize bacterial viability, YS-HepIII cultures were grown overnight and sub-cultured to 0.7 OD the next day. Subcultures were induced with 0.5% L-Arabinose for 1hr at 37℃, shaking, and pellets were then stained with a combination of acridine orange (2mg/mL, A6014, Sigma-Aldrich) and ethidium bromide (1mg/mL, 17898, ThermoFisher). Fluorescence was visualized using the Widefield Observer 7 (Zeiss) and images were captured with the AxioCam 702 sCMOS Mono digital camera (Zeiss). Images were analyzed using Zen Blue 3.12 (Zeiss, RRID:SCR_013672).

### Human PDAC Cell Lines and Culture Conditions

HPAC (CRL-2119, RRID:CVCL_3517), MIAPaCa2 (CRL-1420, RRID:CVCL_0428), and PANC1 (CRL1469, RRID:CVCL_0480) human PDAC cell lines were obtained from ATCC and maintained in complete media consisting of DMEM (10013CV, Corning, Glendale, AZ, USA) supplemented with 10% FBS (Gemini), 2mM L-Glutamine and penicillin/streptomycin (ThermoFisher). The BxPC-3 (CRL-1687, RRID:CVCL_0186) cell line was also obtained from ATCC and maintained in RPMI media (ThermoFisher) supplemented with 10% FBS (Gemini), 2mM L-Glutamine and penicillin/streptomycin (ThermoFisher). Cells were passed <5 times prior to *in vitro* experiments or implantation *in vivo*.

### Digestion of HS on Live Cancer Cell Lines in 2D Culture

#### Immunofluorescence

HPAC, MIAPaCa-2, PANC1, and BxPC-3 cells were plated to 80-90% confluency in 8-well chamber slides (PEZGS0896, MilliporeSigma) overnight in complete DMEM or RPMI without antibiotics. The next day, pre-induced YS-EGFP or YS-HepIII at a concentration of 1000:1 (bacteria: mammalian cells) or HepIII enzyme (5mU/mL, Bio-Techne) was added to cells for three hours at 37℃. At the end of the incubation period, cells were washed 3x with PBS to remove YS and then fixed in 4% paraformaldehyde for 10 minutes at room temperature. Fixed cells were washed 3x with PBS and blocked in 5% normal goat serum for 1hr at room temperature. Cell surface HS was stained using a biotinylated, anti-HS antibody (370255-B-50H, AMS Biotechnology, Milton, England) and nuclei were counterstained with DAPI. HS staining was visualized using AlexaFluor488 conjugated avidin (21832, Invitrogen, Waltham, MA, USA). Fluorescence was visualized using the Widefield Observer 7 (Zeiss) and images were captured with the AxioCam 702 sCMOS Mono digital camera (Zeiss). Images were analyzed using Zen Blue 3.12 (Zeiss).

#### Western blot

HPAC, MIAPaCa-2, PANC1, and BxPC-3 cells were plated to 80-90% confluency 6-well plates and treated as above. After three hours of incubation with YS or enzyme treatments, cells were washed 3x in PBS and lysed in RIPA buffer (ThermoFisher) supplemented with protease inhibitor cocktail (Roche). Protein lysates (70µg) were probed with an anti-Syndecan-1 antibody (12922, Cell Signaling Technology, Danvers, MA, USA) by western blot and visualized with fluorescent secondary antibody (AlexaFluor555, A-21422, ThermoFisher).

### Kaplan-Meier Analysis

Syndecan 1-4 mRNA expression in relation to patient survival was analyzed from KM plotter(*121*) (https://kmplot.com/analysis). The pancreatic ductal adenocarcinoma dataset (n=177) was selected for analysis from the “Pan-Cancer mRNA-seq” module. Analysis was not restricted to any “subtype” or “cellular content”, and overall survival was computed after electing “auto-select best cutoff: percentiles”. Hazard ratios with confidence intervals are reported along with p-values from log-rank tests.

### Phosphoproteomic Microarray Analysis and Kinase-Substrate Enrichment Assay

HPAC cells were seeded in 10cm tissue culture dishes and allowed to grow to 80-90% confluency overnight in DMEM supplemented with 10% FBS (Gemini) and 2mM L-Glutamine (ThermoFisher), but without antibiotics. After overnight culture in complete media, cells were washed 3x with PBS and cultured overnight in Basal Medium Eagle (BME, 21010046, ThermoFisher). The next day, cells were treated with YS (YS-HepIII or YS-EGFP) with 1x10^9^ cfu in BME for three hours. At the end of the three hours plates were washed 3x with PBS and complete media was added to some plates to stimulate cells for 15 minutes. The four treatment groups include 1) untreated, unstimulated cells, 2) untreated, stimulated cells, 3) YS-EGFP treated, stimulated cells, and 4) YS-HepIII treated, stimulated cells. At completion of the stimulation timepoint, all cells were placed on ice and washed immediately in ice-cold PBS, followed by lysis in RIPA with protease and phosphatase inhibitors (78442, ThermoFisher). Phosphoproteomic results were obtained by adding equal protein amounts of these lysates (20µg) to a human phosphorylation screening array according to the manufacturer’s instructions, and having the manufacturers scan the array and report back raw data (AAH-BLG-PHO1-4, RayBiotech, Peachtree Corners, GA, USA). Raw data was analyzed using the background subtracted F532 median – B532 column data entered into the provided L-Analysis Tool (RayBiotech) to retrieve data normalized to microarray positive and negative controls after averaging technical duplicates. For phosphoproteins with data on total protein amounts from the microarray, those targets were first normalized to total protein (e.g. Akt1). These data were then normalized to levels from the untreated, unstimulated group for each target and Log2 transformed. Statistically significant targets for the phosphoproteomic microarray included any target whose fold change was greater than 1.5 or less than 0.65 fold according to manufacturer’s instructions (RayBiotech). A kinase-substrate enrichment assay to detect kinase activity based on our phospho-data was run using the RoKAI algorithm(*122*) through the RokaiXplorer web application (http://explorer.rokai.io).

### Cell Line Culture Macropinocytosis Assay

HPAC cells were seeded in 8-well chamber slides, cultured overnight, starved in minimal media, and treated with YS-EGFP or YS-HepIII cultures for three hours as was previously described for the Phosphoproteomic Microarray. In addition, starved cells were treated with 50µM Ethylisopropylamiloride (EIPA). After three hours of treatment, cells were washed in PBS and 70kDa dextrans conjugated to Texas Red (D1864, ThermoFisher) were added to cells for 30 minutes at a concentration of 20 mg/mL. After dextran incubation, cells were washed on ice in cold PBS x5 washes to remove un-endocytosed dextrans. Cells were then fixed on ice in 4% paraformaldehyde for 12 minutes. Nuclei were counterstained with DAPI at 1µg/mL for 10 minutes at room temperature. After two washes in PBS, slides were mounted with Fluoromount-G (0100-01, Southern Biotech, Birmingham, AL, USA).

### Migration Assay

Migration was measured using a transwell assay with 8µm pores (cat# 3422, Corning). Cell lines in culture were untreated (no YS) or pre-treated with induced YS-EGFP or YS-HepIII (1000:1) for two hours. Cells were scraped, pipetted into a single cell suspension and counted for placement into the upper chamber of the transwell in serum-free DMEM. Cells were allowed to migrate toward the lower chamber (DMEM + 10% FBS) for 20 hours. Membranes were then fixed and stained with DAPI for brightfield and fluorescent imaging with a 10x objective (Widefield Observer 7, Zeiss). Cell nuclei per field (n=10 fields) were counted using CellProfiler (www.cellprofiler.org, The Broad Institute, RRID:SCR_007358).

### Cell Culture Viability Assay

HPAC and PANC1 cells were plated in 96-well plates to 70% confluency. After attachment overnight, cells in complete media (without antibiotics) were co-incubated with PBS (no YS), pre-induced YS-EGFP, pre-induced YS-HepIII, or HepIII enzyme (5mU/mL) for three hours. Other wells were serum-starved overnight in minimal media (BME without serum) and then co-incubated with PBS (no YS), pre-induced YS-EGFP, pre-induced YS-HepIII, or HepIII enzyme (5mU/mL) for three hours prior to 15-minute stimulation with 10% FBS (for “Serum Stimulation” samples only). After completion of treatment and/or stimulation, cells were washed and analyzed for viability using the CellTiter-Glo 2.0^®^ assay (G9241, Promega, Madison, WI, USA) according to manufacturer’s instructions. Briefly, CellTiter-Glo^®^ 2.0 reagent was added to washed cells as a 1:1 volume to total volume of wash media. Wells were mixed using an orbital shaker to lyse cells and liberate ATP from viable cells. Luminescence released by binding of the reagent to free ATP was measured using the SpectraMax iD3 Multimode plate reader (Molecular Devices, San Jose, CA, USA).

### Xenograft Tumor Growth and Treatment, Therapeutic Studies

NOD/SCID gamma (NSG) mice (RRID:IMSR_JAX:005557) were obtained from breeding colonies housed at the City of Hope (COH) Animal Research Center and, for all studies, handled according to standard Institutional Animal Care and Use Committee guidelines (IACUC Protocol # 22037, approved 04/20/2022). Male and female NSG mice aged 6-8 weeks were implanted subcutaneously with 1x10^6^ HPAC cells into the area near the right flank in a total volume of 100µl HBSS. Growth dynamics were not found to be significantly different between sexes (female n=8, male n=12, ANOVA followed by Tukey’s post-hoc test, data not shown). Sex-matched littermates were randomized into groups and treatment began on Day 24 post-implantation (tumor volume averaged 95mm^3^). YS-EGFP or YS-HepIII was administered by tail vein after subculturing to 0.7 O.D. Mice were administered YS for three consecutive days at 2.5x10^6^ cfu/mouse. Forty-eight hours after the third treatment, mice were administered 40mg L-Arabinose intraperitoneally. Tumor measurements were taken thrice weekly, and tumor volume was measured using the following formula: ((short length^2^ * long length)/2). For combination therapy studies, mice were administered YS and induced at the same dose and schedule as above, but on the day of L-arabinose induction, treatments with Torin1 (200µg/mouse by intraperitoneal injection), GSK2194069 (40µg/mouse by oral gavage), or vehicle (10% DMSO in PBS for both) were begun. Torin1, GSK2194069, or vehicle treatments were administered every other day until maximum tumor growth (15mm diameter) was reached in control groups. No significant signs of weight loss or distress were observed in any treatment group.

### In Vivo YS Colonization, Spatial HS Degradation, and Spatial Transcriptomics

HPAC tumors were grown subcutaneously in NSG mice, as above, until day 27 post-implantation (tumor volume average = 130mm^3^) when YS treatment via tail vein began. Forty-eight hours after the third treatment, mice were administered 40mg L-Arabinose intraperitoneally. Twenty-four hours after induction with L-arabinose, mice were euthanized and tumors were excised and divided into thirds. One third was fixed overnight in 4% paraformaldehyde at room temperature followed by processing end embedding in paraffin wax. These blocks were sliced at 5µm sections and stained for HS (biotinylated anti-HS antibody, 370255-B-50H, AMS Biotechnology), nuclei (DAPI), and YS (anti-Salmonella O-antigen conjugated to AlexaFluor647, NB110-16952AF647 RRID:AB_3175574, Novus Biologicals, Centennial, CO, USA, RRID:SCR_004286) without permeabilization or antigen retrieval. Fluorescence was imaged using the Widefield Observer 7 (Zeiss) with the AxioCam 702 sCMOS Mono digital camera (Zeiss). Images were analyzed using Zen Blue 3.12 (Zeiss) and regions of interest (ROI) near YS-colonized regions of tumors were selected for HS degradation analysis. HS degradation in each ROI was calculated using CellProfiler. The other two-thirds of tumor tissues were flash frozen separately for untargeted metabolomic and lipidomic analyses (below).

#### Spatial Transcriptomics

Spatial transcriptomic profiling was performed using the GeoMx-Digital Spatial Profiler (GeoMx-DSP, Nanostring Technologies, Seattle, WA, USA). The 5-um FFPE sections of HPAC tumors were processed following manufacturer’s protocol. Briefly, the slides were deparaffinized and subject to antigen retrieval by proteinase K treatment. The slices were then hybridized with the GeoMx Probe Mix (human whole transcriptome Atlas (GeoMx Hu WTA) panel), (121401102, Nanostring Technologies) overnight at 37°C. The GeoMx Hu WTA panel includes 1,8000+ genes representative of key cancer cell biology and the tumor microenvironment. To visualize the specific markers for regions of interest (ROI), the slides were stained with GeoMx Solid Tumor TME Morphology Kit (Cat. #: 121300310, Nanostring Technologies) and YS (anti-Salmonella O-antigen conjugated to AlexaFluor647, NB110-16952AF647, Novus Biologicals), as well as nuclear staining with GeoMx Nuclear Stain Morphology kit (121300303, Nanostring Technologies). The processed slide was loaded onto GeoMx-DSP instrument for selection of ROI equivalent to those chosen for HS quantification, and also avoiding areas with CD45+ infiltrating immune cells and PanCK staining (assumed to be mouse-derived stromal cells). For each ROI, probes were collected by the instrument and transferred to a PCR plate for library prep using Seq Code and primers (GeoMx Seq Code Pack, 121400205-121400206, Nanostring Technologies). The libraries from each ROI were pooled, purified by AMPure XP beads (A63880 Beckman Coulter), and resuspended in a volume of elution buffer proportional to the number of pooled ROIs. Libraries were assessed using an Agilent TapeStation, then diluted to 1.6 pmol/L and sequenced (paired - end 2 x27) on Illumina NextSeq2000. Fastq files were uploaded to the GeoMxNGSPipeline application to convert Fastq file to DCC file. The DCC files were transferred back to GeoMx-DSP instrument, which will be associated with the corresponding ROIs. The data were initially analyzed using GeoMx-DSP software. After the quality control of ROI segments and probes, the data were filtered following the DSP default setting. Q3 normalized data was used for further analysis. Statistical significance in gene expression between all three groups (no YS, YS-EGFP, and YS-HepIII) was analyzed using a two-way ANOVA followed by Tukey’s post-hoc test for multiple comparisons.

#### Statistical Overrepresentation Analysis

Statistically significant overrepresentation of genes in Biological Processes, Protein Class, and Reactome Pathways was analyzed using the PANTHER (version 18.0, https://pantherdb.org/, RRID:SCR_004869) overrepresentation test (Fisher test with correction by false discovery rate, Benjamini-Hochberg procedure).

### Untargeted Metabolomics

#### Sample Preparation and Untargeted Metabolomics

Tumor tissue was pulverized using a MultiSample BioPulverizer (BioSpec). Approximately 15 ± 3 mg of tissue was extracted with 300 µL of methanol:acetonitrile:water (2:1:1, v/v/v) containing isotopically labeled standards (d4-citric acid, d8-valine, ¹³C₆-phenylalanine, ¹³C₆-adipic acid, d4-succinic acid) as previously described{Zhang, 2023 #99}. Samples were randomized, and analyzed by untargeted metabolomics using an Ultimate HPG 3400 RS UPLC coupled to an Orbitrap Fusion Lumos mass spectrometer (Thermo) (PMID: 37658085). System suitability was verified with IMS-SR plasma standards (retention time <5% RSD, area <25%). A pooled QC, prepared by combining aliquots from all samples, was injected before and after the batch and every five samples for retention time correction and assessment of analytical variation. Data were processed in Compound Discoverer 3.2, with metabolite identification using HMDB, KEGG, LipidMaps, and MzCloud, applying intensity thresholds of 100K/70K (RP positive/negative) and 70K/60K (HILIC positive/negative). Metabolites with <25% RSD in pooled QC were retained for downstream analysis. Unique IDs were assigned to each compound detected in each chromatographic mode, and the data from all four chromatographies were merged for statistical analysis.

#### Statistics

Untargeted metabolomics data were normalized using variance stabilizing normalization. ANOVA was performed to determine differentially abundant metabolites between the three groups: no YS, YS-EGFP induced, and YS-HepII induced, followed by a Tukey post-hoc test for multiple comparisons to examine specific pairwise comparisons. Metabolites with adjusted p<0.05 were considered to be significantly different. Only annotated endogenous metabolites were used for data representation. Principal component analyses were run on all discovered metabolites using the ClustVis(*123*) webtool (https://biit.cs.ut.ee/clustvis/).

### Untargeted Lipidomics

#### Sample Preparation

Twenty-five mg of tissue samples were homogenized in 300 µL cold PBS using hand homogenizer, and lipids were extracted using the Maytash method (MTBE : methanol : water) spiked with Splash Lipidomix MS standard (Avantilipids, SKU 330707). The lipid extracts were dried under argon drier and stored at −80°C until data acquisition. Equal aliquots of lipid extract from each sample were pooled to create a sample/matrix pool QC (PoolQC). For data acquisition samples were reconstituted in 150 µL solvent (5 mM ammonium formate, 0.1% formic acid, 50% acetonitrile, 50% water).

#### Untargeted Lipidomics Data Acquisition

Samples were randomized prior to data acquisition. An in-house plasma lipid extract was used as a system suitability matrix (SSQC) to monitor instrument performance by determining the area and retention times of lipid analytes before and at the end of the acquisition batch. Pool QC was also used to assess instrument performance before, during, and after the data acquisition batch and was used for downstream data analysis. Lipids were separated on an Acclaim C30 column (2.1 mm × 150 mm) using an Ultimate HPG-3400 RS UPLC and analyzed on an Orbitrap Fusion Lumos mass spectrometer (Thermo) in positive and negative modes using a data-dependent acquisition approach as described previously{Liefwalker, 2021 #100}.

#### Data Analysis

SSQC performance was stable before and after the acquisition batch, with lipid analyte areas within 25% RSD and retention time areas within ±2 min. Pool QC and sample raw data were processed using LipidSearch 5.1 for lipid annotation and relative abundance determination, followed by retention time alignment and merging of positive and negative mode data files(*124*). Lipid annotations with grades A, B, and C, along with optimal adduct ions and ionization modes (positive or negative), were selected. Lipids from Pool QC with area under 25% CV were retained for downstream analysis. One-way ANOVA followed by Tukey’s test for multiple comparisons was performed to identify lipids significantly different between three pairwise comparisons. Lipids with an adjusted p-value < 0.05 were considered differentially abundant.

### Graph Statistics

Error bars in line, dot, or bar graphs were created and statistics analyzed using Prism 10.6.0 (GraphPad, Boston, MA, USA, RRID:SCR_021836). Error bars represent standard error of the mean. All graph p-values were obtained from a one-way ANOVA followed by Tukey’s post-hoc test for multiple comparisons, unless otherwise stated.

## Acknowledgments

This research includes work conducted in the Integrated Mass Spectrometry, Molecular Pathology, Animal Resource Center, and Light Microscopy Imaging Shared Resources, supported by the National Cancer Institute of the NIH under grant number P30CA033572. R01 CA266472, R01 CA272732, R21 CA256593, and R21 CA293969. The content is solely the responsibility of the authors and does not necessarily represent the official views of the National Institutes of Health.

## Author contributions

Conceptualization: NDE, ERM

Methodology: NDE, KD, NH, KVP, PP, ERM

Investigation: NDE, SL, LCA, KD, NH, KVP, PP, ERM

Visualization: NDE

Supervision: NDE, PP, ERM

Writing—original draft: NDE, SL, LCA, ERM

Writing—review & editing: NDE, SL, LCA, KD, NH, KVP, PP, ERM

## Use of Generative AI

No generative AI was used in any part of this manuscript or data acquisition.

## Data and materials availability

All data are available in the main text or the supplementary materials. Raw data can be accessed through the following databases and accession numbers:

GeoMx WTA Raw Data: Available through GEO Datasets (NCBI) Study# GSE314835

Metabolomics Raw Data: Waiting for Identifiers

Lipidomics Raw Data: Waiting for Identifiers

