## Supplemental Files for "Precision Targeting of Heparan Sulfate Proteoglycans Broadly Dysregulates Survival Signaling and Metabolic Adaptation in Pancreatic Cancer"

Nancy Danielle Ebelt *et al.*

**This PDF file includes:**

Figs. S1 to S3

Tables S1 to S5

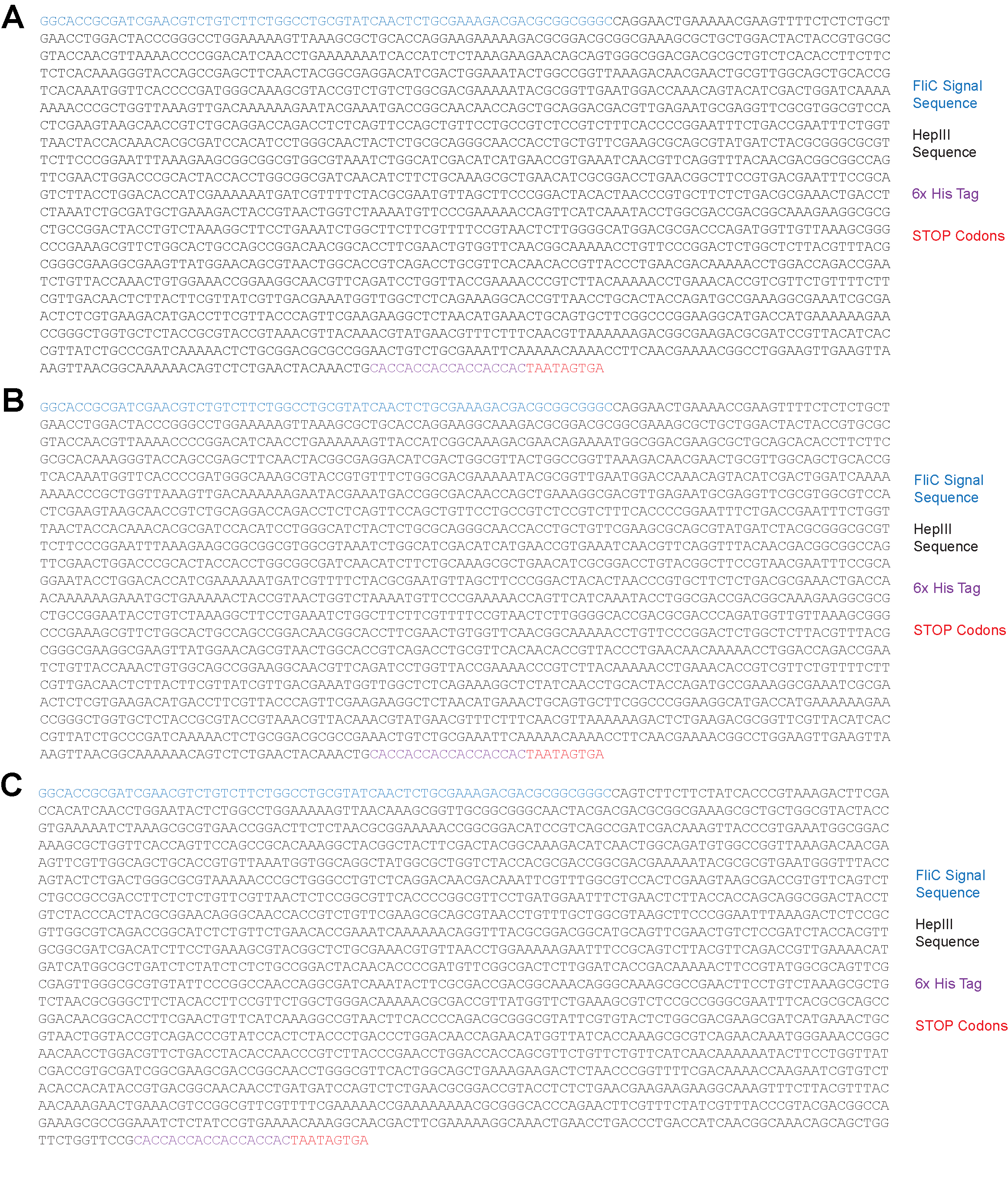

Fig. S1.

**HepIII Transgene Sequences for Expression in YS.** HepIII sequences (black) from *Bacteroides eggerthii* (**A**), *Bacteroides stercoris* (**B**), and *Flavobacterium heparinum* (also known as *Pedobacter heparinus*) (**C**), were modified to include the flagellin signal sequence from *Salmonella typhimurium* (FliC, blue), a 6x His Tag (purple), and three stop codons (red).

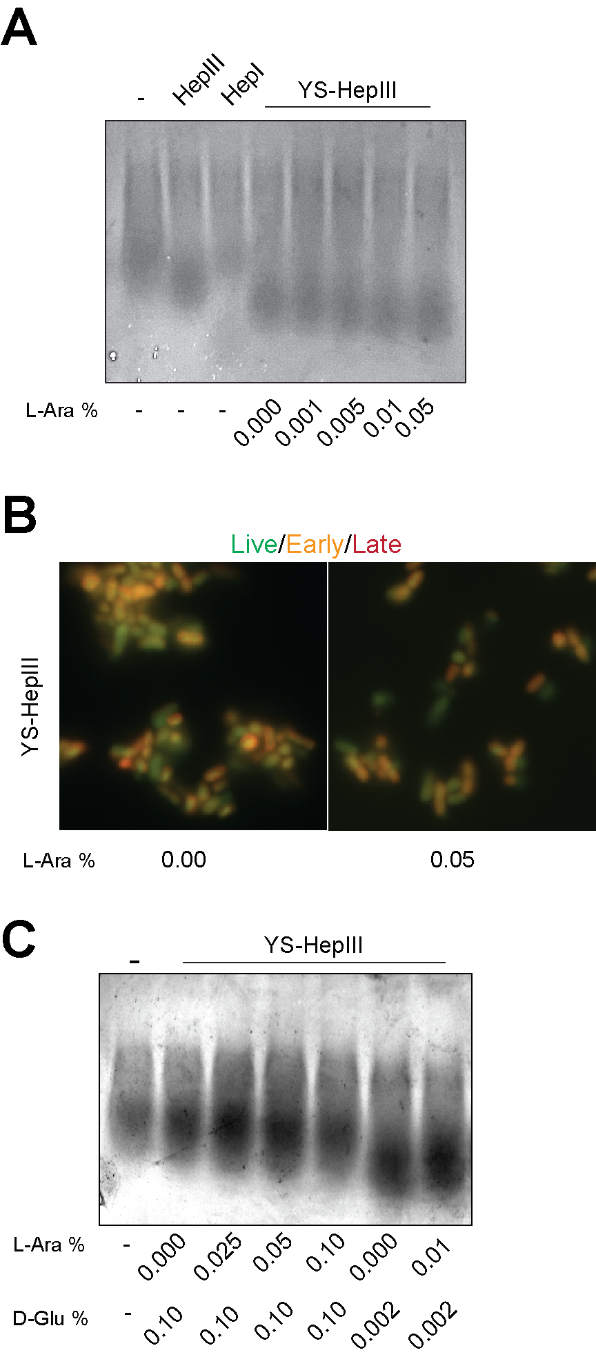

Fig. S2.

**Leaky expression of HepIII from pBAD is Controlled by Glucose.** (**A**) Purified HepIII or HepI enzyme (5mU/mL) or YS-HepIII or HepI enzyme (5mU/mL)YS-HepIII induced with varying levels of L-arabinose was co-incubated with purified HS. After overnight degradation, supernatants were electrophoresed on acrylamide gels to visualize HS degradation compared to undigested HS and purified HepIII enzyme from *F. heparinum.* Gels were stained with Alcian blue to visualize HS smears. (**B**) YS-HepIII uninduced or induced with 0.05% L-arabinose for one hour were stained with acridine orange (live, green) and ethidium bromide (dead, orange) (1:1) and fluorescence was imaged at 100x magnification. (**C**) YS-HepIII was induced with varying concentrations of L-arabinose in the presence of varying concentrations of D-Glucose for 1 hour. Cultures were then co-incubated with HS for 1hr. Supernatants containing HS were run on acrylamide gels to assess degradation. Alcian blue staining was used to visualize HS smears.

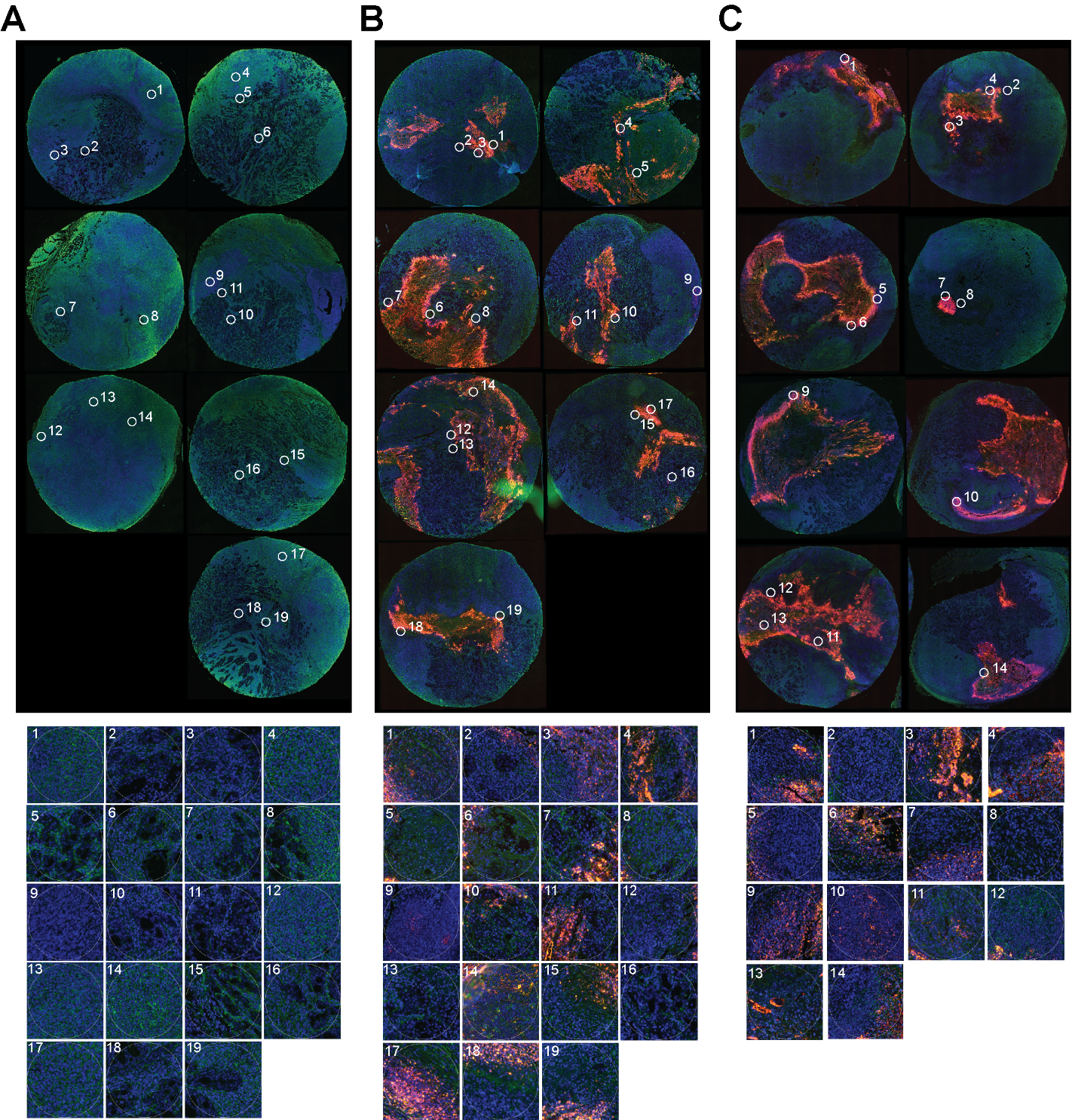

Fig. S3.

**Regions of Interest (ROI) Chosen for HS Degradation Analysis and Spatial Transcriptomics.** Established HPAC xenograft tumors in NSG mice were treated for three consecutive days with PBS (no YS) (**A**), 2.5x10^6^ cfu/mouse YS-EGFP (**B**), or 2.5x10^6^ cfu/mouse YS-HepIII (**C**). Forty-eight hours after the third treatment, YS-EGFP and YS-HepIII-treated mice were administered 40mg/mouse L-arabinose to induce EGFP or HepIII expression. One day after induction with L-arabinose, tumors were excised, fixed, and paraffin embedded. Tumor sections were probed for the presence of YS (red) and heparan sulfate (green), and nuclei were counterstained with DAPI. ROI containing tumor cells near fronts of invading YS were chosen for HS degradation analysis. Serial sections were stained for YS, Pan-Cytokeratin (PanCK), and CD45 and loaded onto the GeoMx Digital Spatial Profiler for spatial transcriptomic ROI selection. Equivalent ROI were chosen and further refined to avoid areas of PanCK or CD45+ mouse cell infiltration. Individual tumors for each treatment are shown with magnified views of ROIs below.

Table S1.

**Genes from Spatial Transcriptomics Analysis Significantly Altered by the Presence of YS Alone.** Full ranked list of differentially expressed genes that were significant between no YS and YS-EGFP-treated mice, and no YS and YS-HepIII-treated mice, but not significant between YS-EGFP and YS-HepIII-treated mice bearing HPAC tumors. Significance was determined by comparing averaged YS-EGFP and YS-HepIII values compared to no YS values. Gene rank values represent fold change divided by adjusted p-values.

| **Gene Name** | **Rank** |
| --- | --- |
| MMP9 | 6.288 |
| NFKBIA | 3.752 |
| ELN | 2.459 |
| SGCA | 2.313 |
| CHRFAM7A | 2.002 |
| GPR4 | 1.518 |
| ELOVL5 | 0.965 |
| GRINA | -0.821 |
| ETV4 | -1.510 |
| ADAR | -1.517 |
| BBLN | -1.520 |
| CSK | -1.564 |
| G6PD | -1.641 |
| DBNL | -1.657 |
| SH3KBP1 | -1.668 |
| KLHL11 | -1.673 |
| BCL2L11 | -1.676 |
| BLOC1S1 | -1.700 |
| PITX1 | -1.712 |
| ACVR1 | -1.727 |
| ZBTB41 | -1.742 |
| VPS25 | -1.749 |
| ITFG2 | -1.769 |
| FOSL2 | -1.798 |
| MUC5AC | -1.815 |
| ETS1 | -1.876 |
| ING1 | -1.929 |
| MDK | -1.930 |
| CDK2AP2 | -1.930 |
| UTP11 | -1.932 |
| ITPR3 | -1.945 |
| DCK | -1.949 |
| ZNF385A | -1.954 |
| MAP2K7 | -2.015 |
| ARID2 | -2.018 |
| GABBR1 | -2.018 |
| IFT57 | -2.032 |
| TAF13 | -2.035 |
| RNF126 | -2.052 |
| RBX1 | -2.060 |
| PAFAH1B2 | -2.064 |
| ATAD3A | -2.064 |
| TLK2 | -2.070 |
| NDST2 | -2.097 |
| BLCAP | -2.103 |
| ZZEF1 | -2.106 |
| **Gene Name** | **Rank** |
| PLPP5 | -2.107 |
| TBC1D22A | -2.111 |
| ABL1 | -2.114 |
| ADGRG1 | -2.119 |
| YWHAB | -2.173 |
| AAMP | -2.176 |
| HNRNPH2 | -2.181 |
| ZFAND6 | -2.184 |
| IRF2BP2 | -2.197 |
| RNF139 | -2.203 |
| YTHDC1 | -2.204 |
| ZNF140 | -2.217 |
| ERI3 | -2.223 |
| RRP1 | -2.227 |
| RNPEP | -2.230 |
| HOOK3 | -2.230 |
| SIRT2 | -2.236 |
| HERPUD1 | -2.256 |
| DPYSL2 | -2.259 |
| CCDC47 | -2.283 |
| FOXJ1 | -2.292 |
| ZNRD2 | -2.309 |
| ZFAND3 | -2.317 |
| TUBA4A | -2.319 |
| SYMPK | -2.320 |
| HDAC1 | -2.332 |
| PRKAG1 | -2.332 |
| SNX13 | -2.333 |
| ROCK2 | -2.334 |
| WAC | -2.346 |
| DPF2 | -2.368 |
| KLC1 | -2.376 |
| PMPCB | -2.400 |
| CAMK2D | -2.432 |
| PPP2R5A | -2.435 |
| BRAF | -2.443 |
| MOB1A | -2.446 |
| FNBP1L | -2.452 |
| ATRN | -2.454 |
| DUX4 | -2.470 |
| SEPTIN7 | -2.473 |
| SHOC2 | -2.473 |
| USO1 | -2.475 |
| HINT1 | -2.486 |
| PAN3 | -2.508 |
| ABCF2 | -2.549 |
| **Gene Name** | **Rank** |
| VPS41 | -2.549 |
| HERC2 | -2.553 |
| AHCTF1 | -2.553 |
| CCNL1 | -2.562 |
| TLK1 | -2.569 |
| ERBIN | -2.579 |
| ADCY6 | -2.582 |
| IL24 | -2.584 |
| PITPNM1 | -2.591 |
| TSNAX | -2.612 |
| CIC | -2.618 |
| HSD17B8 | -2.626 |
| PNPLA2 | -2.635 |
| RBM39 | -2.637 |
| NSUN5 | -2.640 |
| OAT | -2.657 |
| CAMK2N1 | -2.670 |
| GORASP2 | -2.675 |
| YEATS4 | -2.679 |
| MTCH2 | -2.684 |
| TRIO | -2.684 |
| ACSL4 | -2.697 |
| UGDH | -2.703 |
| MIF4GD | -2.712 |
| RBM4 | -2.732 |
| TM9SF1 | -2.735 |
| REX1BD | -2.759 |
| BCL7C | -2.766 |
| COX5A | -2.770 |
| EML4 | -2.771 |
| QRICH1 | -2.774 |
| ZNF391 | -2.784 |
| COPS8 | -2.795 |
| SNU13 | -2.800 |
| E2F4 | -2.802 |
| PTBP2 | -2.808 |
| TMA7 | -2.811 |
| ILRUN | -2.838 |
| GABARAPL1 | -2.840 |
| ERF | -2.846 |
| FHL2 | -2.848 |
| R3HDM2 | -2.856 |
| NCEH1 | -2.866 |
| LRATD2 | -2.872 |
| HSPA4 | -2.902 |
| **Gene Name** | **Rank** |
| CAMK1 | -2.908 |
| DNAJB6 | -2.913 |
| COQ2 | -2.915 |
| CWC15 | -2.942 |
| BCKDHA | -2.944 |
| NDUFS3 | -2.946 |
| GCLM | -2.951 |
| PBRM1 | -2.954 |
| ARL6IP1 | -2.956 |
| VTI1B | -2.960 |
| DAPK3 | -2.964 |
| DDX39B | -2.969 |
| RUNX1 | -2.976 |
| LSM3 | -2.976 |
| CD109 | -2.976 |
| MAP3K3 | -2.989 |
| MACO1 | -3.008 |
| CLK2 | -3.011 |
| SMNDC1 | -3.016 |
| OLA1 | -3.019 |
| NR2F1 | -3.020 |
| RNF4 | -3.025 |
| MYEF2 | -3.033 |
| CNOT10 | -3.041 |
| CNOT11 | -3.052 |
| FNBP4 | -3.060 |
| UQCRC1 | -3.072 |
| UQCRQ | -3.079 |
| MAP2K3 | -3.085 |
| CRADD | -3.099 |
| HOXC4 | -3.102 |
| ROMO1 | -3.104 |
| RFK | -3.111 |
| ZNF787 | -3.116 |
| FAM98A | -3.133 |
| SF1 | -3.177 |
| CBX4 | -3.180 |
| GPSM2 | -3.198 |
| RHEB | -3.198 |
| CPEB2 | -3.204 |
| NTN1 | -3.210 |
| DEPDC5 | -3.225 |
| POSTN | -3.228 |
| WASL | -3.234 |
| REEP4 | -3.235 |
| STUB1 | -3.253 |
| **Gene Name** | **Rank** |
| MEF2D | -3.259 |
| TAGLN3 | -3.297 |
| SYVN1 | -3.314 |
| NBL1 | -3.317 |
| SMAD5 | -3.319 |
| PSPC1 | -3.328 |
| NT5C2 | -3.341 |
| RCN1 | -3.342 |
| AP2A1 | -3.349 |
| BRWD1 | -3.353 |
| CNOT6 | -3.359 |
| CIRBP | -3.361 |
| AGAP3 | -3.373 |
| THAP4 | -3.386 |
| H4C5 | -3.390 |
| BCLAF1 | -3.391 |
| HMGN1 | -3.409 |
| LSM14A | -3.412 |
| DDX50 | -3.431 |
| SYNRG | -3.433 |
| PDZD8 | -3.458 |
| KIF1B | -3.482 |
| DDX4 | -3.488 |
| CD46 | -3.509 |
| STK24 | -3.533 |
| CYRIB | -3.545 |
| MAPKBP1 | -3.551 |
| CELF1 | -3.554 |
| MGST1 | -3.559 |
| FAM3C | -3.562 |
| VAT1 | -3.571 |
| IGHG2 | -3.574 |
| MYLK | -3.605 |
| AP1B1 | -3.611 |
| H2BC5 | -3.620 |
| MAST4 | -3.622 |
| PRPF38B | -3.628 |
| KLHDC3 | -3.633 |
| POLR2I | -3.659 |
| PUM1 | -3.668 |
| CBFB | -3.686 |
| INAFM2 | -3.698 |
| MEPCE | -3.710 |
| PRELID3B | -3.712 |
| EPB41L1 | -3.722 |
| MDH1 | -3.731 |
| ZNF622 | -3.732 |
| TPR | -3.733 |
| TIAL1 | -3.733 |
| PFDN5 | -3.744 |
| ATG3 | -3.749 |
| RARA | -3.766 |
| CDC5L | -3.767 |
| TNPO3 | -3.800 |
| PRRC1 | -3.808 |
| HOXA3 | -3.812 |
| **Gene Name** | **Rank** |
| CTNNA1 | -3.817 |
| ST14 | -3.817 |
| B4GALT5 | -3.831 |
| FEM1B | -3.836 |
| DNAJA1 | -3.843 |
| KIAA1522 | -3.858 |
| YPEL5 | -3.877 |
| OCIAD1 | -3.894 |
| COP1 | -3.904 |
| GPC1 | -3.906 |
| NDUFS6 | -3.925 |
| CCNL2 | -3.927 |
| UQCRC2 | -3.946 |
| MKLN1 | -3.947 |
| ELOVL1 | -3.949 |
| YY1 | -3.951 |
| MLF2 | -3.969 |
| FOXM1 | -3.972 |
| RAB14 | -3.992 |
| ZFP36L1 | -4.001 |
| ATP8B2 | -4.007 |
| AGPS | -4.008 |
| MLLT10 | -4.009 |
| JPH4 | -4.028 |
| CDC37 | -4.053 |
| MCL1 | -4.055 |
| PRKCA | -4.060 |
| PGLS | -4.066 |
| KCMF1 | -4.107 |
| PPP2R5C | -4.109 |
| AKIRIN2 | -4.118 |
| UFC1 | -4.118 |
| AIG1 | -4.131 |
| H2BC8 | -4.134 |
| EBNA1BP2 | -4.149 |
| EML2 | -4.164 |
| TMPO | -4.179 |
| PFDN2 | -4.187 |
| XRCC6 | -4.200 |
| PPP4C | -4.205 |
| H3C15 | -4.213 |
| OGT | -4.240 |
| KLF4 | -4.242 |
| MATN2 | -4.252 |
| ASH2L | -4.258 |
| CD151 | -4.264 |
| FXR2 | -4.290 |
| TCP1 | -4.305 |
| LUC7L | -4.323 |
| GNB4 | -4.359 |
| ATP5F1B | -4.362 |
| CDK9 | -4.405 |
| GIT1 | -4.409 |
| RTN4 | -4.413 |
| VLDLR | -4.417 |
| STK25 | -4.423 |
| **Gene Name** | **Rank** |
| SBNO1 | -4.439 |
| GRK2 | -4.455 |
| KMT2D | -4.471 |
| EBAG9 | -4.481 |
| TOMM40 | -4.490 |
| ATP5F1A | -4.491 |
| TMEM138 | -4.497 |
| CUL1 | -4.512 |
| LASP1 | -4.536 |
| DDX39A | -4.554 |
| INSR | -4.570 |
| RAB43 | -4.591 |
| LAP3 | -4.600 |
| EFHD2 | -4.610 |
| MSLN | -4.642 |
| PCDH19 | -4.651 |
| LRCH1 | -4.658 |
| DGCR2 | -4.672 |
| PRKACA | -4.676 |
| EML3 | -4.680 |
| BHLHE41 | -4.694 |
| IMPDH2 | -4.699 |
| CSNK1G3 | -4.703 |
| LSR | -4.721 |
| NCL | -4.726 |
| TNNI2 | -4.740 |
| HMGN3 | -4.744 |
| DYRK1B | -4.755 |
| MUC16 | -4.785 |
| FBL | -4.793 |
| NF2 | -4.794 |
| GRB2 | -4.800 |
| PLEKHA5 | -4.800 |
| HEXA | -4.810 |
| MRPL45 | -4.843 |
| LOXL1 | -4.843 |
| IPO13 | -4.847 |
| PRPF39 | -4.855 |
| TAOK1 | -4.914 |
| U2SURP | -4.949 |
| TRAPPC14 | -4.970 |
| SRSF10 | -4.973 |
| EIF4A2 | -4.974 |
| EFR3A | -4.982 |
| H4C8 | -4.990 |
| SMO | -4.999 |
| FBXW7 | -5.024 |
| UBE2B | -5.025 |
| LGALS3BP | -5.057 |
| CDH1 | -5.066 |
| SP3 | -5.068 |
| SKAP2 | -5.088 |
| SETX | -5.099 |
| TNFAIP1 | -5.111 |
| KANSL1 | -5.115 |
| SEC11C | -5.134 |
| **Gene Name** | **Rank** |
| ATN1 | -5.135 |
| RND3 | -5.138 |
| TBL1XR1 | -5.180 |
| ITGAV | -5.204 |
| SCAF8 | -5.211 |
| MRPL51 | -5.222 |
| ECPAS | -5.270 |
| ATAD1 | -5.303 |
| TAF2 | -5.321 |
| NR3C1 | -5.324 |
| PCM1 | -5.363 |
| IK | -5.365 |
| UBE2G1 | -5.369 |
| FNDC3B | -5.392 |
| CDK13 | -5.399 |
| H1-2 | -5.443 |
| B2M | -5.489 |
| NPDC1 | -5.495 |
| RAD21 | -5.504 |
| CAPN1 | -5.509 |
| ARL15 | -5.517 |
| GSK3A | -5.555 |
| ZFP91 | -5.568 |
| RPS6KA3 | -5.584 |
| NET1 | -5.587 |
| CHCHD1 | -5.599 |
| POLR2E | -5.612 |
| DHX16 | -5.618 |
| PIAS2 | -5.638 |
| PTGES | -5.643 |
| H2BS1 | -5.644 |
| ELF3 | -5.655 |
| TSPAN13 | -5.684 |
| CSNK1E | -5.736 |
| RPL4 | -5.765 |
| PSMD6 | -5.770 |
| ALPP | -5.831 |
| SPAG9 | -5.853 |
| PHF14 | -5.896 |
| AMD1 | -5.902 |
| STRBP | -5.930 |
| OTUD7B | -5.976 |
| DDX17 | -6.027 |
| STK38L | -6.030 |
| AMMECR1L | -6.055 |
| COX7A2L | -6.090 |
| EP300 | -6.092 |
| RBM26 | -6.130 |
| HSPA8 | -6.134 |
| PIK3R1 | -6.150 |
| ITM2B | -6.163 |
| RGS7BP | -6.168 |
| HLA-DRB1 | -6.175 |
| SCNN1A | -6.183 |
| ANO1 | -6.203 |
| INPPL1 | -6.224 |
| **Gene Name** | **Rank** |
| RAD23B | -6.225 |
| UBE4B | -6.229 |
| CUX1 | -6.240 |
| STAG1 | -6.244 |
| UBE2W | -6.244 |
| GSTP1 | -6.247 |
| CAV2 | -6.280 |
| SMARCB1 | -6.283 |
| SMAP1 | -6.285 |
| RDH10 | -6.304 |
| TCIRG1 | -6.318 |
| LRRC41 | -6.347 |
| DEK | -6.359 |
| NSUN2 | -6.436 |
| SELENOT | -6.450 |
| SNX17 | -6.461 |
| IPO7 | -6.492 |
| NUCKS1 | -6.501 |
| PDLIM1 | -6.540 |
| BCL2 | -6.553 |
| TULP4 | -6.555 |
| IMMT | -6.573 |
| MAZ | -6.585 |
| PPM1A | -6.590 |
| MAP3K7 | -6.609 |
| PPP4R3A | -6.619 |
| ATP6AP2 | -6.644 |
| FAM204A | -6.661 |
| SH3GLB2 | -6.661 |
| NONO | -6.730 |
| SLC25A46 | -6.736 |
| ELF1 | -6.759 |
| OAZ1 | -6.779 |
| ST3GAL4 | -6.795 |
| THRA | -6.813 |
| UNC93B1 | -6.813 |
| EMG1 | -6.850 |
| H4C15 | -6.860 |
| ASS1 | -6.862 |
| B4GALT1 | -6.871 |
| CLINT1 | -6.907 |
| YBX3 | -6.910 |
| KCTD6 | -6.936 |
| FOXO3 | -6.951 |
| RANBP1 | -6.954 |
| ATXN2 | -6.980 |
| KAT6A | -6.988 |
| S100P | -7.026 |
| SNW1 | -7.055 |
| TET3 | -7.115 |
| FAM172A | -7.128 |
| LARP4B | -7.145 |
| TMED10 | -7.244 |
| MED15 | -7.252 |
| VAPB | -7.293 |
| TRPS1 | -7.405 |
| **Gene Name** | **Rank** |
| PSMD3 | -7.417 |
| SCHIP1 | -7.478 |
| ARF3 | -7.484 |
| PDGFRA | -7.498 |
| VPS28 | -7.518 |
| GET3 | -7.611 |
| MAT2B | -7.617 |
| TMPRSS4 | -7.620 |
| GTPBP2 | -7.739 |
| FYTTD1 | -7.830 |
| IP6K2 | -7.880 |
| MYH14 | -7.890 |
| BMPR2 | -7.897 |
| EIF4A3 | -7.939 |
| TSC22D1 | -7.981 |
| RARG | -8.079 |
| MLLT6 | -8.137 |
| NFIC | -8.143 |
| MCM5 | -8.156 |
| RAD23A | -8.175 |
| TRIM33 | -8.198 |
| TFF3 | -8.198 |
| NHP2 | -8.222 |
| EIF4EBP1 | -8.236 |
| SRGAP2B | -8.255 |
| NBR1 | -8.264 |
| FXR1 | -8.284 |
| SNRPF | -8.310 |
| ZBTB44 | -8.364 |
| AGO2 | -8.365 |
| ANXA7 | -8.393 |
| BAX | -8.398 |
| NCKAP1 | -8.399 |
| PAIP1 | -8.431 |
| YTHDF2 | -8.517 |
| INTS6 | -8.638 |
| EXT1 | -8.656 |
| EPS8L2 | -8.717 |
| PRPS1 | -8.729 |
| CHCHD7 | -8.744 |
| OTUB1 | -8.773 |
| SNRPE | -8.777 |
| CHIC2 | -8.785 |
| MBTPS1 | -8.812 |
| GTF2I | -8.855 |
| PCIF1 | -8.856 |
| PPA1 | -8.874 |
| H3C10 | -8.888 |
| GPAT4 | -8.897 |
| PEAK1 | -8.902 |
| RGL1 | -8.907 |
| TRA2A | -8.924 |
| SMC1A | -8.938 |
| SMDT1 | -9.095 |
| ZFP36L2 | -9.258 |
| RPLP0 | -9.281 |
| **Gene Name** | **Rank** |
| PADI1 | -9.286 |
| PLEKHO1 | -9.385 |
| SUMO2 | -9.422 |
| H2AX | -9.445 |
| CNOT9 | -9.494 |
| AP2S1 | -9.509 |
| NANS | -9.559 |
| SPATS2L | -9.615 |
| RPS26 | -9.677 |
| SF3B1 | -9.700 |
| KDM5C | -9.742 |
| KRT4 | -9.790 |
| UQCC2 | -9.813 |
| ETV3 | -9.820 |
| UBE2L3 | -9.833 |
| STT3B | -9.859 |
| POLR2G | -9.926 |
| RBMS1 | -9.952 |
| NACC2 | -9.979 |
| RAB11A | -10.112 |
| EXOSC8 | -10.130 |
| HAT1 | -10.137 |
| KAT5 | -10.151 |
| PHF20L1 | -10.174 |
| MTA2 | -10.206 |
| MTCH1 | -10.305 |
| HM13 | -10.317 |
| TMEM106C | -10.495 |
| FRYL | -10.529 |
| CYCS | -10.606 |
| PLD3 | -10.665 |
| CDC73 | -10.735 |
| RNF11 | -10.750 |
| MANF | -10.915 |
| WIZ | -10.932 |
| BPTF | -10.983 |
| EPRS1 | -11.008 |
| YIPF3 | -11.013 |
| VPS35 | -11.064 |
| EGLN2 | -11.077 |
| MBNL2 | -11.173 |
| DYNLL2 | -11.177 |
| CNOT2 | -11.177 |
| ILK | -11.296 |
| HGS | -11.317 |
| NME2 | -11.334 |
| ZFAND5 | -11.349 |
| CCT7 | -11.370 |
| WDR43 | -11.427 |
| CLTA | -11.450 |
| SRRM2 | -11.502 |
| CSNK2A2 | -11.502 |
| ARHGEF1 | -11.679 |
| SF3B2 | -11.685 |
| NDUFB10 | -11.686 |
| PSMC4 | -11.726 |
| **Gene Name** | **Rank** |
| SLC12A2 | -11.727 |
| SYNCRIP | -11.794 |
| RPL21 | -11.929 |
| IER3IP1 | -11.973 |
| UBL5 | -11.976 |
| ATP2C1 | -11.996 |
| PPL | -12.020 |
| PGRMC2 | -12.040 |
| UBE2Z | -12.051 |
| PSMA2 | -12.084 |
| EPHA7 | -12.151 |
| MID1IP1 | -12.230 |
| KMT2E | -12.299 |
| DHCR24 | -12.321 |
| TMEM258 | -12.383 |
| MPC1 | -12.518 |
| UBXN4 | -12.539 |
| UBR5 | -12.578 |
| MED22 | -12.669 |
| RAP2A | -12.705 |
| RBM14 | -12.719 |
| RPL22L1 | -12.818 |
| ITGA6 | -12.892 |
| DDX47 | -12.915 |
| UBAP2L | -12.924 |
| UQCRHL | -13.035 |
| CDK16 | -13.064 |
| SS18 | -13.070 |
| RAB7A | -13.140 |
| USP15 | -13.240 |
| ANKRD13C | -13.258 |
| ARF1 | -13.312 |
| ACTA2 | -13.365 |
| MAP4K4 | -13.394 |
| MARCHF5 | -13.448 |
| APOBEC3A_B | -13.519 |
| MIB1 | -13.563 |
| NAB1 | -13.576 |
| GDE1 | -13.600 |
| NPM1 | -13.619 |
| TRIM28 | -13.740 |
| TMEM30A | -13.881 |
| ABHD2 | -13.882 |
| GIT2 | -14.070 |
| CAPRIN1 | -14.117 |
| PDCD5 | -14.165 |
| SUGT1 | -14.180 |
| GTF2H1 | -14.278 |
| ALDH1A1 | -14.345 |
| ARPC4 | -14.381 |
| CLTB | -14.414 |
| XRN2 | -14.473 |
| EPN1 | -14.479 |
| SPTBN1 | -14.622 |
| RPS6KB1 | -14.668 |
| **Gene Name** | **Rank** |
| UBE3A | -14.683 |
| ARPC2 | -14.778 |
| C1QBP | -14.861 |
| CYFIP1 | -14.894 |
| MTA1 | -14.905 |
| DAZAP2 | -14.922 |
| SKP1 | -14.967 |
| RPS13 | -14.977 |
| NACA | -15.043 |
| RPL35 | -15.117 |
| SERINC1 | -15.266 |
| MAPK3 | -15.318 |
| PPP3R1 | -15.340 |
| ENSA | -15.355 |
| PCMTD1 | -15.409 |
| EEF1B2 | -15.520 |
| PNISR | -15.537 |
| THRAP3 | -15.558 |
| RIN2 | -15.591 |
| ERGIC1 | -15.668 |
| TSPAN1 | -15.690 |
| RBPJ | -15.751 |
| RPS15A | -15.759 |
| SPCS3 | -15.771 |
| AKAP8L | -15.934 |
| CBX6 | -15.965 |
| KCNN4 | -16.011 |
| PDHA1 | -16.035 |
| SEC61G | -16.096 |
| ANAPC5 | -16.143 |
| CIBAR1 | -16.172 |
| RBM25 | -16.245 |
| BRD4 | -16.333 |
| LGALS3 | -16.483 |
| IST1 | -16.550 |
| MINK1 | -16.667 |
| SARS1 | -16.700 |
| MPRIP | -16.707 |
| TNKS2 | -16.717 |
| HMGB2 | -16.808 |
| PDIA4 | -16.822 |
| TACSTD2 | -16.831 |
| RAB1A | -16.893 |
| CTDNEP1 | -16.980 |
| VKORC1L1 | -17.048 |
| STMN1 | -17.151 |
| MKNK2 | -17.157 |
| BANF1 | -17.162 |
| CNOT1 | -17.210 |
| MYC | -17.553 |
| NFE2L1 | -17.762 |
| EIF4A1 | -17.783 |
| SGPL1 | -17.816 |
| ABCE1 | -17.865 |
| CYC1 | -17.889 |
| NDUFS2 | -17.950 |
| **Gene Name** | **Rank** |
| TUBB4B | -17.972 |
| RPS4X | -18.008 |
| HOXC6 | -18.013 |
| DCN | -18.014 |
| PDE7A | -18.105 |
| KRT5 | -18.148 |
| GOLGA7 | -18.210 |
| KLF13 | -18.225 |
| TSC22D4 | -18.247 |
| TKT | -18.271 |
| SEPTIN2 | -18.279 |
| GABARAP | -18.283 |
| RNASEK | -18.410 |
| RPL13A | -18.440 |
| SCX | -18.534 |
| H2BC18 | -18.620 |
| DCAF8 | -18.678 |
| GALNT1 | -18.768 |
| PDXDC1 | -18.787 |
| ETFA | -18.798 |
| EIF3M | -18.823 |
| DDOST | -18.856 |
| LAMB1 | -18.858 |
| MEDAG | -18.997 |
| XBP1 | -19.000 |
| PPP1CB | -19.018 |
| MARCKS | -19.095 |
| TUSC3 | -19.263 |
| RAB21 | -19.296 |
| TMCO1 | -19.438 |
| PSMA4 | -19.445 |
| TAF15 | -19.532 |
| DDX3X | -19.561 |
| UBE2N | -19.570 |
| KRT7 | -19.570 |
| TM9SF3 | -19.585 |
| NEO1 | -19.616 |
| TOMM22 | -19.659 |
| EGR1 | -19.702 |
| NSD3 | -19.719 |
| BCAP31 | -19.767 |
| LENG8 | -19.777 |
| ARF5 | -19.868 |
| VCP | -19.891 |
| PDIA6 | -19.932 |
| RACK1 | -20.025 |
| SEC63 | -20.041 |
| PAPOLA | -20.067 |
| UBE2V2 | -20.146 |
| CFDP1 | -20.166 |
| MALL | -20.187 |
| CTDSP1 | -20.208 |
| CLIC1 | -20.218 |
| UBE2D2 | -20.226 |
| PLK2 | -20.257 |
| CCT4 | -20.272 |
| **Gene Name** | **Rank** |
| DDRGK1 | -20.277 |
| LY6E | -20.307 |
| BAG5 | -20.308 |
| ATF7 | -20.385 |
| PHLDA3 | -20.428 |
| MUC1 | -20.478 |
| SNRPB2 | -20.502 |
| PRKAA1 | -20.514 |
| TRIP12 | -20.560 |
| TRA2B | -20.587 |
| RAB35 | -20.597 |
| EIF3K | -20.612 |
| DNAJB11 | -20.616 |
| THBS1 | -20.635 |
| SLC39A7 | -20.736 |
| SMARCA5 | -20.742 |
| RAB11B | -20.761 |
| ACTR3 | -20.803 |
| RAB28 | -20.817 |
| ARPC1B | -20.821 |
| WDR33 | -20.822 |
| PHIP | -20.833 |
| TMBIM6 | -20.941 |
| CDK4 | -20.959 |
| EZR | -20.974 |
| AGR2 | -20.986 |
| PPP2CA | -21.001 |
| PRDX1 | -21.072 |
| EIF5 | -21.099 |
| MSI2 | -21.113 |
| EIF4B | -21.149 |
| EIF4G3 | -21.168 |
| NFYA | -21.174 |
| H3-3A | -21.223 |
| DYNC1H1 | -21.260 |
| ANKRD17 | -21.277 |
| PTP4A1 | -21.296 |
| GLYR1 | -21.312 |
| ZSWIM8 | -21.353 |
| CCDC9 | -21.383 |
| CP | -21.468 |
| YWHAE | -21.493 |
| HPRT1 | -21.535 |
| MSL1 | -21.576 |
| ZRANB2 | -21.589 |
| BTG1 | -21.625 |
| SON | -21.634 |
| RTN3 | -21.807 |
| RBM42 | -21.823 |
| FKBP4 | -21.850 |
| CDC42 | -21.986 |
| CDK11A | -22.047 |
| RAB5C | -22.117 |
| SRP68 | -22.121 |
| CRIP1 | -22.279 |
| PSMD4 | -22.310 |
| **Gene Name** | **Rank** |
| SUPT6H | -22.340 |
| HUWE1 | -22.388 |
| CCNI | -22.429 |
| PKN2 | -22.450 |
| STIP1 | -22.467 |
| SEC62 | -22.509 |
| PSMD2 | -22.548 |
| UBA52 | -22.558 |
| CALU | -22.561 |
| ZNF703 | -22.578 |
| EIF3D | -22.641 |
| GNAI2 | -22.649 |
| CSNK2B | -22.650 |
| HTRA1 | -22.666 |
| ARFIP2 | -22.687 |
| BRD2 | -22.712 |
| MARCHF6 | -22.806 |
| RAP2B | -22.814 |
| PTCH1 | -22.845 |
| CUL3 | -22.858 |
| XPO7 | -22.876 |
| PNN | -22.900 |
| SRSF3 | -22.914 |
| CCND1 | -22.944 |
| CAPG | -22.953 |
| CSNK2A3 | -23.030 |
| PRMT1 | -23.067 |
| PSMB7 | -23.138 |
| CLPTM1 | -23.160 |
| YBX1 | -23.185 |
| SF3B4 | -23.187 |
| PURA | -23.213 |
| MGST2 | -23.301 |
| CLTC | -23.308 |
| H2BC4 | -23.312 |
| PSMD13 | -23.347 |
| NDUFS4 | -23.422 |
| FARP1 | -23.423 |
| PRDX2 | -23.459 |
| RAB2A | -23.466 |
| PTP4A2 | -23.609 |
| ARL6IP4 | -23.708 |
| NSD2 | -23.725 |
| PSMA1 | -23.779 |
| ATP5MC3 | -23.784 |
| DYNLL1 | -23.805 |
| EIF3F | -23.819 |
| GARS1 | -23.837 |
| APP | -23.871 |
| UBE2E3 | -23.886 |
| TPM4 | -23.887 |
| PCBP1 | -24.019 |
| EWSR1 | -24.030 |
| APH1A | -24.048 |
| CNOT3 | -24.083 |
| TAF10 | -24.097 |
| **Gene Name** | **Rank** |
| COPS6 | -24.106 |
| CCT6A | -24.129 |
| CEMIP2 | -24.149 |
| MED13 | -24.161 |
| GOLPH3 | -24.183 |
| RANBP9 | -24.189 |
| VEZF1 | -24.211 |
| HNRNPA3 | -24.280 |
| TPM3 | -24.347 |
| NCK1 | -24.383 |
| U2AF2 | -24.451 |
| JOSD1 | -24.495 |
| SKI | -24.579 |
| RTRAF | -24.586 |
| COX6B1 | -24.704 |
| MAX | -24.724 |
| AXL | -24.763 |
| RPN1 | -24.775 |
| GLUL | -24.829 |
| LRP6 | -24.902 |
| HNRNPM | -24.921 |
| PPP1R14B | -24.928 |
| ERH | -25.002 |
| FUS | -25.023 |
| PSMA5 | -25.028 |
| COMMD3 | -25.049 |
| TNFRSF21 | -25.139 |
| EFTUD2 | -25.207 |
| HMGB1 | -25.302 |
| SHC1 | -25.345 |
| HNRNPH1 | -25.361 |
| **Gene Name** | **Rank** |
| EEA1 | -25.371 |
| RBBP7 | -25.391 |
| NUDT21 | -25.400 |
| TMEM59 | -25.407 |
| PIK3CA | -25.412 |
| HMGN2 | -25.463 |
| BTF3 | -25.464 |
| RAN | -25.472 |
| SH3GLB1 | -25.504 |
| RRAGA | -25.550 |
| SPARC | -25.622 |
| PBX1 | -25.676 |
| SLC25A5 | -25.730 |
| SELENOW | -25.750 |
| SYPL1 | -25.792 |
| EZH2 | -25.794 |
| H2AC19 | -25.859 |
| PAIP2 | -25.961 |
| SLC6A6 | -26.046 |
| FKBP3 | -26.123 |
| ATF4 | -26.132 |
| PURB | -26.150 |
| PPIAL4F | -26.153 |
| LTBP4 | -26.242 |
| HNRNPDL | -26.287 |
| SLC35E3 | -26.294 |
| CTTN | -26.337 |
| QKI | -26.367 |
| AZIN1 | -26.385 |
| DDX5 | -26.421 |
| ANXA11 | -26.463 |
| **Gene Name** | **Rank** |
| CHTOP | -26.593 |
| PSMC1 | -26.602 |
| UBE2K | -26.706 |
| C6orf62 | -26.791 |
| NAP1L1 | -26.876 |
| PABPN1 | -27.062 |
| H4C12 | -27.152 |
| MTDH | -27.308 |
| ZNF207 | -27.324 |
| DNAJA2 | -27.343 |
| EIF4E | -27.355 |
| HNRNPU | -27.379 |
| ZNF706 | -27.408 |
| AMFR | -27.450 |
| SUMO1 | -27.551 |
| WLS | -27.638 |
| H2AZ1 | -27.756 |
| MRFAP1 | -27.849 |
| GSN | -27.911 |
| RASA1 | -27.950 |
| MATR3 | -28.014 |
| CAND1 | -28.100 |
| PPP5C | -28.165 |
| CNBP | -28.225 |
| WDR35 | -28.370 |
| CCNA2 | -28.375 |
| FAM102B | -28.550 |
| HMGCS1 | -28.559 |
| DHX15 | -28.617 |
| FOXP1 | -28.636 |
| SFPQ | -28.652 |
| **Gene Name** | **Rank** |
| SERBP1 | -28.687 |
| CSDE1 | -28.878 |
| TFAP2A | -28.931 |
| KPNB1 | -28.948 |
| TNS3 | -29.252 |
| CAPNS1 | -29.381 |
| YWHAZ | -29.456 |
| KLF5 | -29.737 |
| NR1D2 | -29.817 |
| LMO4 | -30.258 |
| HNRNPUL1 | -30.411 |
| HNRNPD | -30.665 |
| NFIX | -30.914 |
| UBTF | -30.933 |
| FUBP1 | -31.076 |
| SRSF5 | -31.485 |
| CBX5 | -31.820 |
| SMG6 | -32.466 |
| SRSF1 | -33.120 |
| CALM2 | -33.885 |
| LYPD2 | -33.951 |
| CPSF6 | -33.980 |
| NR2F2 | -34.242 |
| ANGPT1 | -34.952 |
| PSIP1 | -35.589 |
| FXYD3 | -35.917 |
| TCF4 | -38.076 |
| HLA-DQB1 | -39.407 |

Table S2.

**Genes from Spatial Transcriptomics Analysis Significantly Altered by the Addition of HepIII to YS.** Full ranked list of differentially expressed genes that were significant between all three groups (no YS, YS-EGFP, or YS-HepIII treated mice bearing HPAC tumors) wherein YS-HepIII effects are additive to YS-EGFP effects in the same direction compared to no YS. Significance was determined by comparing YS-HepIII values compared to no YS values. Gene rank values represent fold change divided by adjusted p-values.

| **Gene Name** | **Rank** |
| --- | --- |
| UBALD1 | 27.3513 |
| GJC2 | 7.83527 |
| CERS2 | -1.6959 |
| BMP4 | -5.5551 |
| FTH1 | -6.8698 |
| HSPA1A | -8.1882 |
| ALDOA | -12.642 |
| FXYD5 | -19.227 |
| H3-3B | -19.644 |
| GPX1 | -20.105 |
| ATP1B1 | -20.830 |
| SOX9 | -21.117 |
| FN1 | -22.381 |
| CTSD | -23.309 |
| HDLBP | -23.630 |
| CD63 | -23.937 |
| RPL35A | -24.627 |
| OST4 | -24.742 |
| **Gene Name** | **Rank** |
| EIF5A | -25.0779 |
| ITGA3 | -25.7958 |
| ELOB | -25.8959 |
| HSP90AB1 | -25.9389 |
| TLE5 | -26.084 |
| RPL23 | -26.3424 |
| GNAS | -26.38 |
| UBE2S | -26.7732 |
| RPS7 | -26.9895 |
| RPS29 | -27.161 |
| SND1 | -27.2497 |
| ADAM9 | -27.2633 |
| RPL27 | -27.5176 |
| RPL15 | -27.8951 |
| RPL14 | -28.2981 |
| MYL6 | -28.6529 |
| RPS12 | -29.085 |
| RPL5 | -29.099 |
| **Gene Name** | **Rank** |
| RPS17 | -29.51 |
| MYOF | -29.5707 |
| LMNA | -29.6239 |
| F3 | -29.8256 |
| HNRNPA2B1 | -30.129 |
| EIF4G2 | -30.1915 |
| RPS27 | -30.3968 |
| HSPA9 | -30.5844 |
| RPL37 | -31.0867 |
| RPL34 | -31.3487 |
| PCBP2 | -32.219 |
| H3C13 | -33.0914 |
| RPL7A | -33.5533 |
| RPSA | -33.8371 |
| FLNB | -35.3367 |
| S100A10 | -35.3591 |
| HSP90AA1 | -35.8487 |
| RPL17 | -35.9063 |
| **Gene Name** | **Rank** |
| RPL36A | -37.1571 |
| RPL26 | -37.3572 |
| RPL38 | -37.5663 |
| ITGB4 | -37.7634 |
| H1-5 | -39.5134 |
| RPS5 | -40.8229 |
| CD9 | -41.0048 |
| RPS23 | -41.9778 |
| RPS3A | -42.12 |
| RPL19 | -42.5168 |
| TMSB4X | -43.4495 |
| RPL37A | -43.5239 |
| RPL28 | -43.9508 |
| S100A4 | -44.532 |
| TXN | -48.7331 |
| ANXA1 | -50.4781 |
| KRT13 | -61.7594 |
| KRT8 | -64.9592 |

**Table S3.**

**Genes from Spatial Transcriptomics Analysis Rescued by YS-HepIII Treatment.** Full ranked list of differentially expressed genes that were significant between no YS and YS-EGFP-treated mice, and YS-EGFP and YS-HepIII-treated mice, but not significant between no YS and YS-HepIII-treated mice bearing HPAC tumors. Significance was determined by comparing YS-HepIII values compared to YS-EGFP values. Gene rank values represent fold change divided by adjusted p-values.

| **Gene Name** | **Rank** |
| --- | --- |
| SLC2A1 | 8.294427 |
| LAMC | 24.19208 |
| MZT2B | 4.11012 |
| NDRG1 | 3.037457 |
| LCN2 | 2.83707 |
| PTBP1 | 1.546886 |
| ERO1A | 1.28859 |
| LAMB3 | 1.128121 |
| LGALS1 | 0.567674 |
| MT2A | 0.456578 |
| VEGFA | 0.436809 |
| CD55 | 0.216639 |
| GOLGA8A | -0.06485 |
| H4C1 | -0.42581 |
| BMERB1 | -1.51417 |
| COL1A2 | -17.382 |
| PRMT8 | -22.9166 |

**Table S4**

**Genes from Spatial Transcriptomics Analysis Uniquely Altered by the Presence of YS-HepIII.** Full ranked list of differentially expressed genes that were significant between no YS and YS-HepIII, and YS-EGFP and YS-HepIII, but not significant between no YS and YS-EGFP. Significance was determined by comparing averaged no YS and YS-EGFP values compared to YS-HepIII values. Gene rank values represent fold change divided by adjusted p-values.

| **Gene Name** | **Rank** |
| --- | --- |
| STX1B | 8.818089 |
| WNT9A | 7.892822 |
| PTPRCAP | 5.099282 |
| LDLRAD4 | 3.75124 |
| SLC5A10 | 2.400667 |
| HOXB3 | 2.140134 |
| SLC16A3 | -2.30698 |
| PODXL | -3.07938 |
| MIF | -3.73038 |
| ITGB1 | -5.71712 |
| GPI | -5.81151 |
| HMGA1 | -13.5878 |
| CSTB | -32.1339 |

**Table S5**

**Total Biological Processes, Pathways, and Protein Classes Overrepresented in the Spatial Transcriptomics Gene Set.** The full ranked (fold change/adjusted p-value) list of differentially expressed genes that were significant between YS-HepIII and no YS-treated mice bearing HPAC tumors was uploaded to the PANTHER database for overrepresentation analysis. Terms, number of genes in the reference dataset (total WTA gene set) that are represented in each term, fold enrichment in YS-HepIII compared to no YS-treated tumors, and the adjusted p-values (Fisher’s Exact test with False Discovery Rate) are reported.

| **PANTHER GO-Slim Biological Process** | **REFLIST- Total WTA Gene Set** | **YS-HepIII /noYS Fold Enrichment** | **Adjusted P-Value** |
| --- | --- | --- | --- |
| mitochondrial electron transport, ubiquinol to cytochrome c (GO:0006122) | 5 | 10.43 | 2.91E-02 |
| protein processing involved in protein targeting to mitochondrion (GO:0006627) | 6 | 8.69 | 4.65E-02 |
| protein N-linked glycosylation via asparagine (GO:0018279) | 9 | 7.72 | 1.96E-02 |
| alternative mRNA splicing, via spliceosome (GO:0000380) | 9 | 7.72 | 1.94E-02 |
| cytoplasmic translation (GO:0002181) | 38 | 5.95 | 6.56E-06 |
| negative regulation of translation (GO:0017148) | 19 | 5.49 | 1.02E-02 |
| negative regulation of cell migration (GO:0030336) | 16 | 5.43 | 2.70E-02 |
| negative regulation of cell motility (GO:2000146) | 17 | 5.11 | 3.35E-02 |
| negative regulation of locomotion (GO:0040013) | 17 | 5.11 | 3.32E-02 |
| translational initiation (GO:0006413) | 31 | 5.05 | 1.35E-03 |
| intrinsic apoptotic signaling pathway in response to DNA damage (GO:0008630) | 18 | 4.83 | 4.17E-02 |
| protein targeting to ER (GO:0045047) | 18 | 4.83 | 4.14E-02 |
| mRNA 3'-end processing (GO:0031124) | 18 | 4.83 | 4.12E-02 |
| establishment of protein localization to endoplasmic reticulum (GO:0072599) | 18 | 4.83 | 4.09E-02 |
| MAPK cascade (GO:0000165) | 22 | 4.74 | 2.11E-02 |
| positive regulation of mRNA metabolic process (GO:1903313) | 43 | 4.45 | 7.66E-04 |
| regulation of translation (GO:0006417) | 61 | 4.27 | 6.81E-05 |
| regulation of amide metabolic process (GO:0034248) | 64 | 4.07 | 1.22E-04 |
| regulation of mRNA splicing, via spliceosome (GO:0048024) | 39 | 4.01 | 6.96E-03 |
| positive regulation of Wnt signaling pathway (GO:0030177) | 26 | 4.01 | 4.31E-02 |
| mRNA processing (GO:0006397) | 166 | 3.66 | 2.86E-09 |
| regulation of RNA splicing (GO:0043484) | 72 | 3.62 | 4.76E-04 |
| mRNA splicing, via spliceosome (GO:0000398) | 140 | 3.6 | 1.52E-07 |
| RNA splicing, via transesterification reactions with bulged adenosine as nucleophile (GO:0000377) | 140 | 3.6 | 1.45E-07 |
| RNA splicing, via transesterification reactions (GO:0000375) | 140 | 3.6 | 1.38E-07 |
| ribosomal large subunit biogenesis (GO:0042273) | 49 | 3.55 | 8.51E-03 |
| cell-matrix adhesion (GO:0007160) | 41 | 3.39 | 3.24E-02 |
| mRNA metabolic process (GO:0016071) | 227 | 3.37 | 2.08E-10 |
| post-transcriptional regulation of gene expression (GO:0010608) | 125 | 3.34 | 1.01E-05 |
| regulation of mRNA processing (GO:0050684) | 47 | 3.33 | 2.17E-02 |
| RNA splicing (GO:0008380) | 157 | 3.32 | 4.92E-07 |
| heterochromatin formation (GO:0031507) | 42 | 3.31 | 3.59E-02 |
| peptide biosynthetic process (GO:0043043) | 163 | 3.31 | 3.23E-07 |
| translation (GO:0006412) | 160 | 3.26 | 7.47E-07 |
| protein polyubiquitination (GO:0000209) | 101 | 3.1 | 5.81E-04 |
| regulation of mRNA metabolic process (GO:1903311) | 107 | 3.09 | 4.24E-04 |
| regulation of mRNA stability (GO:0043488) | 58 | 3 | 2.71E-02 |
| peptidyl-serine phosphorylation (GO:0018105) | 93 | 2.99 | 2.26E-03 |
| peptidyl-serine modification (GO:0018209) | 94 | 2.96 | 2.47E-03 |
| proteasome-mediated ubiquitin-dependent protein catabolic process (GO:0043161) | 141 | 2.96 | 7.90E-05 |
| amide biosynthetic process (GO:0043604) | 200 | 2.95 | 1.01E-06 |
| regulation of RNA stability (GO:0043487) | 59 | 2.95 | 3.00E-02 |
| mRNA catabolic process (GO:0006402) | 54 | 2.9 | 4.74E-02 |
| regulation of mRNA catabolic process (GO:0061013) | 62 | 2.8 | 3.96E-02 |
| peptide metabolic process (GO:0006518) | 227 | 2.76 | 2.21E-06 |
| transcription by RNA polymerase II (GO:0006366) | 70 | 2.73 | 3.22E-02 |
| proteasomal protein catabolic process (GO:0010498) | 190 | 2.65 | 7.19E-05 |
| protein folding (GO:0006457) | 106 | 2.62 | 7.82E-03 |
| ubiquitin-dependent protein catabolic process (GO:0006511) | 263 | 2.58 | 3.62E-06 |
| positive regulation of catabolic process (GO:0009896) | 81 | 2.57 | 3.38E-02 |
| protein phosphorylation (GO:0006468) | 243 | 2.57 | 1.00E-05 |
| modification-dependent macromolecule catabolic process (GO:0043632) | 277 | 2.57 | 1.94E-06 |
| modification-dependent protein catabolic process (GO:0019941) | 271 | 2.57 | 2.88E-06 |
| protein-RNA complex assembly (GO:0022618) | 91 | 2.48 | 3.25E-02 |
| negative regulation of gene expression (GO:0010629) | 176 | 2.47 | 8.73E-04 |
| intracellular protein transport (GO:0006886) | 226 | 2.46 | 1.67E-04 |
| ribosome biogenesis (GO:0042254) | 156 | 2.45 | 2.51E-03 |
| ribonucleoprotein complex biogenesis (GO:0022613) | 215 | 2.43 | 3.77E-04 |
| chromatin organization (GO:0006325) | 151 | 2.42 | 3.97E-03 |
| protein targeting (GO:0006605) | 101 | 2.41 | 3.07E-02 |
| phosphorylation (GO:0016310) | 260 | 2.41 | 5.79E-05 |
| protein-RNA complex organization (GO:0071826) | 95 | 2.38 | 4.34E-02 |
| establishment of protein localization to organelle (GO:0072594) | 157 | 2.32 | 7.90E-03 |
| proteolysis involved in protein catabolic process (GO:0051603) | 338 | 2.31 | 1.01E-05 |
| chromatin remodeling (GO:0006338) | 138 | 2.27 | 2.14E-02 |
| protein catabolic process (GO:0030163) | 347 | 2.25 | 1.73E-05 |
| amide metabolic process (GO:0043603) | 324 | 2.25 | 4.27E-05 |
| protein-DNA complex organization (GO:0071824) | 174 | 2.2 | 9.58E-03 |
| gene expression (GO:0010467) | 919 | 2.19 | 5.81E-13 |
| regulation of catabolic process (GO:0009894) | 159 | 2.19 | 1.64E-02 |
| macromolecule catabolic process (GO:0009057) | 436 | 2.15 | 6.63E-06 |
| RNA processing (GO:0006396) | 415 | 2.14 | 1.70E-05 |
| peptidyl-amino acid modification (GO:0018193) | 257 | 2.03 | 5.24E-03 |
| establishment of protein localization (GO:0045184) | 382 | 2 | 4.46E-04 |
| protein maturation (GO:0051604) | 201 | 1.99 | 3.04E-02 |
| RNA metabolic process (GO:0016070) | 639 | 1.99 | 1.53E-06 |
| cellular nitrogen compound biosynthetic process (GO:0044271) | 456 | 1.98 | 1.46E-04 |
| protein transport (GO:0015031) | 308 | 1.98 | 2.98E-03 |
| macromolecule biosynthetic process (GO:0009059) | 1144 | 1.94 | 6.11E-11 |
| positive regulation of RNA metabolic process (GO:0051254) | 316 | 1.87 | 1.04E-02 |
| proteolysis (GO:0006508) | 530 | 1.87 | 2.79E-04 |
| organonitrogen compound catabolic process (GO:1901565) | 497 | 1.85 | 5.36E-04 |
| protein modification by small protein conjugation (GO:0032446) | 302 | 1.84 | 1.83E-02 |
| cellular macromolecule localization (GO:0070727) | 571 | 1.83 | 2.75E-04 |
| protein localization (GO:0008104) | 571 | 1.83 | 2.69E-04 |
| protein metabolic process (GO:0019538) | 1715 | 1.79 | 2.75E-12 |
| positive regulation of nucleobase-containing compound metabolic process (GO:0045935) | 341 | 1.78 | 1.65E-02 |
| cellular response to stress (GO:0033554) | 416 | 1.75 | 8.32E-03 |
| positive regulation of nitrogen compound metabolic process (GO:0051173) | 526 | 1.75 | 2.42E-03 |
| protein-containing complex organization (GO:0043933) | 462 | 1.73 | 7.94E-03 |
| regulation of protein metabolic process (GO:0051246) | 372 | 1.73 | 2.48E-02 |
| cellular biosynthetic process (GO:0044249) | 1513 | 1.72 | 3.65E-09 |
| cellular nitrogen compound metabolic process (GO:0034641) | 1327 | 1.72 | 7.90E-08 |
| macromolecule metabolic process (GO:0043170) | 2631 | 1.7 | 7.11E-16 |
| organic substance biosynthetic process (GO:1901576) | 1553 | 1.69 | 1.29E-08 |
| biosynthetic process (GO:0009058) | 1557 | 1.69 | 1.32E-08 |
| catabolic process (GO:0009056) | 809 | 1.65 | 5.17E-04 |
| organonitrogen compound metabolic process (GO:1901564) | 2190 | 1.65 | 3.42E-11 |
| nitrogen compound transport (GO:0071705) | 495 | 1.65 | 1.48E-02 |
| organonitrogen compound biosynthetic process (GO:1901566) | 591 | 1.65 | 6.99E-03 |
| positive regulation of macromolecule metabolic process (GO:0010604) | 574 | 1.64 | 7.84E-03 |
| organic substance catabolic process (GO:1901575) | 694 | 1.63 | 3.14E-03 |
| nitrogen compound metabolic process (GO:0006807) | 3052 | 1.62 | 3.12E-15 |
| positive regulation of metabolic process (GO:0009893) | 643 | 1.62 | 6.36E-03 |
| intracellular transport (GO:0046907) | 550 | 1.61 | 1.42E-02 |
| nucleic acid metabolic process (GO:0090304) | 909 | 1.61 | 5.58E-04 |
| macromolecule localization (GO:0033036) | 748 | 1.6 | 2.95E-03 |
| protein modification process (GO:0036211) | 1008 | 1.57 | 6.68E-04 |
| positive regulation of cellular metabolic process (GO:0031325) | 570 | 1.56 | 2.95E-02 |
| negative regulation of cellular metabolic process (GO:0031324) | 528 | 1.55 | 4.36E-02 |
| nucleobase-containing compound metabolic process (GO:0006139) | 1128 | 1.54 | 5.77E-04 |
| macromolecule modification (GO:0043412) | 1118 | 1.54 | 5.64E-04 |
| cellular metabolic process (GO:0044237) | 2833 | 1.52 | 3.67E-10 |
| primary metabolic process (GO:0044238) | 3363 | 1.5 | 2.81E-11 |
| cellular localization (GO:0051641) | 961 | 1.48 | 8.01E-03 |
| heterocycle metabolic process (GO:0046483) | 1184 | 1.48 | 1.85E-03 |
| phosphate-containing compound metabolic process (GO:0006796) | 727 | 1.48 | 3.09E-02 |
| phosphorus metabolic process (GO:0006793) | 731 | 1.47 | 3.23E-02 |
| cellular aromatic compound metabolic process (GO:0006725) | 1186 | 1.47 | 2.89E-03 |
| metabolic process (GO:0008152) | 3777 | 1.45 | 4.17E-11 |
| negative regulation of cellular process (GO:0048523) | 984 | 1.45 | 1.33E-02 |
| organic substance metabolic process (GO:0071704) | 3571 | 1.45 | 5.16E-10 |
| organic cyclic compound metabolic process (GO:1901360) | 1256 | 1.4 | 1.06E-02 |
| negative regulation of biological process (GO:0048519) | 1069 | 1.38 | 3.45E-02 |
| regulation of macromolecule metabolic process (GO:0060255) | 2686 | 1.29 | 2.35E-03 |
| regulation of metabolic process (GO:0019222) | 2828 | 1.29 | 1.95E-03 |
| regulation of gene expression (GO:0010468) | 2336 | 1.29 | 7.81E-03 |
| regulation of nitrogen compound metabolic process (GO:0051171) | 2435 | 1.28 | 7.38E-03 |
| regulation of cellular metabolic process (GO:0031323) | 2668 | 1.28 | 4.00E-03 |
| regulation of macromolecule biosynthetic process (GO:0010556) | 2363 | 1.28 | 9.50E-03 |
| regulation of primary metabolic process (GO:0080090) | 2461 | 1.28 | 8.45E-03 |
| regulation of biosynthetic process (GO:0009889) | 2386 | 1.27 | 1.11E-02 |
| regulation of cellular biosynthetic process (GO:0031326) | 2380 | 1.27 | 1.30E-02 |
| cellular component organization or biogenesis (GO:0071840) | 2251 | 1.27 | 2.12E-02 |
| cellular process (GO:0009987) | 7925 | 1.19 | 3.91E-06 |
| biological_process (GO:0008150) | 11637 | 1.15 | 1.60E-08 |
| Unclassified (UNCLASSIFIED) | 6925 | 0.76 | 1.50E-08 |
| G protein-coupled receptor signaling pathway (GO:0007186) | 449 | 0.35 | 4.59E-03 |
| immune response (GO:0006955) | 273 | 0.32 | 3.57E-02 |
| defense response to other organism (GO:0098542) | 215 | 0.24 | 4.08E-02 |
| system process (GO:0003008) | 364 | 0.24 | 1.12E-03 |
| response to external biotic stimulus (GO:0043207) | 248 | 0.21 | 1.02E-02 |
| response to other organism (GO:0051707) | 248 | 0.21 | 1.01E-02 |
| biological process involved in interspecies interaction between organisms (GO:0044419) | 250 | 0.21 | 1.01E-02 |
| response to biotic stimulus (GO:0009607) | 254 | 0.21 | 7.96E-03 |
| defense response (GO:0006952) | 283 | 0.18 | 2.49E-03 |
| nervous system process (GO:0050877) | 270 | 0.13 | 9.94E-04 |
| sensory perception of chemical stimulus (GO:0007606) | 199 | 0.09 | 4.35E-03 |
| sensory perception (GO:0007600) | 243 | 0.07 | 6.46E-04 |
| detection of chemical stimulus involved in sensory perception (GO:0050907) | 155 | < 0.01 | 4.45E-03 |
| detection of stimulus involved in sensory perception (GO:0050906) | 161 | < 0.01 | 3.15E-03 |
| detection of stimulus (GO:0051606) | 193 | < 0.01 | 5.86E-04 |
| detection of chemical stimulus (GO:0009593) | 161 | < 0.01 | 3.11E-03 |
| **PANTHER Protein Class** | **REFLIST- Total WTA Gene Set** | **YS-HepIII /noYS Fold Enrichment** | **Adjusted P-Value** |
| chaperonin (PC00073) | 12 | 5.79 | 3.17E-02 |
| translation elongation factor (PC00222) | 14 | 4.97 | 4.63E-02 |
| Hsp70 family chaperone (PC00027) | 14 | 4.97 | 4.48E-02 |
| ribosomal protein (PC00202) | 171 | 3.96 | 5.08E-12 |
| integrin (PC00126) | 28 | 3.72 | 3.54E-02 |
| RNA helicase (PC00032) | 71 | 3.67 | 1.77E-04 |
| RNA splicing factor (PC00148) | 129 | 3.5 | 5.73E-07 |
| translational protein (PC00263) | 322 | 3.4 | 5.83E-16 |
| translation factor (PC00223) | 108 | 3.38 | 1.49E-05 |
| translation initiation factor (PC00224) | 78 | 3.34 | 4.90E-04 |
| C4 zinc finger nuclear receptor (PC00169) | 49 | 2.84 | 4.67E-02 |
| RNA processing factor (PC00147) | 245 | 2.77 | 2.74E-07 |
| RNA metabolism protein (PC00031) | 778 | 2.59 | 2.15E-19 |
| small GTPase (PC00208) | 121 | 2.59 | 2.32E-03 |
| kinase modulator (PC00140) | 93 | 2.43 | 2.35E-02 |
| G-protein (PC00020) | 185 | 2.35 | 1.09E-03 |
| general transcription factor (PC00259) | 99 | 2.28 | 4.53E-02 |
| chaperone (PC00072) | 198 | 2.19 | 3.09E-03 |
| non-receptor serine/threonine protein kinase (PC00167) | 340 | 1.94 | 1.11E-03 |
| chromatin/chromatin-binding, or -regulatory protein (PC00077) | 291 | 1.73 | 3.56E-02 |
| ubiquitin-protein ligase (PC00234) | 394 | 1.72 | 9.16E-03 |
| protein modifying enzyme (PC00260) | 1564 | 1.43 | 4.11E-04 |
| protein class (PC00000) | 13037 | 1.1 | 4.24E-06 |
| Unclassified (UNCLASSIFIED) | 5525 | 0.76 | 3.71E-06 |
| hydrolase (PC00121) | 462 | 0.49 | 3.55E-02 |
| C2H2 zinc finger transcription factor (PC00248) | 618 | 0.45 | 2.39E-03 |
| intercellular signal molecule (PC00207) | 386 | 0.36 | 8.28E-03 |
| transmembrane signal receptor (PC00197) | 1075 | 0.26 | 4.86E-11 |
| structural protein (PC00211) | 181 | 0.19 | 2.92E-02 |
| G-protein coupled receptor (PC00021) | 386 | 0.14 | 8.71E-06 |
| **REACTOME Pathways** | **REFLIST- Total WTA Gene Set** | **YS-HepIII /noYS Fold Enrichment** | **Adjusted P-Value** |
| 2-LTR circle formation (R-HSA-164843) | 7 | 9.93 | 2.63E-03 |
| ABC transporter disorders (R-HSA-5619084) | 76 | 3.2 | 8.95E-04 |
| ABC-family proteins mediated transport (R-HSA-382556) | 101 | 2.41 | 1.15E-02 |
| activated TAK1 mediates p38 MAPK activation (R-HSA-450302) | 22 | 3.95 | 3.76E-02 |
| Activation of anterior HOX genes in hindbrain development during early embryogenesis (R-HSA-5617472) | 88 | 4.74 | 3.11E-09 |
| Activation of APC/C and APC/C:Cdc20 mediated degradation of mitotic proteins (R-HSA-176814) | 75 | 3.71 | 5.17E-05 |
| Activation of BAD and translocation to mitochondria (R-HSA-111447) | 15 | 5.79 | 7.36E-03 |
| Activation of BH3-only proteins (R-HSA-114452) | 29 | 4.79 | 1.46E-03 |
| Activation of BIM and translocation to mitochondria (R-HSA-111446) | 3 | 11.59 | 4.83E-02 |
| Activation of HOX genes during differentiation (R-HSA-5619507) | 88 | 4.74 | 3.15E-09 |
| Activation of NF-kappaB in B cells (R-HSA-1169091) | 65 | 4.55 | 1.86E-06 |
| Activation of the mRNA upon binding of the cap-binding complex and eIFs, and subsequent binding to 43S (R-HSA-72662) | 60 | 6.66 | 3.63E-12 |
| Advanced glycosylation endproduct receptor signaling (R-HSA-879415) | 12 | 7.24 | 2.77E-03 |
| Aggrephagy (R-HSA-9646399) | 40 | 3.91 | 2.83E-03 |
| Amyloid fiber formation (R-HSA-977225) | 75 | 3.01 | 2.53E-03 |
| Antigen processing: Ubiquitination & Proteasome degradation (R-HSA-983168) | 304 | 2.34 | 5.59E-06 |
| Antigen processing-Cross presentation (R-HSA-1236975) | 103 | 2.87 | 7.29E-04 |
| Antiviral mechanism by IFN-stimulated genes (R-HSA-1169410) | 142 | 2.82 | 7.09E-05 |
| APC/C:Cdc20 mediated degradation of mitotic proteins (R-HSA-176409) | 74 | 3.76 | 4.41E-05 |
| APC/C:Cdh1 mediated degradation of Cdc20 and other APC/C:Cdh1 targeted proteins in late mitosis/early G1 (R-HSA-174178) | 72 | 3.62 | 1.32E-04 |
| APC/C-mediated degradation of cell cycle proteins (R-HSA-174143) | 86 | 3.64 | 2.00E-05 |
| APC:Cdc20 mediated degradation of cell cycle proteins prior to satisfation of the cell cycle checkpoint (R-HSA-179419) | 72 | 3.86 | 3.14E-05 |
| APOBEC3G mediated resistance to HIV-1 infection (R-HSA-180689) | 4 | 13.04 | 5.21E-03 |
| Apoptosis (R-HSA-109581) | 172 | 3.64 | 3.83E-10 |
| Apoptosis induced DNA fragmentation (R-HSA-140342) | 13 | 6.68 | 4.08E-03 |
| Assembly and release of respiratory syncytial virus (RSV) virions (R-HSA-9820962) | 2 | 17.38 | 1.92E-02 |
| Assembly of the pre-replicative complex (R-HSA-68867) | 106 | 3.94 | 1.47E-07 |
| Asymmetric localization of PCP proteins (R-HSA-4608870) | 62 | 3.64 | 4.31E-04 |
| Attenuation phase (R-HSA-3371568) | 14 | 7.45 | 6.92E-04 |
| AUF1 (hnRNP D0) binds and destabilizes mRNA (R-HSA-450408) | 52 | 5.01 | 2.46E-06 |
| Autodegradation of Cdh1 by Cdh1:APC/C (R-HSA-174084) | 62 | 4.2 | 2.14E-05 |
| Autodegradation of the E3 ubiquitin ligase COP1 (R-HSA-349425) | 50 | 4.87 | 8.03E-06 |
| Autointegration results in viral DNA circles (R-HSA-177539) | 3 | 17.38 | 1.58E-03 |
| Autophagy (R-HSA-9612973) | 143 | 2.92 | 2.61E-05 |
| Axon guidance (R-HSA-422475) | 540 | 3.25 | 2.61E-24 |
| Bacterial Infection Pathways (R-HSA-9824439) | 64 | 4.07 | 3.15E-05 |
| Beta-catenin independent WNT signaling (R-HSA-3858494) | 141 | 2.96 | 2.09E-05 |
| B-WICH complex positively regulates rRNA expression (R-HSA-5250924) | 52 | 3.68 | 1.30E-03 |
| Cap-dependent Translation Initiation (R-HSA-72737) | 119 | 7.01 | 2.92E-26 |
| Cdc20:Phospho-APC/C mediated degradation of Cyclin A (R-HSA-174184) | 71 | 3.92 | 2.60E-05 |
| CDK-mediated phosphorylation and removal of Cdc6 (R-HSA-69017) | 71 | 3.92 | 2.61E-05 |
| Cell Cycle (R-HSA-1640170) | 636 | 2.51 | 1.16E-14 |
| Cell Cycle Checkpoints (R-HSA-69620) | 265 | 2.62 | 4.84E-07 |
| Cell Cycle, Mitotic (R-HSA-69278) | 506 | 2.61 | 6.12E-13 |
| Cellular response to chemical stress (R-HSA-9711123) | 203 | 3.34 | 8.23E-10 |
| Cellular response to hypoxia (R-HSA-1234174) | 73 | 4.29 | 2.07E-06 |
| Cellular response to starvation (R-HSA-9711097) | 156 | 4.79 | 1.50E-16 |
| Cellular responses to stimuli (R-HSA-8953897) | 763 | 3.28 | 1.08E-35 |
| Cellular responses to stress (R-HSA-2262752) | 749 | 3.34 | 1.41E-36 |
| Cellular Senescence (R-HSA-2559583) | 159 | 3.17 | 6.39E-07 |
| Chaperonin-mediated protein folding (R-HSA-390466) | 85 | 2.66 | 7.03E-03 |
| Chromatin modifications during the maternal to zygotic transition (MZT) (R-HSA-9821002) | 36 | 3.38 | 2.19E-02 |
| Chromatin modifying enzymes (R-HSA-3247509) | 234 | 2.6 | 3.44E-06 |
| Chromatin organization (R-HSA-4839726) | 234 | 2.6 | 3.41E-06 |
| Circadian Clock (R-HSA-400253) | 64 | 2.99 | 6.43E-03 |
| Class I MHC mediated antigen processing & presentation (R-HSA-983169) | 376 | 2.13 | 2.00E-05 |
| Clathrin-mediated endocytosis (R-HSA-8856828) | 141 | 2.47 | 1.44E-03 |
| CLEC7A (Dectin-1) signaling (R-HSA-5607764) | 95 | 3.84 | 1.51E-06 |
| Constitutive Signaling by EGFRvIII (R-HSA-5637810) | 13 | 6.68 | 4.11E-03 |
| Constitutive Signaling by Ligand-Responsive EGFR Cancer Variants (R-HSA-1236382) | 16 | 6.52 | 1.48E-03 |
| Constitutive Signaling by Overexpressed ERBB2 (R-HSA-9634285) | 9 | 7.72 | 7.00E-03 |
| Cooperation of PDCL (PhLP1) and TRiC/CCT in G-protein beta folding (R-HSA-6814122) | 37 | 3.29 | 2.54E-02 |
| Cooperation of Prefoldin and TriC/CCT in actin and tubulin folding (R-HSA-389958) | 30 | 4.63 | 1.80E-03 |
| Cross-presentation of soluble exogenous antigens (endosomes) (R-HSA-1236978) | 48 | 4.35 | 1.38E-04 |
| C-type lectin receptors (CLRs) (R-HSA-5621481) | 136 | 3.32 | 1.12E-06 |
| Cyclin A:Cdk2-associated events at S phase entry (R-HSA-69656) | 83 | 4.4 | 1.57E-07 |
| Cyclin D associated events in G1 (R-HSA-69231) | 45 | 3.09 | 2.12E-02 |
| Cyclin E associated events during G1/S transition (R-HSA-69202) | 81 | 4.51 | 1.01E-07 |
| Cytokine Signaling in Immune system (R-HSA-1280215) | 747 | 2.02 | 9.82E-09 |
| DCC mediated attractive signaling (R-HSA-418885) | 13 | 6.68 | 4.07E-03 |
| Deadenylation of mRNA (R-HSA-429947) | 25 | 9.73 | 3.68E-10 |
| Deadenylation-dependent mRNA decay (R-HSA-429914) | 55 | 5.06 | 9.79E-07 |
| Dectin-1 mediated noncanonical NF-kB signaling (R-HSA-5607761) | 58 | 4.49 | 9.49E-06 |
| Defective CFTR causes cystic fibrosis (R-HSA-5678895) | 59 | 4.12 | 5.37E-05 |
| Defective Intrinsic Pathway for Apoptosis (R-HSA-9734009) | 23 | 6.8 | 3.25E-05 |
| Defective pyroptosis (R-HSA-9710421) | 38 | 3.66 | 7.70E-03 |
| Degradation of AXIN (R-HSA-4641257) | 53 | 4.59 | 1.61E-05 |
| Degradation of beta-catenin by the destruction complex (R-HSA-195253) | 81 | 4.51 | 1.02E-07 |
| Degradation of DVL (R-HSA-4641258) | 55 | 4.74 | 5.09E-06 |
| Degradation of GLI1 by the proteasome (R-HSA-5610780) | 58 | 5.09 | 3.58E-07 |
| Degradation of GLI2 by the proteasome (R-HSA-5610783) | 58 | 5.09 | 3.61E-07 |
| Deubiquitination (R-HSA-5688426) | 264 | 2.24 | 1.30E-04 |
| Developmental Biology (R-HSA-1266738) | 1204 | 2.6 | 2.29E-31 |
| Disease (R-HSA-1643685) | 1734 | 2.48 | 1.19E-40 |
| Diseases of programmed cell death (R-HSA-9645723) | 68 | 4.35 | 3.43E-06 |
| Diseases of signal transduction by growth factor receptors and second messengers (R-HSA-5663202) | 441 | 2.8 | 1.40E-13 |
| DNA Double Strand Break Response (R-HSA-5693606) | 57 | 3.96 | 1.73E-04 |
| DNA Double-Strand Break Repair (R-HSA-5693532) | 143 | 2.19 | 1.02E-02 |
| DNA methylation (R-HSA-5334118) | 30 | 3.48 | 3.40E-02 |
| DNA Replication (R-HSA-69306) | 151 | 3.22 | 7.56E-07 |
| DNA Replication Pre-Initiation (R-HSA-69002) | 123 | 3.39 | 2.17E-06 |
| Downregulation of SMAD2/3:SMAD4 transcriptional activity (R-HSA-2173795) | 28 | 4.35 | 5.70E-03 |
| Downstream signaling events of B Cell Receptor (BCR) (R-HSA-1168372) | 78 | 4.01 | 5.36E-06 |
| Downstream TCR signaling (R-HSA-202424) | 93 | 4.3 | 5.22E-08 |
| Drug resistance in ERBB2 KD mutants (R-HSA-9665230) | 4 | 13.04 | 5.07E-03 |
| Drug resistance in ERBB2 TMD/JMD mutants (R-HSA-9665737) | 4 | 13.04 | 5.03E-03 |
| Drug-mediated inhibition of ERBB2 signaling (R-HSA-9652282) | 4 | 13.04 | 5.19E-03 |
| E3 ubiquitin ligases ubiquitinate target proteins (R-HSA-8866654) | 52 | 3.68 | 1.30E-03 |
| Early Phase of HIV Life Cycle (R-HSA-162594) | 14 | 4.97 | 3.58E-02 |
| Entry of Influenza Virion into Host Cell via Endocytosis (R-HSA-168275) | 2 | 17.38 | 1.91E-02 |
| EPHB-mediated forward signaling (R-HSA-3928662) | 38 | 3.66 | 7.68E-03 |
| EPH-ephrin mediated repulsion of cells (R-HSA-3928665) | 48 | 2.9 | 3.04E-02 |
| EPH-Ephrin signaling (R-HSA-2682334) | 88 | 3.75 | 7.43E-06 |
| Epigenetic regulation of gene expression (R-HSA-212165) | 148 | 3.05 | 5.19E-06 |
| ER Quality Control Compartment (ERQC) (R-HSA-901032) | 20 | 4.35 | 2.57E-02 |
| ERCC6 (CSB) and EHMT2 (G9a) positively regulate rRNA expression (R-HSA-427389) | 40 | 4.35 | 6.46E-04 |
| ER-Phagosome pathway (R-HSA-1236974) | 88 | 3.16 | 3.86E-04 |
| ESR-mediated signaling (R-HSA-8939211) | 183 | 3.89 | 1.81E-12 |
| Estrogen-dependent gene expression (R-HSA-9018519) | 114 | 4.88 | 1.53E-12 |
| Eukaryotic Translation Elongation (R-HSA-156842) | 91 | 7.26 | 1.66E-21 |
| Eukaryotic Translation Initiation (R-HSA-72613) | 119 | 7.01 | 2.66E-26 |
| Eukaryotic Translation Termination (R-HSA-72764) | 93 | 6.91 | 3.80E-20 |
| Extra-nuclear estrogen signaling (R-HSA-9009391) | 73 | 2.86 | 5.94E-03 |
| FBXL7 down-regulates AURKA during mitotic entry and in early mitosis (R-HSA-8854050) | 53 | 5.25 | 5.83E-07 |
| FBXW7 Mutants and NOTCH1 in Cancer (R-HSA-2644605) | 4 | 13.04 | 5.04E-03 |
| Fc epsilon receptor (FCERI) signaling (R-HSA-2454202) | 126 | 3.45 | 9.85E-07 |
| FCERI mediated NF-kB activation (R-HSA-2871837) | 80 | 4.13 | 1.83E-06 |
| Fcgamma receptor (FCGR) dependent phagocytosis (R-HSA-2029480) | 85 | 4.09 | 1.09E-06 |
| FCGR3A-mediated phagocytosis (R-HSA-9664422) | 59 | 3.53 | 1.02E-03 |
| FGFR2 alternative splicing (R-HSA-6803529) | 25 | 4.17 | 1.46E-02 |
| Folding of actin by CCT/TriC (R-HSA-390450) | 9 | 7.72 | 7.03E-03 |
| Formation of a pool of free 40S subunits (R-HSA-72689) | 101 | 7.23 | 1.21E-23 |
| Formation of Incision Complex in GG-NER (R-HSA-5696395) | 41 | 3.82 | 3.40E-03 |
| Formation of paraxial mesoderm (R-HSA-9793380) | 68 | 4.09 | 1.55E-05 |
| Formation of TC-NER Pre-Incision Complex (R-HSA-6781823) | 50 | 3.48 | 3.64E-03 |
| Formation of the beta-catenin:TCF transactivating complex (R-HSA-201722) | 56 | 4.03 | 1.43E-04 |
| Formation of the ternary complex, and subsequently, the 43S complex (R-HSA-72695) | 52 | 6.02 | 8.73E-09 |
| Formation of tubulin folding intermediates by CCT/TriC (R-HSA-389960) | 24 | 4.35 | 1.18E-02 |
| Formation of WDR5-containing histone-modifying complexes (R-HSA-9772755) | 42 | 2.9 | 4.78E-02 |
| FOXO-mediated transcription (R-HSA-9614085) | 65 | 2.94 | 7.10E-03 |
| G1 Phase (R-HSA-69236) | 45 | 3.09 | 2.12E-02 |
| G1/S Transition (R-HSA-69206) | 129 | 3.37 | 1.49E-06 |
| G2/M Transition (R-HSA-69275) | 190 | 2.93 | 8.80E-07 |
| Gain-of-function MRAS complexes activate RAF signaling (R-HSA-9726842) | 8 | 8.69 | 4.63E-03 |
| Gastrulation (R-HSA-9758941) | 123 | 2.68 | 7.22E-04 |
| Gene and protein expression by JAK-STAT signaling after Interleukin-12 stimulation (R-HSA-8950505) | 37 | 3.76 | 6.67E-03 |
| Gene expression (Transcription) (R-HSA-74160) | 1466 | 2.03 | 9.12E-18 |
| Gene Silencing by RNA (R-HSA-211000) | 100 | 2.61 | 4.28E-03 |
| GLI3 is processed to GLI3R by the proteasome (R-HSA-5610785) | 58 | 5.09 | 3.64E-07 |
| Global Genome Nucleotide Excision Repair (GG-NER) (R-HSA-5696399) | 81 | 2.57 | 1.32E-02 |
| Golgi Cisternae Pericentriolar Stack Reorganization (R-HSA-162658) | 13 | 5.35 | 2.78E-02 |
| GSK3B and BTRC:CUL1-mediated-degradation of NFE2L2 (R-HSA-9762114) | 50 | 5.56 | 2.47E-07 |
| GTP hydrolysis and joining of the 60S ribosomal subunit (R-HSA-72706) | 112 | 7.29 | 1.44E-26 |
| HATs acetylate histones (R-HSA-3214847) | 104 | 2.67 | 2.31E-03 |
| HCMV Early Events (R-HSA-9609690) | 94 | 2.96 | 7.96E-04 |
| HCMV Infection (R-HSA-9609646) | 118 | 2.65 | 1.21E-03 |
| HDACs deacetylate histones (R-HSA-3214815) | 57 | 3.05 | 8.67E-03 |
| HDR through Homologous Recombination (HRR) or Single Strand Annealing (SSA) (R-HSA-5693567) | 107 | 2.27 | 2.72E-02 |
| Hedgehog ligand biogenesis (R-HSA-5358346) | 63 | 4.14 | 2.62E-05 |
| Hedgehog 'off' state (R-HSA-5610787) | 109 | 3.99 | 5.63E-08 |
| Hedgehog 'on' state (R-HSA-5632684) | 83 | 3.98 | 3.17E-06 |
| Hh mutants abrogate ligand secretion (R-HSA-5387390) | 57 | 4.57 | 7.86E-06 |
| Hh mutants are degraded by ERAD (R-HSA-5362768) | 54 | 4.83 | 3.98E-06 |
| Homology Directed Repair (R-HSA-5693538) | 113 | 2.15 | 3.46E-02 |
| Host Interactions of HIV factors (R-HSA-162909) | 123 | 4.38 | 7.00E-11 |
| HSF1 activation (R-HSA-3371511) | 11 | 6.32 | 1.53E-02 |
| HSF1-dependent transactivation (R-HSA-3371571) | 24 | 5.07 | 2.38E-03 |
| HSP90 chaperone cycle for steroid hormone receptors (SHR) in the presence of ligand (R-HSA-3371497) | 54 | 4.51 | 1.99E-05 |
| Infection with Mycobacterium tuberculosis (R-HSA-9635486) | 26 | 5.35 | 6.87E-04 |
| Infectious disease (R-HSA-5663205) | 955 | 3.26 | 4.11E-44 |
| Inflammasomes (R-HSA-622312) | 21 | 4.14 | 3.13E-02 |
| Influenza Infection (R-HSA-168255) | 152 | 5.49 | 3.94E-21 |
| Influenza Viral RNA Transcription and Replication (R-HSA-168273) | 132 | 5.53 | 1.15E-18 |
| Inhibition of DNA recombination at telomere (R-HSA-9670095) | 44 | 3.16 | 1.89E-02 |
| Integration of provirus (R-HSA-162592) | 9 | 7.72 | 7.02E-03 |
| Integration of viral DNA into host genomic DNA (R-HSA-175567) | 3 | 17.38 | 1.59E-03 |
| Interferon Signaling (R-HSA-913531) | 253 | 2.06 | 1.51E-03 |
| Interleukin-1 family signaling (R-HSA-446652) | 148 | 2.58 | 4.05E-04 |
| Interleukin-1 signaling (R-HSA-9020702) | 110 | 3.48 | 4.17E-06 |
| Interleukin-12 signaling (R-HSA-9020591) | 46 | 3.02 | 2.40E-02 |
| Interleukin-17 signaling (R-HSA-448424) | 70 | 2.73 | 1.24E-02 |
| Intracellular signaling by second messengers (R-HSA-9006925) | 290 | 2.7 | 3.35E-08 |
| Intrinsic Pathway for Apoptosis (R-HSA-109606) | 52 | 4.01 | 3.13E-04 |
| ISG15 antiviral mechanism (R-HSA-1169408) | 69 | 3.02 | 3.95E-03 |
| Josephin domain DUBs (R-HSA-5689877) | 11 | 7.9 | 1.78E-03 |
| KEAP1-NFE2L2 pathway (R-HSA-9755511) | 120 | 4.35 | 1.94E-10 |
| L13a-mediated translational silencing of Ceruloplasmin expression (R-HSA-156827) | 111 | 7.2 | 8.19E-26 |
| L1CAM interactions (R-HSA-373760) | 113 | 2.77 | 7.30E-04 |
| Laminin interactions (R-HSA-3000157) | 30 | 3.48 | 3.42E-02 |
| Late Phase of HIV Life Cycle (R-HSA-162599) | 130 | 2.14 | 2.09E-02 |
| Late SARS-CoV-2 Infection Events (R-HSA-9772573) | 69 | 2.77 | 1.12E-02 |
| LDL clearance (R-HSA-8964038) | 19 | 5.49 | 3.87E-03 |
| Loss of Function of FBXW7 in Cancer and NOTCH1 Signaling (R-HSA-2644607) | 4 | 13.04 | 5.06E-03 |
| Lysosome Vesicle Biogenesis (R-HSA-432720) | 35 | 4.47 | 1.08E-03 |
| M Phase (R-HSA-68886) | 365 | 2.76 | 6.72E-11 |
| Macroautophagy (R-HSA-1632852) | 129 | 2.96 | 4.91E-05 |
| Major pathway of rRNA processing in the nucleolus and cytosol (R-HSA-6791226) | 181 | 4.71 | 1.72E-18 |
| MAPK family signaling cascades (R-HSA-5683057) | 304 | 2.52 | 3.63E-07 |
| MAPK1/MAPK3 signaling (R-HSA-5684996) | 265 | 2.43 | 9.49E-06 |
| MAPK6/MAPK4 signaling (R-HSA-5687128) | 87 | 4 | 1.53E-06 |
| Maternal to zygotic transition (MZT) (R-HSA-9816359) | 85 | 5.11 | 2.35E-10 |
| Maturation of hRSV A proteins (R-HSA-9828806) | 12 | 7.24 | 2.78E-03 |
| M-decay: degradation of maternal mRNAs by maternally stored factors (R-HSA-9820841) | 25 | 10.43 | 2.02E-11 |
| Metabolism of amino acids and derivatives (R-HSA-71291) | 366 | 2.85 | 7.85E-12 |
| Metabolism of polyamines (R-HSA-351202) | 57 | 4.57 | 7.82E-06 |
| Metabolism of proteins (R-HSA-392499) | 1975 | 2.05 | 1.01E-25 |
| Metabolism of RNA (R-HSA-8953854) | 706 | 3.54 | 1.73E-39 |
| MHC class II antigen presentation (R-HSA-2132295) | 120 | 2.32 | 1.32E-02 |
| mRNA Splicing (R-HSA-72172) | 212 | 3.94 | 8.64E-15 |
| MTOR signalling (R-HSA-165159) | 41 | 4.24 | 7.82E-04 |
| mTORC1-mediated signalling (R-HSA-166208) | 24 | 5.07 | 2.39E-03 |
| MyD88 cascade initiated on plasma membrane (R-HSA-975871) | 94 | 2.59 | 6.27E-03 |
| Myogenesis (R-HSA-525793) | 28 | 4.35 | 5.67E-03 |
| Neddylation (R-HSA-8951664) | 241 | 2.38 | 4.57E-05 |
| Nef-mediates down modulation of cell surface receptors by recruiting them to clathrin adapters (R-HSA-164938) | 21 | 4.14 | 3.12E-02 |
| Negative epigenetic regulation of rRNA expression (R-HSA-5250941) | 72 | 2.66 | 1.53E-02 |
| Negative regulation of MAPK pathway (R-HSA-5675221) | 41 | 3.39 | 1.23E-02 |
| Negative regulation of NOTCH4 signaling (R-HSA-9604323) | 51 | 5.79 | 4.88E-08 |
| Nephrin family interactions (R-HSA-373753) | 23 | 3.78 | 4.53E-02 |
| Neurodegenerative Diseases (R-HSA-8863678) | 21 | 6.62 | 1.34E-04 |
| Neutrophil degranulation (R-HSA-6798695) | 466 | 2.2 | 3.14E-07 |
| NFE2L2 regulates pentose phosphate pathway genes (R-HSA-9818028) | 8 | 6.52 | 4.42E-02 |
| NFE2L2 regulating anti-oxidant/detoxification enzymes (R-HSA-9818027) | 18 | 4.83 | 1.67E-02 |
| N-glycan trimming in the ER and Calnexin/Calreticulin cycle (R-HSA-532668) | 34 | 4.09 | 4.13E-03 |
| Noncanonical NF-kB signaling (R-HSA-5676590) | 57 | 4.57 | 7.91E-06 |
| Nonhomologous End-Joining (NHEJ) (R-HSA-5693571) | 48 | 3.26 | 9.14E-03 |
| Non-integrin membrane-ECM interactions (R-HSA-3000171) | 59 | 2.65 | 3.32E-02 |
| Nonsense-Mediated Decay (NMD) (R-HSA-927802) | 115 | 6.05 | 2.40E-19 |
| NOTCH1 Intracellular Domain Regulates Transcription (R-HSA-2122947) | 45 | 3.86 | 1.59E-03 |
| Nuclear Envelope (NE) Reassembly (R-HSA-2995410) | 66 | 2.63 | 2.37E-02 |
| Nuclear events mediated by NFE2L2 (R-HSA-9759194) | 92 | 4.91 | 2.42E-10 |
| Nucleotide Excision Repair (R-HSA-5696398) | 106 | 2.79 | 1.02E-03 |
| Nucleotide-binding domain, leucine rich repeat containing receptor (NLR) signaling pathways (R-HSA-168643) | 55 | 2.84 | 2.17E-02 |
| Oncogenic MAPK signaling (R-HSA-6802957) | 80 | 3.04 | 1.47E-03 |
| Orc1 removal from chromatin (R-HSA-68949) | 69 | 4.79 | 1.78E-07 |
| Oxidative Stress Induced Senescence (R-HSA-2559580) | 87 | 3.8 | 6.23E-06 |
| Oxygen-dependent proline hydroxylation of Hypoxia-inducible Factor Alpha (R-HSA-1234176) | 64 | 4.62 | 1.52E-06 |
| p53-Dependent G1 DNA Damage Response (R-HSA-69563) | 63 | 4.41 | 5.84E-06 |
| p53-Independent DNA Damage Response (R-HSA-69610) | 50 | 4.52 | 4.12E-05 |
| PCP/CE pathway (R-HSA-4086400) | 90 | 3.67 | 9.79E-06 |
| Pentose phosphate pathway (R-HSA-71336) | 15 | 4.63 | 4.56E-02 |
| Peptide chain elongation (R-HSA-156902) | 87 | 7.39 | 2.80E-21 |
| PIP3 activates AKT signaling (R-HSA-1257604) | 250 | 2.71 | 3.04E-07 |
| PKMTs methylate histone lysines (R-HSA-3214841) | 46 | 3.02 | 2.40E-02 |
| PKR-mediated signaling (R-HSA-9833482) | 72 | 2.41 | 4.11E-02 |
| Plasma lipoprotein assembly, remodeling, and clearance (R-HSA-174824) | 70 | 2.73 | 1.23E-02 |
| Plasma lipoprotein clearance (R-HSA-8964043) | 35 | 4.47 | 1.08E-03 |
| Platelet activation, signaling and aggregation (R-HSA-76002) | 258 | 2.02 | 1.85E-03 |
| Positive epigenetic regulation of rRNA expression (R-HSA-5250913) | 67 | 3.89 | 5.35E-05 |
| Post NMDA receptor activation events (R-HSA-438064) | 74 | 2.35 | 4.85E-02 |
| Potential therapeutics for SARS (R-HSA-9679191) | 97 | 3.58 | 7.93E-06 |
| PRC2 methylates histones and DNA (R-HSA-212300) | 38 | 3.66 | 7.66E-03 |
| Prefoldin mediated transfer of substrate to CCT/TriC (R-HSA-389957) | 26 | 5.35 | 6.90E-04 |
| Pre-NOTCH Expression and Processing (R-HSA-1912422) | 74 | 3.52 | 1.81E-04 |
| Pre-NOTCH Transcription and Translation (R-HSA-1912408) | 58 | 3.9 | 2.08E-04 |
| Prevention of phagosomal-lysosomal fusion (R-HSA-9636383) | 8 | 6.52 | 4.41E-02 |
| Processing of Capped Intron-Containing Pre-mRNA (R-HSA-72203) | 277 | 3.64 | 3.03E-16 |
| Processing of Capped Intronless Pre-mRNA (R-HSA-75067) | 28 | 4.35 | 5.69E-03 |
| Processing of DNA double-strand break ends (R-HSA-5693607) | 75 | 3.01 | 2.54E-03 |
| Processing of Intronless Pre-mRNAs (R-HSA-77595) | 19 | 4.57 | 2.10E-02 |
| Programmed Cell Death (R-HSA-5357801) | 204 | 3.49 | 6.65E-11 |
| Protein folding (R-HSA-391251) | 91 | 2.48 | 1.24E-02 |
| Protein ubiquitination (R-HSA-8852135) | 72 | 4.1 | 7.67E-06 |
| PTEN Regulation (R-HSA-6807070) | 135 | 3.86 | 3.88E-09 |
| RAB geranylgeranylation (R-HSA-8873719) | 65 | 2.94 | 7.11E-03 |
| RAC2 GTPase cycle (R-HSA-9013404) | 87 | 2.6 | 8.52E-03 |
| RAF activation (R-HSA-5673000) | 33 | 4.21 | 3.45E-03 |
| RAF/MAP kinase cascade (R-HSA-5673001) | 260 | 2.47 | 7.86E-06 |
| Recruitment and ATM-mediated phosphorylation of repair and signaling proteins at DNA double strand breaks (R-HSA-5693565) | 56 | 4.03 | 1.42E-04 |
| Recycling pathway of L1 (R-HSA-437239) | 43 | 3.64 | 4.65E-03 |
| Regulated Necrosis (R-HSA-5218859) | 57 | 2.74 | 2.71E-02 |
| Regulation of actin dynamics for phagocytic cup formation (R-HSA-2029482) | 61 | 4.27 | 1.79E-05 |
| Regulation of activated PAK-2p34 by proteasome mediated degradation (R-HSA-211733) | 48 | 4.71 | 2.63E-05 |
| Regulation of APC/C activators between G1/S and early anaphase (R-HSA-176408) | 79 | 3.96 | 6.38E-06 |
| Regulation of Apoptosis (R-HSA-169911) | 51 | 4.43 | 5.07E-05 |
| Regulation of BACH1 activity (R-HSA-9708530) | 10 | 6.95 | 1.06E-02 |
| Regulation of expression of SLITs and ROBOs (R-HSA-9010553) | 168 | 5.48 | 2.75E-23 |
| Regulation of mitotic cell cycle (R-HSA-453276) | 86 | 3.64 | 2.01E-05 |
| Regulation of mRNA stability by proteins that bind AU-rich elements (R-HSA-450531) | 84 | 4.14 | 9.15E-07 |
| Regulation of necroptotic cell death (R-HSA-5675482) | 30 | 3.48 | 3.41E-02 |
| Regulation of ornithine decarboxylase (ODC) (R-HSA-350562) | 50 | 4.87 | 7.93E-06 |
| Regulation of PLK1 Activity at G2/M Transition (R-HSA-2565942) | 83 | 2.72 | 5.91E-03 |
| Regulation of PTEN gene transcription (R-HSA-8943724) | 59 | 3.53 | 1.01E-03 |
| Regulation of PTEN stability and activity (R-HSA-8948751) | 66 | 4.48 | 2.31E-06 |
| Regulation of RAS by GAPs (R-HSA-5658442) | 65 | 4.28 | 8.53E-06 |
| Regulation of TP53 Activity (R-HSA-5633007) | 156 | 2.23 | 5.07E-03 |
| Resolution of Sister Chromatid Cohesion (R-HSA-2500257) | 120 | 2.03 | 4.97E-02 |
| Respiratory electron transport (R-HSA-611105) | 92 | 2.64 | 5.16E-03 |
| Response of EIF2AK4 (GCN2) to amino acid deficiency (R-HSA-9633012) | 101 | 6.54 | 1.02E-19 |
| Response of Mtb to phagocytosis (R-HSA-9637690) | 22 | 5.53 | 1.41E-03 |
| Response to elevated platelet cytosolic Ca2+ (R-HSA-76005) | 130 | 2.54 | 1.40E-03 |
| Retrograde neurotrophin signalling (R-HSA-177504) | 14 | 4.97 | 3.57E-02 |
| RHO GTPase Effectors (R-HSA-195258) | 276 | 2.77 | 2.22E-08 |
| RHOBTB GTPase Cycle (R-HSA-9706574) | 34 | 5.62 | 2.36E-05 |
| Ribosomal scanning and start codon recognition (R-HSA-72702) | 59 | 6.78 | 2.46E-12 |
| RIPK1-mediated regulated necrosis (R-HSA-5213460) | 30 | 3.48 | 3.42E-02 |
| RMTs methylate histone arginines (R-HSA-3214858) | 47 | 4.07 | 5.65E-04 |
| RNA Polymerase I Transcription (R-HSA-73864) | 73 | 3.33 | 6.16E-04 |
| RNA Polymerase II Transcription (R-HSA-73857) | 1306 | 2.05 | 2.61E-16 |
| Role of phospholipids in phagocytosis (R-HSA-2029485) | 28 | 3.72 | 2.49E-02 |
| rRNA processing (R-HSA-72312) | 201 | 4.32 | 3.22E-17 |
| rRNA processing in the nucleus and cytosol (R-HSA-8868773) | 191 | 4.55 | 3.24E-18 |
| S Phase (R-HSA-69242) | 161 | 3.13 | 8.38E-07 |
| SCF(Skp2)-mediated degradation of p27/p21 (R-HSA-187577) | 58 | 5.39 | 5.72E-08 |
| SCF-beta-TrCP mediated degradation of Emi1 (R-HSA-174113) | 53 | 4.92 | 3.16E-06 |
| Selective autophagy (R-HSA-9663891) | 75 | 3.71 | 5.14E-05 |
| Selenoamino acid metabolism (R-HSA-2408522) | 116 | 5.84 | 2.79E-18 |
| Selenocysteine synthesis (R-HSA-2408557) | 93 | 7.1 | 3.89E-21 |
| Senescence-Associated Secretory Phenotype (SASP) (R-HSA-2559582) | 76 | 2.97 | 2.85E-03 |
| Separation of Sister Chromatids (R-HSA-2467813) | 183 | 2.75 | 9.41E-06 |
| SHOC2 M1731 mutant abolishes MRAS complex function (R-HSA-9726840) | 8 | 8.69 | 4.62E-03 |
| Signal transduction by L1 (R-HSA-445144) | 19 | 4.57 | 2.10E-02 |
| Signaling by ALK (R-HSA-201556) | 28 | 3.72 | 2.50E-02 |
| Signaling by ALK in cancer (R-HSA-9700206) | 88 | 3.95 | 1.83E-06 |
| Signaling by BRAF and RAF1 fusions (R-HSA-6802952) | 62 | 3.36 | 1.55E-03 |
| Signaling by EGFR in Cancer (R-HSA-1643713) | 22 | 4.74 | 7.65E-03 |
| Signaling by EGFRvIII in Cancer (R-HSA-5637812) | 13 | 6.68 | 4.12E-03 |
| Signaling by ERBB2 in Cancer (R-HSA-1227990) | 22 | 4.74 | 7.63E-03 |
| Signaling by FGFR (R-HSA-190236) | 76 | 2.74 | 7.92E-03 |
| Signaling by FGFR2 (R-HSA-5654738) | 63 | 3.31 | 1.78E-03 |
| Signaling by FGFR3 (R-HSA-5654741) | 31 | 3.36 | 3.96E-02 |
| Signaling by Hedgehog (R-HSA-5358351) | 145 | 3.6 | 2.23E-08 |
| Signaling by Interleukins (R-HSA-449147) | 452 | 2.23 | 2.18E-07 |
| Signaling by Ligand-Responsive EGFR Variants in Cancer (R-HSA-5637815) | 16 | 6.52 | 1.48E-03 |
| Signaling by MET (R-HSA-6806834) | 75 | 2.55 | 2.06E-02 |
| Signaling by MRAS-complex mutants (R-HSA-9660537) | 8 | 8.69 | 4.65E-03 |
| Signaling by NOTCH (R-HSA-157118) | 196 | 3.64 | 1.85E-11 |
| Signaling by NOTCH2 (R-HSA-1980145) | 32 | 3.26 | 4.55E-02 |
| Signaling by NOTCH3 (R-HSA-9012852) | 47 | 2.96 | 2.70E-02 |
| Signaling by NOTCH4 (R-HSA-9013694) | 79 | 4.84 | 1.11E-08 |
| Signaling by NTRK1 (TRKA) (R-HSA-187037) | 112 | 2.79 | 6.65E-04 |
| Signaling by NTRKs (R-HSA-166520) | 131 | 2.52 | 1.51E-03 |
| Signaling by Nuclear Receptors (R-HSA-9006931) | 255 | 3.2 | 4.43E-11 |
| Signaling by Receptor Tyrosine Kinases (R-HSA-9006934) | 512 | 2.21 | 5.12E-08 |
| Signaling by Rho GTPases (R-HSA-194315) | 656 | 2.28 | 2.72E-11 |
| Signaling by ROBO receptors (R-HSA-376176) | 213 | 4.9 | 1.69E-23 |
| Signaling by TGFB family members (R-HSA-9006936) | 121 | 3.16 | 1.87E-05 |
| Signaling by TGF-beta Receptor Complex (R-HSA-170834) | 91 | 3.63 | 1.15E-05 |
| Signaling by the B Cell Receptor (BCR) (R-HSA-983705) | 108 | 3.54 | 3.15E-06 |
| Signaling by VEGF (R-HSA-194138) | 102 | 2.9 | 6.60E-04 |
| Signaling by WNT (R-HSA-195721) | 291 | 3.28 | 2.18E-13 |
| SMAD2/SMAD3:SMAD4 heterotrimer regulates transcription (R-HSA-2173796) | 35 | 3.97 | 4.91E-03 |
| Somitogenesis (R-HSA-9824272) | 54 | 4.18 | 9.66E-05 |
| Spry regulation of FGF signaling (R-HSA-1295596) | 13 | 5.35 | 2.77E-02 |
| SRP-dependent cotranslational protein targeting to membrane (R-HSA-1799339) | 111 | 6.73 | 7.81E-23 |
| Stabilization of p53 (R-HSA-69541) | 54 | 4.51 | 2.00E-05 |
| SUMO E3 ligases SUMOylate target proteins (R-HSA-3108232) | 157 | 2.44 | 8.98E-04 |
| SUMOylation (R-HSA-2990846) | 163 | 2.35 | 1.87E-03 |
| SUMOylation of transcription cofactors (R-HSA-3899300) | 42 | 3.31 | 1.42E-02 |
| Suppression of apoptosis (R-HSA-9635465) | 7 | 7.45 | 3.01E-02 |
| Suppression of phagosomal maturation (R-HSA-9637687) | 12 | 5.79 | 2.10E-02 |
| Switching of origins to a post-replicative state (R-HSA-69052) | 90 | 4.06 | 6.44E-07 |
| Syndecan interactions (R-HSA-3000170) | 27 | 4.51 | 4.76E-03 |
| Synthesis of active ubiquitin: roles of E1 and E2 enzymes (R-HSA-8866652) | 29 | 5.99 | 3.45E-05 |
| Synthesis of DNA (R-HSA-69239) | 119 | 3.07 | 4.64E-05 |
| TCF dependent signaling in response to WNT (R-HSA-201681) | 195 | 3.48 | 2.45E-10 |
| TCR signaling (R-HSA-202403) | 114 | 3.81 | 1.45E-07 |
| TGF-beta receptor signaling activates SMADs (R-HSA-2173789) | 45 | 3.09 | 2.11E-02 |
| The citric acid (TCA) cycle and respiratory electron transport (R-HSA-1428517) | 164 | 2.12 | 1.12E-02 |
| The role of GTSE1 in G2/M progression after G2 checkpoint (R-HSA-8852276) | 74 | 3.99 | 1.05E-05 |
| TNFR2 non-canonical NF-kB pathway (R-HSA-5668541) | 97 | 2.69 | 3.22E-03 |
| Toll Like Receptor 10 (TLR10) Cascade (R-HSA-168142) | 94 | 2.59 | 6.24E-03 |
| Toll Like Receptor 2 (TLR2) Cascade (R-HSA-181438) | 113 | 2.15 | 3.45E-02 |
| Toll Like Receptor 3 (TLR3) Cascade (R-HSA-168164) | 101 | 2.58 | 4.67E-03 |
| Toll Like Receptor 5 (TLR5) Cascade (R-HSA-168176) | 94 | 2.59 | 6.26E-03 |
| Toll Like Receptor 7/8 (TLR7/8) Cascade (R-HSA-168181) | 101 | 2.41 | 1.15E-02 |
| Toll Like Receptor 9 (TLR9) Cascade (R-HSA-168138) | 104 | 2.51 | 5.94E-03 |
| TP53 Regulates Metabolic Genes (R-HSA-5628897) | 81 | 3.65 | 3.49E-05 |
| TP53 Regulates Transcription of Cell Cycle Genes (R-HSA-6791312) | 49 | 5.32 | 1.15E-06 |
| TRAF6 mediated induction of NFkB and MAP kinases upon TLR7/8 or 9 activation (R-HSA-975138) | 99 | 2.46 | 9.71E-03 |
| Transcription of the HIV genome (R-HSA-167172) | 69 | 2.77 | 1.12E-02 |
| Transcriptional activity of SMAD2/SMAD3:SMAD4 heterotrimer (R-HSA-2173793) | 49 | 4.26 | 1.70E-04 |
| Transcriptional regulation by RUNX1 (R-HSA-8878171) | 197 | 3.53 | 8.43E-11 |
| Transcriptional regulation by RUNX2 (R-HSA-8878166) | 116 | 4.2 | 2.12E-09 |
| Transcriptional regulation by RUNX3 (R-HSA-8878159) | 92 | 4.35 | 4.34E-08 |
| Transcriptional regulation by small RNAs (R-HSA-5578749) | 68 | 3.07 | 3.48E-03 |
| Transcriptional regulation by the AP-2 (TFAP2) family of transcription factors (R-HSA-8864260) | 35 | 3.97 | 4.92E-03 |
| Transcriptional Regulation by TP53 (R-HSA-3700989) | 351 | 2.72 | 3.88E-10 |
| Transcriptional regulation of granulopoiesis (R-HSA-9616222) | 54 | 4.51 | 2.01E-05 |
| trans-Golgi Network Vesicle Budding (R-HSA-199992) | 71 | 3.18 | 1.53E-03 |
| Translation (R-HSA-72766) | 288 | 3.74 | 5.42E-18 |
| Translation initiation complex formation (R-HSA-72649) | 59 | 6.48 | 2.26E-11 |
| Translation of Structural Proteins (R-HSA-9694635) | 56 | 3.1 | 7.64E-03 |
| Transport of Mature mRNA derived from an Intron-Containing Transcript (R-HSA-159236) | 70 | 2.48 | 3.43E-02 |
| Transport of Mature Transcript to Cytoplasm (R-HSA-72202) | 79 | 2.86 | 4.07E-03 |
| Type I hemidesmosome assembly (R-HSA-446107) | 11 | 6.32 | 1.53E-02 |
| Ub-specific processing proteases (R-HSA-5689880) | 187 | 2.42 | 4.27E-04 |
| UCH proteinases (R-HSA-5689603) | 92 | 3.21 | 1.83E-04 |
| Unfolded Protein Response (UPR) (R-HSA-381119) | 91 | 2.29 | 3.13E-02 |
| Uptake and function of diphtheria toxin (R-HSA-5336415) | 5 | 10.43 | 1.07E-02 |
| VEGFA-VEGFR2 Pathway (R-HSA-4420097) | 92 | 3.02 | 6.49E-04 |
| Vif-mediated degradation of APOBEC3G (R-HSA-180585) | 51 | 5.11 | 1.91E-06 |
| Viral Infection Pathways (R-HSA-9824446) | 769 | 3.41 | 1.34E-39 |
| Viral mRNA Translation (R-HSA-192823) | 89 | 7.23 | 6.21E-21 |
| VLDLR internalisation and degradation (R-HSA-8866427) | 15 | 6.95 | 1.03E-03 |
| Vpu mediated degradation of CD4 (R-HSA-180534) | 50 | 4.87 | 7.98E-06 |
| WNT mediated activation of DVL (R-HSA-201688) | 7 | 7.45 | 3.01E-02 |
| WNT5A-dependent internalization of FZD4 (R-HSA-5099900) | 15 | 6.95 | 1.03E-03 |
| Z-decay: degradation of maternal mRNAs by zygotically expressed factors (R-HSA-9820865) | 12 | 10.14 | 1.73E-05 |
